# Extrapolating Heterogeneous Time-Series Gene Expression Data using Sagittarius

**DOI:** 10.1101/2022.12.24.521845

**Authors:** Addie Woicik, Mingxin Zhang, Janelle Chan, Jianzhu Ma, Sheng Wang

**Affiliations:** Paul G. Allen School of Computer Science and Engineering, University of Washington, Seattle, WA; Department of Information and Communications Engineering, Tokyo Institute of Technology, Japan; Department of Electrical Engineering Tsinghua University, China; Institute for AI Industry Research, Tsinghua University, China

## Abstract

Understanding the temporal dynamics of gene expression is crucial for developmental biology, tumor biology, and biogerontology. However, some timepoints remain challenging to measure in the lab, particularly during very early or very late stages of a biological process. Here we propose Sagittarius, a transformer-based model that can accurately simulate gene expression profiles at timepoints outside of the range of times measured in the lab. The key idea behind Sagittarius is to learn a shared reference space for time series measurements, thereby explicitly modeling unaligned timepoints and conditional batch effects between time series, and making the model widely applicable to diverse biological settings. We show Sagittarius’s promising performance when extrapolating mammalian developmental gene expression, simulating drug-induced expression at unmeasured dose and treatment times, and augmenting datasets to accurately predict drug sensitivity. We also used Sagittarius to extrapolate mutation profiles for early-stage cancer patients, which enabled us to discover a gene set connected to the Hedgehog signaling pathway that may be related to tumorigenesis in sarcoma patients, including *PTCH1*, *ARID2*, and *MYCBP2*. By augmenting experimental temporal datasets with crucial but difficult-to-measure extrapolated datapoints, Sagittarius enables deeper insights into the temporal dynamics of heterogeneous transcriptomic processes and can be broadly applied to biological time series extrapolation.

## Main

The temporal dynamics of the transcriptome are key to the study of developmental biology,^1, 2^ tumor biology,^3, 4^ immunobiology,^5, 6^ and pharmacogenomics.^7, 8^ As bulk- and single-cell RNA-sequencing technologies have become cheaper,^4, 9–11^ more transcriptomic datasets include gene expression measurements at multiple timepoints.^12–19^ Still, it often remains challenging to measure transcriptomic profiles at very early or late stages of a biological process. For instance, senescent and extremely diseased tissue can be challenging to measure, but are of extreme interest for aging and therapeutics.

The underlying problem here is temporal extrapolation, where timepoints of interest are outside the range of time with experimental measurements. Accurate extrapolation on a single time series is challenging due to non-stationary features and temporal out-of-domain adaptation.^20^ One possible solution for the extrapolation problem is to combine sparse time series measurements from heterogeneous sequences. For example, mouse^12^ and roundworm^21^ transcriptomic time series measurements, combined with developmental human measurements, can help simulate early-stage embryonic transcriptomic profiles for human.^22^ There are two major challenges in effectively utilizing other time series: unaligned measured timepoints and batch effects between experimental conditions. Existing methods are unable to simultaneously consider the full sequence of measured timepoints^23, 24^ or take into account the temporal batch effects between time series.^25–28^

To address these limitations we propose Sagittarius, a model that maps heterogeneous gene expression time series to a shared reference space based on inferred biological age rather than the observed age, enabling multiple sparsely measured time series to jointly inform extrapolation. Sagittarius leverages a transformer-based architecture with multi-head attention^29^ to map the heterogeneous measurements from the irregular, unaligned, and sparse time series to a latent reference space shared by all time series, using high-frequency sinusoidal embeddings of the timestamp^27, 30^ and experimental condition labels of each time series to define the mapping. After alignment in the shared reference space, Sagittarius can accurately predict new genomic profiles at extrapolated timepoints, as well as predict measurements for unmeasured combinations of experimental conditions.

We evaluated Sagittarius in three diverse settings in developmental biology, pharmacogenomics, and cancer genomics. On the Evo-devo development dataset,^12^ Sagittarius accurately extrapolated gene expression profiles with a 0.983 Pearson correlation, enabling an in-depth analysis of mouse organ differentiation. To evaluate Sagittarius’s robustness to extremely sparse measurements, we next applied it to the LINCS pharmacogenomics dataset^15^ and found that Sagittarius was able to predict drug repurposing opportunities across drugs and cell lines. Finally, we applied Sagittarius to The Cancer Genome Atlas (TCGA) dataset,^31^ where Sagittarius was able to accurately extrapolate mutation profiles for patients with a long survival time. Our findings implicated a gene set related to the Hedgehog signaling pathway and *GLI* oncogene that can potentially drive tumorigenesis in sarcoma patients.

## Results

### Overview of Sagittarius

Given a heterogeneous, unaligned, sparse, and irregular genomic time series dataset, Sagittarius is able to extrapolate gene expression profiles for unmeasured timepoints (**Fig. 1**). Fundamentally, we hypothesize that the input time series follow a common latent trajectory, such as a general developmental trajectory shared by humans and model organisms. The key idea behind Sagittarius is to learn a shared reference space that models this trajectory. We use a transformer-based architecture to map each input measurement to and from the reference space, where the learnable mapping is informed by the experimental conditions associated with the time series. This parameterization addresses both temporal extrapolation and batch effects between experimental conditions (**Methods**). During inference, Sagittarius can extrapolate gene expression profiles for a timepoint and experimental condition of interest.

**Fig. 1.**
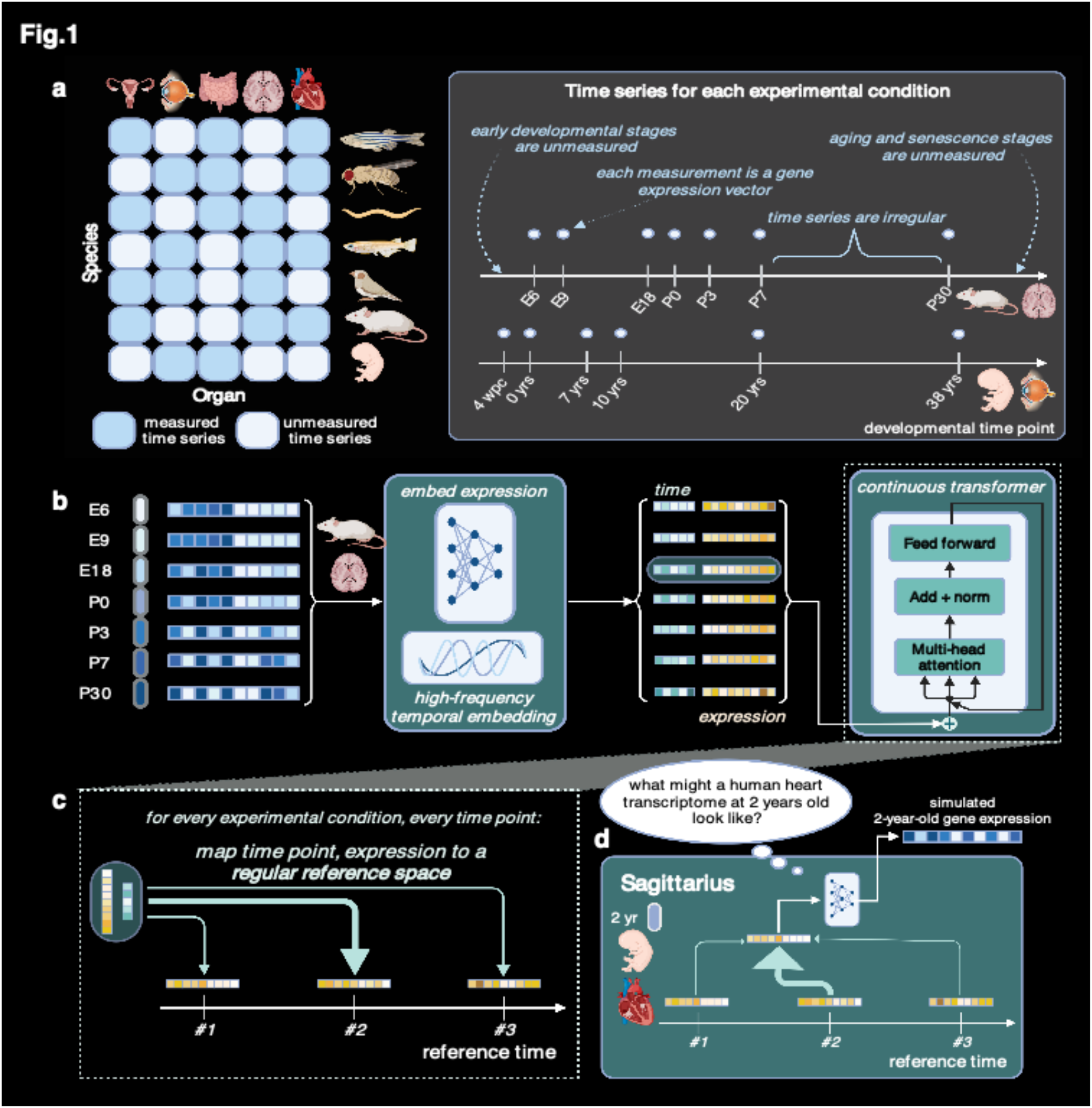
Sagittarius model overview. **a**, Sagittarius is useful in settings with many diverse time series measurements, such as developmental gene expression data across species and organs, many combinations of which are unmeasured. The measurements in each time series are also sparse and unaligned. **b**, For each time series, Sagittarius computes a conditional high-frequency sinusoidal embedding of the measured timepoints and a conditional embedding of the gene expression measurements at each timepoint based on the species and organ. It then uses a continuous, multi-head attention transformer to map the embedded timepoints and expression vectors to the reference space. **c**, The continuous transformer takes each pair of species- and organ-conditioned time and expression embeddings and learns a mapping to the regular reference space, translating from measured age to a shared biological age. **d**, Users can request extrapolated expression vectors from Sagittarius, such as the expression profile of a human 2-year-old heart that has not been measured in the original dataset (**a**). Sagittarius maps the request from the regular reference space back to the data space to predict the unmeasured profile.

### Extrapolating gene expression to unmeasured timepoints

To assess the merit of our approach, we evaluated whether Sagittarius can extrapolate profiles for gene expression time series from multiple experimental conditions. We used the Evo-devo time series data, which contains bulk RNA-seq data from 7 species and 7 organs, where each time series ranges between 9 and 23 distinct measured timepoints that are not biologically aligned across species. Importantly, the developmental ranges measured by each species differ: primates include senescence measurements, while rhesus macaque and chicken do not contain early embryonic data. Therefore, the Evo-devo dataset can be used to assess whether Sagittarius can handle unaligned timepoints and differing biological ages measured across species.

To initially validate our model, we hid the last four measured timepoints from each species’ organ time series to use as a test set. After training on the remaining Evo-devo data, we predicted the gene expression vectors for each species and organ combination at the four hidden timepoints and compared them to the held-out expression vectors. To benchmark Sagittarius’s performance, we also evaluated the deep learning methods Conditional Variational Autoencoder^23^ (cVAE), Compositional Perturbation Autoencoder^24^ (CPA), Multi-Time Attention Network^27^ (mTAN), PRESCIENT,^32^ Neural ODE,^26, 28^ and Recurrent Neural Network^25^ (RNN), as well as classical mean and linear methods. Overall, Sagittarius achieved the best average performance between the extrapolated and measured gene expression profiles in terms of Pearson correlation comparing genes (ρ = 0.983), Pearson correlation comparing timepoints (ρ = 0.458), and root mean squared error (RMSE = 0.087), compared to 0.926, 0.142, and 0.163 respectively for the best-performing comparison approach (**Methods**, **Supplementary Fig. 4,5**), and this improvement was robust to many hyperparameter settings (**Supplementary Note 1, Supplementary Fig. 6-8**). Sagittarius was also able to accurately extrapolate expression for many genes, with at least a 0.3 test Pearson correlation comparing timepoints for more than 55% of the modeled genes (**Methods**).

We further stratified our extrapolation results by species and organ (**Figure 2**). We found that our model achieved the best performance on all species and organs, with best absolute performance on the mouse testis time series. This demonstrates the benefit of the shared reference space, as mouse’s final training timepoint is postnatal day 0 (P0) but the model is able to learn from later development in other species to inform extrapolation for mouse. In contrast, the two worst-performing species were human and chicken, which we believe to reflect larger distributional shifts in the data. All methods struggled on human test data, which are at much later developmental stages than the training dataset. We therefore conducted an analogous Evo-devo experiment, this time extrapolating to the earliest four timepoints as test data (**Methods**). We found that Sagittarius was still the best-performing method, and had stronger performance for extrapolation to early-stage human development (**Supplementary Fig. 9,10**), supporting this hypothesis. We believe that the relatively poor chicken performance also stems from a distributional shift, as chicken is the only non-mammal in the dataset. Therefore, the chicken time series are less evolutionarily related to the other time series in the dataset, and may be harder to represent as a transformation of a shared, otherwise mammalian developmental trajectory.^12^ After better understanding Sagittarius’s strengths and weaknesses, we then studied whether Sagittarius could simulate samples for unmeasured timepoints to gain new insights into tissue differentiation and aging.

**Fig. 2.**
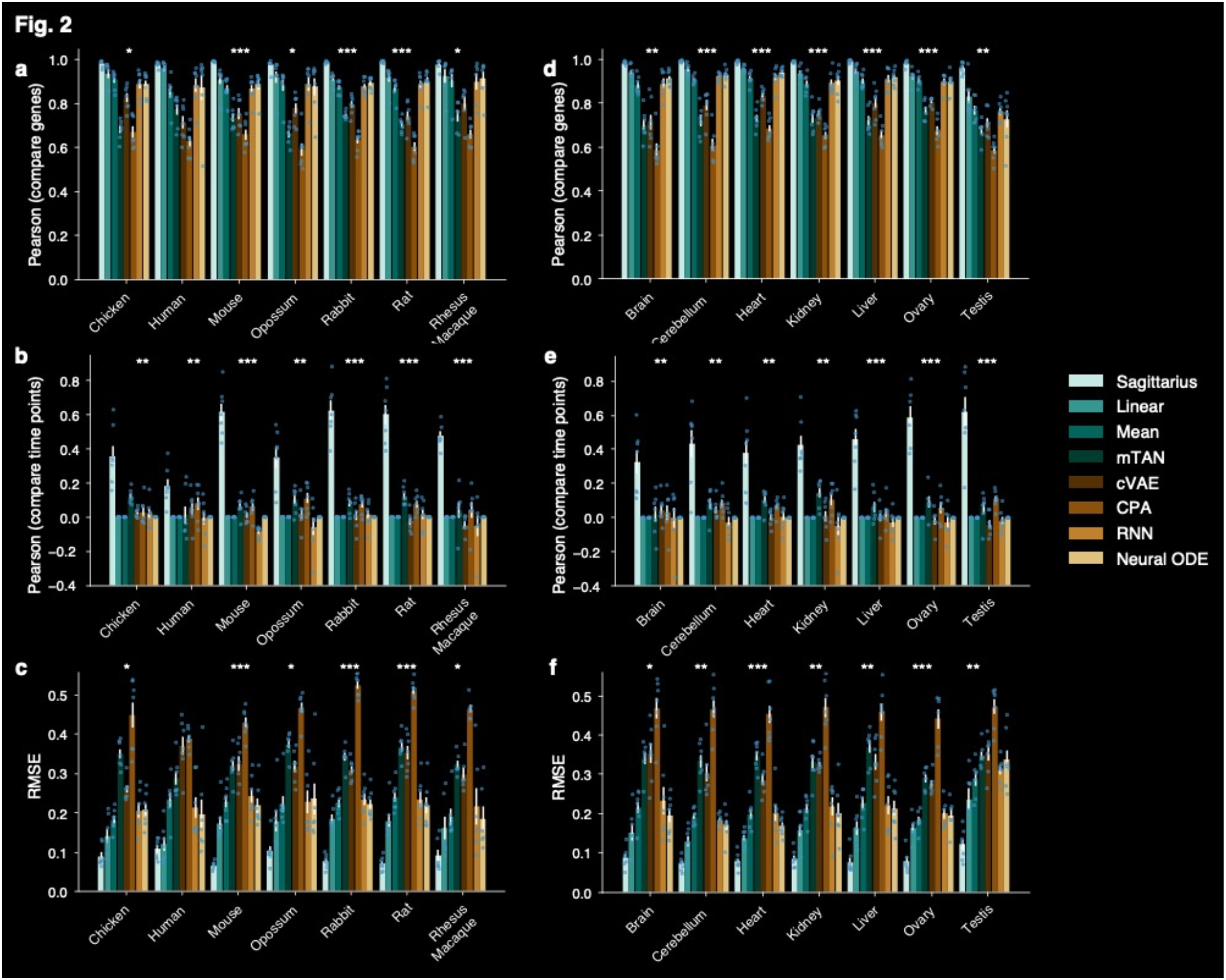
Gene expression prediction for extrapolated timepoints in later-stage development. **a-f**, Bar plots comparing the performance of Sagittarius and existing approaches when extrapolating to the four latest timepoints in the Evo-devo dataset. Test sequences are subdivided by species (**a-c**) and by organ (**d-f**). For Pearson correlation, comparing genes (**a,d**) or comparing timepoints (**b,e**), higher correlations indicate better performance; for RMSE (**c,f**), lower error indicates better performance. Data are presented as mean values +/- standard error. When stratified by species (**a-c**), n=6 organ time series for Rhesus Macaque and n=7 organ time series for all other species. When stratified by organ (**d-f**), n=6 species time series for ovary and n=7 species time series for all other organs. The * indicates that Sagittarius outperforms the next-best-performing model in the metric, with significance levels of p-value < 5e-2 for *, p-value < 5e-3 for **, and p-value < 5e-4 for ***. We use a one-sided Fisher z-transformed test for Pearson correlation comparing genes and comparing timepoints, and a one-sided t-test for RMSE.

### Transcriptomic dynamics reveal organ-specific aging genes

To further examine the Sagittarius’s extrapolated expression profiles, we next predicted developmental trajectories for each mouse organ, beginning at embryonic day 5.5 (E5.5) and continuing to P63. By extrapolating to early timepoints, we expect to observe a hypothetical trajectory that includes organogenesis, which takes place between E6.5 and E8.5 in mouse development.^33, 34^ Specifically, we expect that the earliest extrapolated timepoints result in very similar expression profiles across the different queried organs, which would not have differentiated at this stage. In subsequent days, we would then expect the organs to diverge according to germ layer, before finally separating by organ.^12, 13, 33, 35, 36^ We visualized the uniform manifold approximation and projection^37^ (UMAP) embedding of the simulated organ time series (**Fig. 3a,b**), as well as the top principal components^38^ (**Supplementary Fig. 12**). Our findings largely aligned with the understanding of mouse organogenesis. Namely, the developmental stage dominated the gene extrapolation measurements at the earliest timepoints, with multiple organs grouped in the same location of the UMAP space. At later timepoints, we found that the predicted expression values for brain and cerebellum were more closely grouped together, as were expression measurements for the heart, ovary, and testis, consistent with the ectoderm, mesoderm, and endoderm tissue germ layer classifications.^12^

**Fig. 3.**
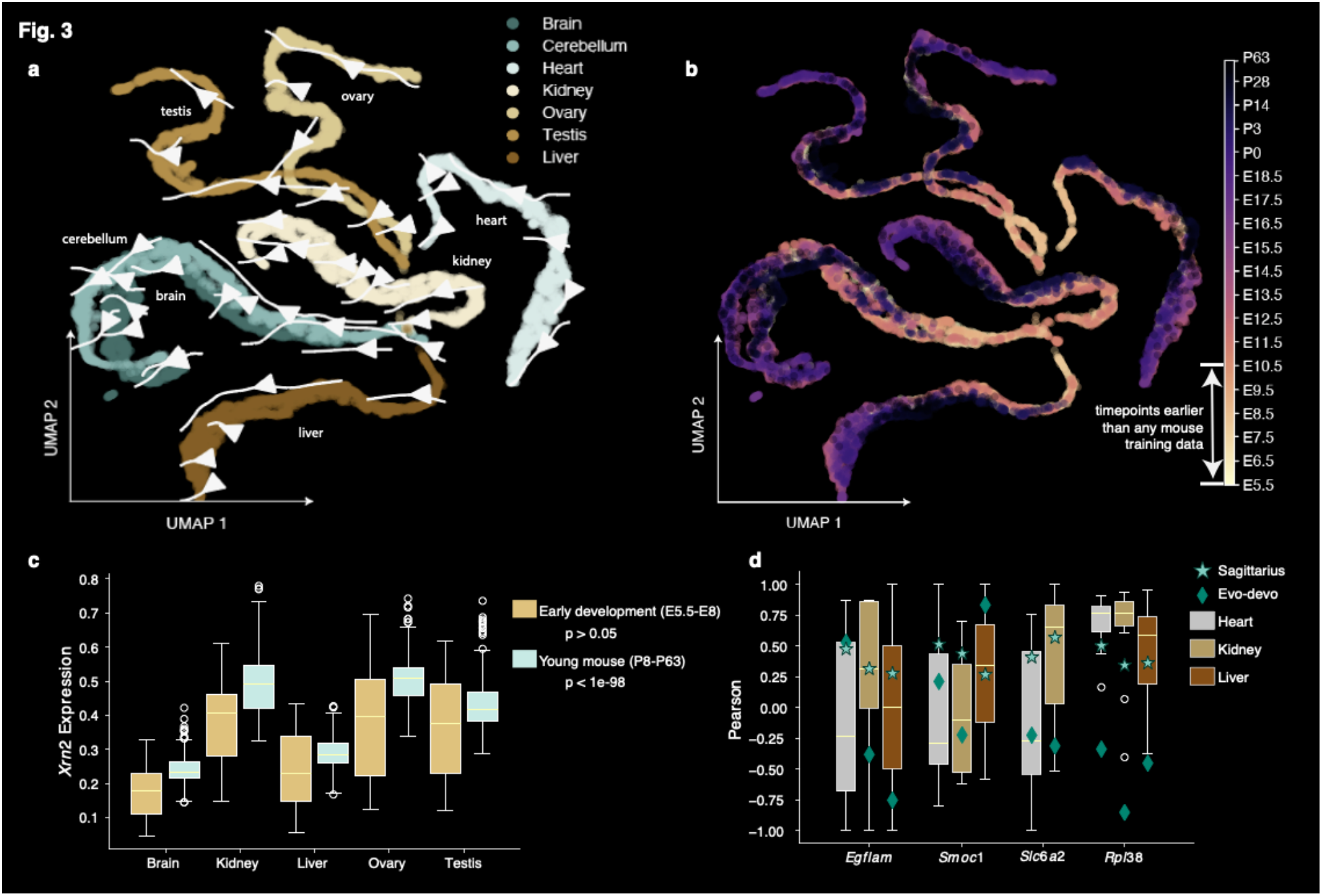
Mouse transcriptomic velocity across organs. **a,b**, UMAP plots showing simulated mouse gene expression from E5.5 to P63 for 7 organs, colored by organ (**a**) and time (**b**). The arrows in (**a**) indicate the transcriptomic velocity of each organ. **c**, Box plot comparing the simulated expression of *Xrn2* at early development (E5.5-E8) to young mouse (P8-P63) across five organs, with n=250 simulated measurements per organ. *Xrn2* expression is not statistically different between the brain, kidney, liver, ovary, and testis organs at the early development (ANOVA p-value = 0.83), but differs between organs at the young mouse time range, particularly with lower expression levels in the liver relative to other organs (ANOVA p-value = 9.37e-78). **d**, Box plot examining the consistency of gene expression temporal patterns between simulated data and scRNA-seq data for *Egflam*, *Smoc1*, *Slc6a2*, and *Rpl38* in different mouse tissues over time. Boxes indicate the distribution of cell type correlations for each tissue in Tabula Muris Senis, with n=7,5,2 cell types in heart, kidney, and liver respectively for *Egflam*; n=7,6,3 respectively for *Smoc1*; n=5,7 in heart and kidney respectively for *Slc6a2*; and n=9,19,8 cell types in heart, kidney, and liver for *Rpl38*. Better predictions are closer to the distribution of Tabula Muris Senis cell type correlations for each tissue. The star shows the Pearson correlation from Sagittarius’s simulated correlation for aging mouse tissues (140 timepoints beginning at P14), and the diamond shows the correlation with respect to time of the younger mouse organs measurements in the Evo-devo dataset. In both **c,d**, the box bounds show the interquartile range (IQR) from quartiles 1 to 3 (Q1-Q3), with the centerline indicating the median and the whiskers extending 1.5 IQR from the box.

Given the increasing tissue-specific signal in Sagittarius’s simulated gene expression vectors at later timepoints, we then investigated which genes most contributed to the differentiation of organ trajectories during development. Excluding the heart and cerebellum, which we found to be the most developmentally distinct for many genes, we found that mouse *Xrn2* expression levels were comparable across organs at early extrapolated timepoints but differed significantly at later timepoints (ANOVA p-value > 0.05 and p-value < 1e-98 respectively), with lower expression levels in the liver than other organs at late timepoints (**Fig. 3c**). Existing work has found that mouse *Xrn2* and its roundworm orthologue *xrn-2* play important biological roles during development,^39–46^ and *XRN2* overexpression in human has also been tied to poor liver cancer prognosis.^47^

We next sought to further examine Sagittarius’s organ-specific extrapolation potential in mouse. We predicted a transcriptomic profile trajectory beginning at P14 (**Methods**). The latest mouse measurement in the Evo-devo dataset is taken at P63, so we used the Tabula Muris Senis single cell RNA-seq dataset,^16^ which spans from a 1-month-old mouse to a 30-month-old mouse, to validate our results. We compared the Pearson correlation of the gene expression over time between the extrapolated profiles and the Tabula Muris Senis data for each tissue, and for mouse genes including *Egflam*, *Smoc1*, *Slc6a2*, and especially *Rpl38*, which previous work has suggested could regulate developmental processes in a tissue-specific way,^48^ found that Sagittarius’s extrapolated aging trajectory aligned with the Tabula Muris Senis tissue measurements better than younger mouse trajectory taken directly from Evo-devo (**Methods**, **Fig. 3d**). We attribute this to the shared reference space, which can identify aging and senescence patterns from other species like human and rhesus macaque to inform transcriptomic extrapolation for mouse aging. After applying Sagittarius to the Evo-devo dataset, we next considered whether the model could successfully extrapolate unmeasured experimental combinations in settings with multiple continuous variables.

### Sagittarius simulates unmeasured drug perturbations

We next evaluated Sagittarius on extremely sparse multivariate data with multiple continuous temporal variables, thereby exponentially increasing the space of possible experimental settings. We applied Sagittarius to the larger, high-dimensional LINCS L1000 pharmacogenomics dataset.^15^ In the LINCS dataset, compounds are experimentally applied to cell lines at specific doses and for a given treatment time before measuring the drug-induced expression profiles, although only 1.77% of possible drug and cell line combinations are screened (**Fig. 4a**). Sagittarius models each treatment experiment in two continuous dimensions: dose and treatment time.

**Fig. 4.**
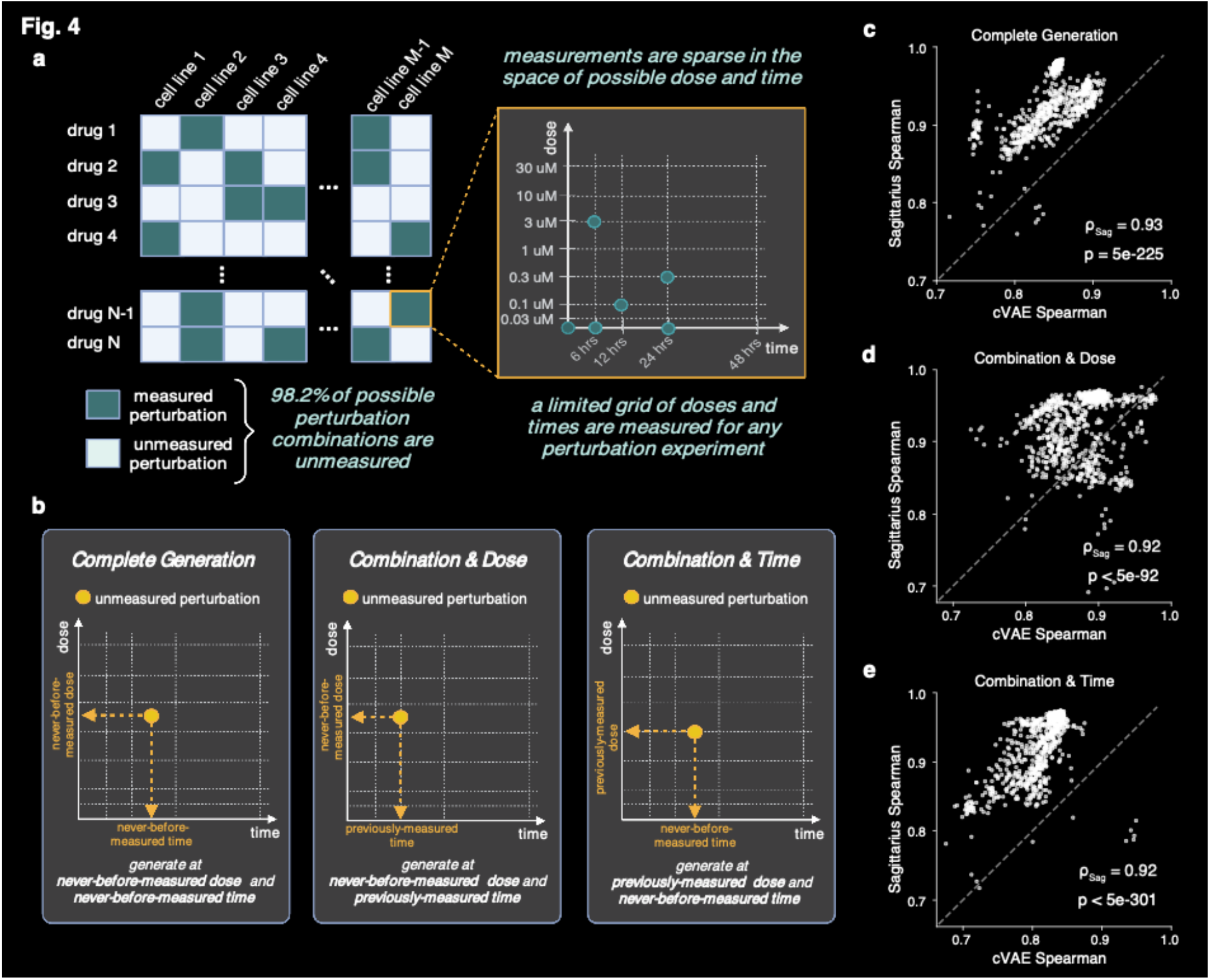
Drug-induced gene expression extrapolation at unmeasured experimental combinations, doses, and times. **a**, The LINCS pharmacogenomic dataset contains gene expression measurements from a set of experiments where a cancer cell line is treated with a therapeutic compound. The set of measured cell lines and compounds is sparse, with less than 1.77% of possible experiments measured. The measured experiments are also only measured at select dose and treatment times, and the entire dataset includes a limited number of dose and treatment times. **b**, Illustration of the three extrapolation tasks we evaluate for the LINCS dataset: complete generation, where we predict an unmeasured cell line and compound experiment at both a dose and time that are unmeasured by any experiment in the dataset; combination & dose, where we predict an unmeasured cell line and compound experiment at a time that has been measured in the dataset but a dose that is unmeasured by all experiments; and combination & time, where we predict an unmeasured cell line and compound experiment at a dose that has been measured in the dataset but a time that is unmeasured by all experiments. **c-e**, Scatter plots comparing the average Spearman correlation of simulated test combinations from Sagittarius and the existing cVAE model for each test drug on the complete generation (**c**), combination & dosage (**d**), and combination & time (**e**) extrapolation tasks. Sagittarius’s average correlation across test datapoints is reported inline, along with the one-sided Fisher z-transformed test p-values testing whether Sagittarius outperforms cVAE in each settings, with p-value = 3.94e-225, p-value = 2.73e-92, and p-value = 2.50e-301 respectively.

To validate Sagittarius’s ability to extrapolate to new perturbation experiments, we then designed three extrapolation tasks: complete generation, combination & dose, and combination & time (**Methods**, **Fig. 4b**). For each task, we trained Sagittarius on a subset of the LINCS dataset, withholding the remaining measurements as test data. We compared Sagittarius’s performance to a cVAE, the only comparison approach that could handle multiple continuous variables off-the-shelf. Evaluating model predictions based on held-out test data, we found that Sagittarius had an average Spearman correlation of 0.93, 0.92, and 0.88 for the three tasks respectively, compared to 0.85, 0.88, and 0.81 for the cVAE (**Fig. 4c-e**, one-sided Fisher z-transformed test p-value < 5e-225, 5e-92, and 5e-301 respectively). This indicates that aligning all perturbations experiments to the shared reference space enables Sagittarius to accurately extrapolate drug-induced gene expression vectors for unmeasured drug treatment experiments at doses and times that are not contained in the training data, enabling an easy, unbiased search approach to drug sensitivity markers. This can greatly increase our understanding of the molecular basis of cancer and of drug response.

### A drug sensitivity similarity network for drug repurposing

Gene expression has been widely used to identify the drug-induced and diseased-induced gene expression signatures in drug repurposing studies,^49–51^ partly due to the scale at which analyses can be efficiently performed and validated. As Sagittarius can accurately predict expression for any perturbation combinations, we applied Sagittarius to drug repurposing by constructing a similarity network of extrapolated perturbation experiments (**Methods**). We found that communities within the network demonstrated a pattern with respect to treatment sensitivity, with average half-maximal inhibitory concentration (IC50) doses of 1.68, 1.83, 1.90, and 2.40 uM in the Genomics of Drug Sensitivity in Cancer (GDSC) dataset^52^ (**Fig. 5a**). To further investigate the potential for drug-repurposing opportunities, we conducted a case study on an 8-experiment subgraph from the sensitive community, shown in the inset of **Fig. 5a**.

**Fig. 5.**
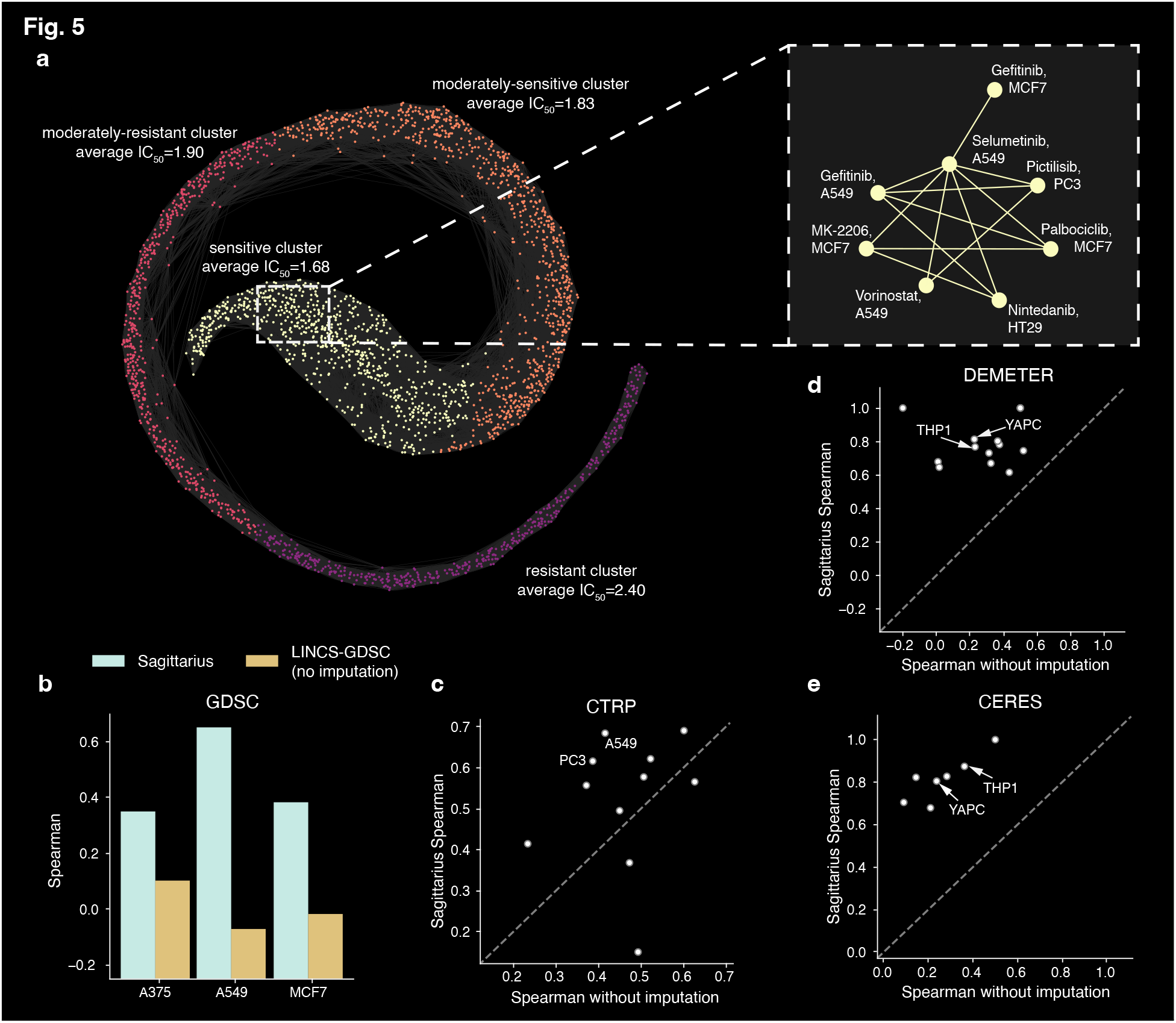
Drug and cell line treatment efficacy extrapolation analysis. **a**, k-nearest neighbor network where each node represents a drug and cell line combination, with edges between the most similar drug-induced expression effect. The four communities in the graph are shown in different colors and labeled according to the average GDSC-measured IC50 dose of that community, measured in uM. The inset shows a connected 8-node subgraph from the sensitive community, made up of the NSCLC cell line A549 treated with Selumetinib, Gefitinib, and Vorniostat; the breast cancer cell line MCF7 treated with Gefitinib, MK-2206, and Palbociclib; the prostate carcinoma cell line PC3 treated with Pictilisib; and the colorectal adenocarcinoma cell line HT29, treated with Ninetedanib. **b,c**, Bar plot (**b**) and scatter plot (**c**) of Spearman correlation between predicted and GDSC-measured (**b**) or CTRP-measured (**c**) IC50 doses per cell line, comparing a neural network trained with imputed data from Sagittarius and a neural network trained with only the GDSC or CTRP treatment combinations that are also measured in LINCS. Each cell line has one correlation coefficient, meaning n=1 cell line sensitivity ranking for the bar plot in **b**. Points above the *y* = *x* line are cell lines for which Sagittarius’s imputed dataset improved the downstream prediction accuracy (**c**). **d,e**, Scatter plot of Spearman correlation between predicted and DepMap-measured cancer gene essentiality scores for each cancer line, with the DEMETER (**d**) and CERES (**e**) DepMap versions. A neural network trained with imputed data from Sagittarius is compared to a model trained on the LINCS treatment combinations that correspond to cell line and gene pairs in the DEMETER or CERES datasets. All points are above the *y* = *x* line, meaning Sagittarius improved downstream gene essentiality prediction performance for all cell lines on both DepMap versions.

The subnetwork includes six perturbations for the breast carcinoma cell line MCF7 and non-small cell lung cancer cell line A549, all of which are measured in the GDSC dataset. A549 is sensitive to treatment with Vorinostat, Gefitinib, and Selumetinib (IC50 of 0.49, 0.67, 0.83 uM respectively), and MCF7 is sensitive to treatment with Palbociclib, MK-2206, and Gefitinib (IC50 of 0.40, 0.89, and 1.03 uM respectively). The existence of edges between different drugs for the same cell line and the edge between two cell lines for the same drug connect to drug-repurposing work based on cell line gene expression signatures^53^ and drug mechanisms of action.^54^

Importantly, by comparing extrapolated differential expression signatures to those of successful treatments, Sagittarius enables drug repurposing recommendations where neither the drug nor cell line needs to occur in a known successful therapy. The 8-perturbation subnetwork also includes the prostate adenocarcinoma cell line PC3 treated with Pictilisib and the colorectal adenocarcinoma cell line HT29 treated with Nintedanib, although these drugs and cell lines do not appear elsewhere in the subnetwork. The GDSC dataset does not screen either of these treatment combinations, but previous work has found that Pictilisib inhibited PC3 proliferation (IC50=0.28 uM)^55^ and Nintedanib had significant antitumor efficacy in HT29 xenograft models^56, 57^ and cell lines (IC50=1.40 uM).^57^ This implies that Sagittarius can extrapolate perturbation experiments to identify candidate drug repurposing targets across cell lines, cancer types, and therapeutic compounds, creating new opportunities for inexpensive and unbiased drug screening as an initial step in the precision medicine pipeline.

### Perturbation augmentation improves drug response prediction

Given its drug repurposing potential, we systematically evaluated Sagittarius on two large-scale cell line drug response prediction datasets, GDSC and the Cancer Therapeutic Response Portal (CTRP) dataset.^58^ Drug-induced expression profiles have been useful for drug response prediction,^59^ but are expensive to measure compared to basal cell line expression, making Sagittarius’s extrapolated data especially valuable. We used a downstream neural network to predict the IC50 label for treatment perturbations, and compared the performance of a model trained on extrapolated data from Sagittarius to a model trained on the available measured perturbations in LINCS (**Methods**). When trained on data from Sagittarius, the downstream model had an average Spearman correlation of 0.46 and 0.52 per test cell line, comparing the predicted drug sensitivities to the measured sensitivities in GDSC and CTRP respectively (**Fig. 5b,c**). In comparison, the model trained on the available experimentally-measured data had an average correlation of 0.004 in GDSC and 0.42 in CTRP. We attribute the poor GDSC performance to the little overlap in treatments screened by the two datasets and, consequently, to the extremely small size of the training dataset. Sagittarius, in contrast, was able to extrapolate missing profiles and improve downstream performance, particularly for cell lines and drugs that were among the most frequently measured in the LINCS dataset (**Supplementary Fig. 14,15**). This shows that Sagittarius can take advantage of the many perturbation experiments to inform better predictions for each drug and cell line, even when applied to unmeasured or sparsely measured combinations.

### Improved cancer-essential gene prediction using Sagittarius

In addition to drug response, we analyzed Sagittarius’s ability to predict cancer gene essentiality. We used the Cancer Dependency Map (DepMap), considering both the DEMETER^60^ and the CERES^61^ versions. We used a downstream neural network to predict gene essentiality in a given cell line, where we featurized each cell line and gene pair using a drug-induced expression vector from the given cell line and a LINCS drug with the target gene of interest (**Methods**). We trained one version of the downstream model on extrapolated data from Sagittarius and another on the available experimentally-measured drug-induced profiles in LINCS. Comparing the predicted gene essentiality scores with the labels in DepMap, the model trained on data from Sagittarius obtained an average test Spearman correlation of 0.789 and 0.816 per cell line for DEMETER and CERES respectively, relative to 0.278 and 0.261 for the model trained solely on the *in vitro* LINCS data (**Fig. 5d,e**). Again, we found that the Sagittarius-backed model particularly improved predictions for well-measured LINCS cell lines including the THP1 leukemia and YAPC pancreatic cell lines (**Supplementary Fig. 16**). We attribute the strong performance across many different cancer types and drug targets to the shared reference space, where dose- and treatment-time response can be compared across cancer cell lines and compounds.

### Simulating mutation profiles for early-stage cancer patients

Having extrapolated gene expression time series in one- and two-continuous dimensions, we then sought to apply Sagittarius to cancer survival time data. We focus on extrapolating somatic mutation profiles as strong signals of disease, but also validate our results on patient gene expression profiles (**Methods**, **Supplementary Fig. 18**). It remains very challenging to measure genomic profiles from patients with nascent tumors, as they are rarely diagnosed at this stage,^62^ and yet these initial mutations can be the most informative as to the cancer’s mechanisms and potential early-intervention therapies before other passenger mutations accumulate.^63^ We therefore designed a time-series formulation for patient data from 24 cancer types in The Cancer Genome Atlas (TCGA) dataset,^31^ where extrapolation to later timepoints corresponds to the mutation profiles of patients with longer survival times (**Fig. 6a**). To account for censored event times, we leverage recent machine learning techniques^64, 65^ to exclude censored patients whose event time likely differs from their survival time (**Methods**). As a result, the sarcoma (SARC) cancer type time series contains 115 patients, 31 of whom had a censored death event (**Fig. 6b**).

**Fig. 6.**
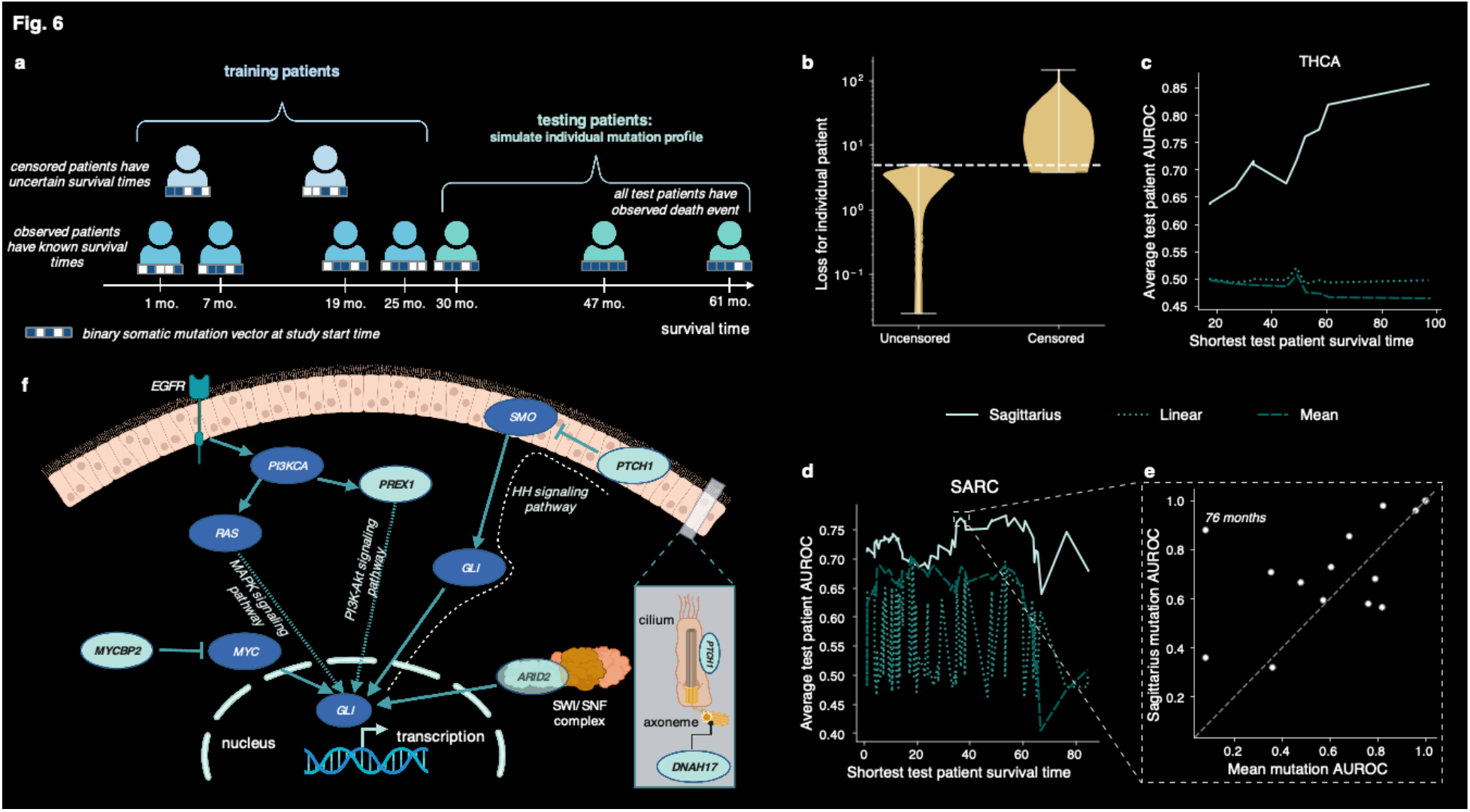
Early cancer patient mutation profile extrapolation. **a**, Illustration of the training and testing splits for a given cancer type in the TCGA extrapolation task, where training patients have the shortest survival times and test patients have longer survival times for that cancer type. **b**, Violin plot of the survival time regression model’s absolute error per patient for the SARC cancer type, divided according to the patient’s censoring label. We remove all patients with a loss above the dashed line from the dataset, and train Sagittarius only on the patients below the dashed line. **c,d**, Plot of the average predicted mutation profile AUROC for each of the THCA (**c**) and SARC (**d**) cancer type test splits, ordered according to the shortest survival time in that test split. **e**, Scatter plot comparing the per-patient predicted mutation profile AUROC from Sagittarius and the mean comparison approach for the SARC test split including patients with an observed death event more than 37 months after diagnosis. Points above the *y* = *x* line indicate that Sagittarius had a better predicted mutation profile than the comparison approach. **f**, Illustration of the ties between the *GLI* oncogene in the Hedgehog (HH) signaling pathway and the *PTCH1*, *PREX1*, *MYCBP2*, *ARID2*, and *DNAH17* genes that Sagittarius predicted as among the most likely mutations in early-stage sarcoma patients.

Focusing on the SARC and thyroid carcinoma (THCA) cancer types as case studies, we designated the longest-surviving patients as test data and used Sagittarius to extrapolate mutation profiles with the same survival times. Varying the number of test patients to examine the models’ performance in different settings, we found that Sagittarius had an average test set AUROC of 0.72 and 0.73 between the extrapolated and measured mutation profiles for THCA and SARC respectively (**Methods**, **Fig. 6c,d**), representing 45% and 11% improvements respectively over classical methods, and also improved over other deep learning methods (**Supplementary Fig. 21**). As a further case study, we focused on a sarcoma patient with a 76-month post-diagnosis survival time, where Sagittarius had particularly improved the test AUROC (**Fig. 6e**). The patient had a mutation in *LRP1B*, but the mean method assigned 0 probability to this mutation, reflecting the observed distribution of sarcoma patients with worse prognosis. Meanwhile, Sagittarius predicts *LRP1B* as the fourth most likely mutation, perhaps learning that *LRP1B* mutations are associated with good therapeutic response in many cancer types.^66^ Sagittarius also assigns higher likelihood to common sarcoma mutations like *ADGRV1*,^67^ suggesting that the model can jointly leverage patterns within the SARC training data and patterns from other cancer types to improve extrapolation.

### Hedgehog signaling pathway in simulated early-stage sarcoma

Having confirmed our ability to extrapolate mutation profiles for sarcoma patients with longer survival times, we retrained Sagittarius on all cancer type time series and then extrapolated gene mutation profiles for 27 early-stage sarcoma patients, resulting in the most-likely mutated gene set *DNAH17, PREX1, EGFLAM, FAM47B, DSEL, ARID2, TRPM1, NLGN1, PTCH1,* and *MYCBP2* (**Methods, Supplementary Fig. 23**). We found that many of these genes are related to the Hedgehog (HH) signaling pathway and improper activation of the *GLI* oncogene (**Fig. 6f**), which has been connected to improved survival outcomes in sarcoma patients.^68, 69^ For instance, *PTCH1* is a tumor suppressor gene^70^ in the HH pathway connected to some sarcomas;^70–72^ *MYCBP2* encodes a protein that has been shown to interact with *GLI*^73^ via *MYC* upregulation;^74, 75^ the protein encoded by *ARID2* directly interacts with *GLI1*;^76, 77^ *DNAH17* encodes a protein that affects the HH pathway through its role in the primary cilia;^78, 79^ *PREX1* encodes a protein whose pathway has been associated with *GLI* code regulation^80^ and cross-talk with the HH pathway in melanoma.^72, 81^ In addition to these connections to the *GLI* oncogene, *EGFLAM* has been shown to induce activation of *PREX1*,^82^ and *NLGN1* encodes a protein that was found to be significantly enriched with the HH pathway in colorectal carcinoma^83^ (**Supplementary Table 3**). Furthermore, previous gene expression analyses found that the HH signaling pathway was enriched for differentially expressed genes in multiple sarcomas.^84, 85^ Sagittarius’s extrapolated mutation profiles therefore point to the HH signaling pathway and particularly the hyperactivation of the *GLI* oncogene as potentially significant sources of tumorigenesis in sarcomas.

## Discussion

Sagittarius enables extrapolation of genomic profiles from sparse, heterogeneous time series without requiring aligned timepoints or batch correction between different experimental conditions. Although Sagittarius can extrapolate to unseen timepoints, the model still struggles with large domain shifts between developmental stages in training and test, as we identify in the Evo-devo human extrapolation task. Similarly, Sagittarius is unable to extrapolate to precise timepoints. The learned mapping to and from the shared reference space enables comparison between diverse and unaligned time series, but also warps the queried and measured timepoints to align with biological age and thereby alters the timepoints outside the range of the dataset in potentially unforeseen ways. In the future, Sagittarius could be integrated with models to predict the chronological and biological age associated with a new sample. Furthermore, Sagittarius could benefit from work in model calibration^86^ to output a model confidence score along with a predicted profile.

Sagittarius is inspired by decades of work in modeling cell dynamics. One key difference between Sagittarius and pseudotime cell fate models like Monocle,^13^ Palantir,^87^ and Slingshot^88^ is that these models reconstruct cell lineage within the bounds of one or more originator states and one or more terminal states, while Sagittarius is able to extrapolate beyond the range of timepoints in the data. Pseudodynamics^89^ later extended cell fate modeling to time-resolved steps, enabling extrapolation, but operates in a low-dimensional cell state space. Most recently, PRESCIENT^32^ enabled single-cell transcriptomic trajectory modeling, but suffers worse performance in bulk expression modeling. Sagittarius strives to build upon these works by leveraging time series from related yet heterogeneous biological time series for accurate high-dimensional profile extrapolation.

## Methods

The Sagittarius model is divided into encoder and decoder modules around the shared reference space (**Supplementary Fig. 1**). For a set of *N* heterogeneous input time gene expression time series 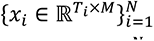 each associated with *C* ≥ 1categorical environmental labels {*y_i_* ∈ 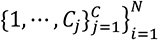 and *B* ≥ 1 continuous variables 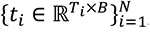, the encoder first embeds each expression measurement, disentangling the environmental conditions from the embedded representation. The encoder also embeds the timepoint of each measurement. First, we use a high-frequency sine wave that both maps each timepoint to the range [0,1], mitigating potential out-of-domain challenges with increasingly large scalar inputs, and helps Sagittarius learn high-frequency patterns in the data, which neural networks have been shown to learn more slowly than low-frequency patterns.^27, 30^ We then learn a full timepoint embedding conditioned on the time series’s environmental conditions, effectively learning a representation of the biological age of each measurement. Finally, we map the embedded expression measurements, along with their inferred biological age embeddings, to the reference space using a transformer encoder. This maps each input or source time series *i* to the common trajectory that the model learns for the heterogeneous dataset (**Supplementary Fig. 2**).

The Sagittarius decoder proceeds in reverse. First, we use the decoder layers of the transformer and learned representations of biological age in each environmental condition to map from the reference space back to embedded measurements for the queried timepoints and condition of interest. Then we decode the embeddings, conditioned on the queried embeddings, to predict the full output expression vectors for each timepoint (**Supplementary Fig. 3**). A detailed discussion of the model is presented in **Supplementary Note 1**.

### Dataset processing

#### Evo-devo

The Evo-devo dataset^12^ contains gene expression vectors for 7 species and 7 organs measured at multiple pre- and post-natal timepoints. We associated each measurement with the environmental variables *y_i_* =[*species_i_*, *organ_i_*]. We restricted the dataset to orthologous genes in all 7 species, identified with the python `pybiomart` package^90^ and the provided Ensembl gene IDs, resulting in 5,037 genes. The measured timepoints for each species were given as strings with different units by species. As a pre-processing step, we computed each timepoint’s rank within the ordered timepoints for that species to use as the timepoint label. We then randomly selected the rabbit heart time series and used the Augmented Dickey-Fuller (ADF) test, which tests for stationary, and only retained genes with p-value > 0.05, resulting in 4,533 genes.

#### LINCS

We used the LINCS L1000 Platform level 3 pharmacogenomic dataset.^15^ We restricted the data experiments to dose measurements in μM that did not exceed 20 μM. We then further restricted the dataset to only include drug and cell line treatment combinations that had more than 15 measurements in the restricted dataset, resulting in 2,687 retained combinations that each had between 16 and 78 measurements (**Supplementary Fig. 13**). We interpreted these as time series in two continuous variables, with *y_i_* = [*drug_i_*, *cell line_i_*], *t_i_*[*dose_i_*, *time_i_*], and *x_i_*[*dose_d_*, *time t*] ∈ ℝ^978^.

#### TCGA

We used the TCGA Firehose legacy dataset,^31^ independently considering the somatic mutation and RNA-seq datasets. Both datasets measure 20,501 genes: we restricted the dataset to the 1,000 most-frequently-mutated genes for the mutation experiments and the 1,000 most-highly-variable genes for the gene expression experiments, jointly considering all cancer types. We removed all patients with missing event times, and excluded mutation patients with no mutations in the 1,000 remaining genes. Finally, we excluded cancer types with fewer than 12 remaining patients. We finally constructed a time series for each cancer type *y_i_*, where the *r*th patient in the time series had event time *t_i_*[*r*] and mutation or gene expression profile *x_i_*[*r*]. Although this formulation enables extrapolation to cancer patients with good prognosis, censored patients may confound the temporal ordering of patients. For the mutation experiments, we exclude all censored patients whose reported event times likely differ dramatically from their survival times, where this probability is learned with a machine learning model (**Supplementary Note 1**).

### Quantitative experiments

For each experiment, we define the test set 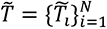, which we use to evaluate model performance (**Supplementary Note 1**).

#### Evo-devo

For the late developmental stage extrapolation experiment, we defined the test set *T̃*_i_ to contain the final four measurements from each time series *i*; analogously, we set *T̃*_i_ to contain the first four measurements from each time series for the early development extrapolation task. In both settings, this resulted in 192 test measurements and 471 measurements available for training or validation. At evaluation time, we used the models to predict the expression vectors at the test timepoints *T̃* and compared the predicted results to the masked measurements in the Evo-devo dataset.

#### LINCS

We defined three main quantitative LINCS experiments: complete generation, combination & dose, and combination & time. For each setting, we first randomly select 5 drug and cell line treatment combinations to mask from the training data, requiring that both the drug and the cell line appear at least once somewhere else in the dataset. We always add all measurements from these five treatments as test points *T̃*. In complete generation, we also randomly select 3 non-zero doses and 1 non-zero treatment time; in combination & dose, we randomly select 3 non-zero doses; in combination & time we select 1 non-zero treatment time. In all settings, any measurement at any selected dose or time are also added to the test set *T̃*. This resulted in 2,144 training sequences and 269 validation sequences for each of the three tasks. There were 7651, 27242, and 10417 total training measurements and 924, 3326, and 1202 validation measurements for the complete generation, combination & dose, and combination & time tasks respectively. Finally, the three tasks had 7441, 7377, and 7395 test sequences with 15068, 14905, and 14966 test measurements respectively.

#### TCGA

For a cancer type *i* with *Γ* observed patients, we defined a training split of the *k* observed patients with shortest survival time, as well as all censored patients with a shorter event time than the *k*th-longest observed survival time. We then used the remaining *Γ* – *k* observed patients as the test patients *T̃_i_*. We further varied k = 1,2,…, *Γ* – 1 to study the effect of different training and test split sizes. In the TCGA gene expression extrapolation experiment, this leads to 13 distinct THCA test splits and 92 SARC test splits. In the TCGA mutation extrapolation experiment, this leads to 9 unique THCA test splits and 61 SARC test splits.

### Evo-devo gene-level predictions

To evaluate Sagittarius’s performance at the level of a single gene, we used the extrapolation to late timepoint experiment and computed the Pearson correlation comparing timepoints for each gene. To identify house-keeping genes^91^ (HKGs), we first stratified the Evo-devo dataset by species and computed the standard deviation of each gene across all organs and timepoints. We then identified 114 HKGs per species using the one-sided rank-sum test with p-value < 0.05. To test whether HKGs and non-HKGs had different performance, we used Fisher’s exact test at a 0.4 correlation threshold (**Supplementary Table 1**). Excluding Gene Ontology^92^ (GO) terms with 1,000 or more associated genes, we then ran a gene set enrichment analysis for the gene set with correlation of at least 0.4 for each time series, retaining the GO terms with p-value < 0.05 after Bonferroni adjustment (**Supplementary Table 2**).

### Mouse developmental analysis

After training Sagittarius on the complete Evo-devo dataset, we used Sagittarius to simulate 10 gene expression time series for each of the 7 mouse organs, using the original mouse organ time series as source sequences for the model. To study early development and young mouse, we extrapolated to timepoint indices ranging from -5 to 13 with granularity 0.1, resulting in 180 predicted measurements per organ per time series. To study aging mouse, we extrapolated to timepoint indices ranging from 11 to 25 with granularity 0.1, resulting in 140 predicted measurements per organ per time series. We smoothed the trajectories by computing the moving average with a window size of 10 (averaging over 5 measurements in each direction).

#### Transcriptomic velocity

We computed the UMAP^37^ embeddings of the smoothed trajectories for early development and young mouse. We also computed the developmental velocity in the UMAP space as the vector between the predicted measurements at time *t* + 0.1 and time *t* in the embedded space for all trajectories and all *t*. We then smoothed the velocity vectors by taking the average of the velocity at time *t* and time *t* – 0.1. We then took the normalized average (mean) over the smoothed velocities associated with each of the 10 simulated trajectories for a given organ. To decrease clutter in the organ development plot, we took the resulting velocity at integer timepoints. We then projected all gene expression measurements to a grid in the UMAP space, and defined the velocity at each point as the weighted average of the 100 velocity vectors nearest to that grid point using the `NearestNeighbors` module from `sklearn.neighbors`.^93^ Finally, we discarded the 5% of velocities with smallest magnitude to simplify the plot. We repeated these steps using a PCA embedding space for the analogous PCA visualization (**Supplementary Fig. 12**).

#### Organ development genes

To identify genes that had a very similar expression at early developmental stages but differing expression levels at later developmental stages, we considered the first and last 25 timepoints from each of the smoothed organ trajectories corresponding to early development and young mouse, resulting in a total of 250 timepoints each for the early development and young mouse time ranges across all of the samples. We used the ANOVA test with Bonferroni multiple hypothesis testing correction to compare the gene expression values for each gene across organs in the early development time ranges and again to compare expression of each gene across organs in the young mouse development time ranges.

#### Tabula Muris Senis gene evaluation

To evaluate whether Sagittarius could accurately predict gene expression patterns in aging mouse, we computed the Pearson correlation over time for each of the genes in the aging mouse extrapolated time series. We also computed the Pearson correlation over time for the mouse organ time series measurements in the Evo-devo dataset, which end at P63. Finally, we used the heart and aorta, kidney, and liver tissue data from the single-cell Tabula Muris Senis droplet dataset,^16^ which were the three tissues that aligned with the Evo-devo organs. We computed the average expression at each timepoint for each cell type in the tissue data, and then took the Pearson correlation of the average cell type expression over time. We compare the correlation over time for Sagittarius’s extrapolated data and the measured Evo-devo data to the distribution of cell type correlation in the Tabula Muris Senis dataset.

### Drug dosage similarity network

After training Sagittarius on the complete LINCS dataset we randomly selected 78 distinct doses from the dataset, which ranged from 8.33e-5 to 19.9998, and selected a treatment time of 6 hours. For each drug and cell line treatment in the dataset, we then used Sagittarius to predict the drug-induced expression profile for the treatment at each of the 78 doses with a 6-hour treatment time, using the actual treatment measurements in the dataset as the source sequence. To remove the strong cell-type-specific signal in the profiles, we subtracted Sagittarius’s predicted basal cell line expression from the drug-induced expression vectors.

We took the average over all 78 doses of the differential expression vectors to produce a single 978-dimensional vector for each of the 2,687 treatment combinations. We then computed a similarity matrix *∑* as

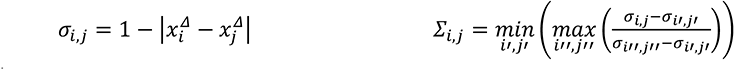

where 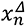 indicates the differential expression of treatment combination *n*. We constructed an average differential expression k-nearest-neighbors (KNN) network *G_KNN_*, beginning from a fully connected graph with edge weights ∑, by first removing all edges with *∑_ij_* < 0.95, then removing the *degree*(*i*) – 50 edges with lowest weight for each node *i*, removing all nodes with degree less than 30, and finally reducing the remaining graph to its largest-connected subgraph. We then ran Louvain community detection^94^ from the Python `community` package^95^ (`python-louvain`) to identify communities in *G_KNN_*. To simplify the analysis, we combined neighboring communities until 4 remained, and then calculated the community IC50 by averaging the individual treatment IC50 over all treatments in the community that were also measured in the GDSC dataset. We visualized *G_KNN_* using the edge-weight spring embedded layout in Cytoscape,^96^ with minimum, maximum, and default edge weights of 0, 1, and 0.5 respectively. We ran 200 average iterations for each node. The spring strength parameter was set to 15, spring rest length to 45, the disconnected spring strength to 0.05, and the disconnected spring rest length to 2000. We did not add any spring strength to avoid collisions, and used 2 layout passes. Finally, we randomized the graph before computing the layout.

### Drug sensitivity prediction

To evaluate Sagittarius’s utility for drug sensitivity prediction, we again randomly selected 78 doses in the LINCS dataset and fixed the treatment time as 6 hours. We then used Sagittarius’s learned weights to compute the transformer encoder’s average key representation over the doses for a given drug and cell line combination, corresponding to the average treatment efficacy relative to the reference space. We fix the treatment time while varying dose in order to best capture the impact of dose on the treatment response, as IC50 is based on drug dose-response _curves._52,58,97

#### GDSC dataset

For the GDSC-based prediction, we computed the key representations for each GDSC^52^-measured combination of drug and cell line, provided that both the drug and cell line appeared somewhere in the LINCS dataset (although not necessarily together). We then constructed one dataset with 271 datapoints using Sagittarius’s representations, and another dataset with 151 datapoints that contained the drug-induced gene expression profiles for treatment combinations that also appeared in the GDSC dataset, denoted Sagittarius and LINCS-GDSC respectively. We ran 3-fold cross-validation on the LINCS-GDSC model, where the test set made up 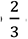 of the data in each fold. We divided Sagittarius’s simulated dataset such that the test split for each fold matched the LINCS-based model split, and used the remaining data for training.

#### CTRP dataset

For this experiment we computed Sagittarius’s key representations for each CTRP^58^-measured experimental combination. The Sagittarius dataset had 2,929 datapoints and the LINCS-CTRP dataset had 625 datapoints.

#### Dataset evaluation

To evaluate the quality of the Sagittarius and LINCS-based IC50 prediction models, we computed the average Spearman correlation between the model’s IC50 predictions and the dataset labels (either from GDSC or CTRP) for each test cell line. To compare overall test performance, we restricted our analysis to cell lines where at least one of the models had significant correlation (Spearman rank-order p-value < 0.05). We used the Spearman correlation between all predicted and measured validation data to quantify validation set performance.

#### Model hyperparameters

We used 3-fold cross validation on the LINCS-based dataset, where 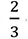 of the data was used as test for each fold. We defined test splits in the Sagittarius dataset to match the LINCS-based test split for each fold. We held out 10% of the remaining training data for both the Sagittarius and LINCS-based datasets to determine the best regression model for the drug sensitivity prediction task in each dataset. We considered both Support Vector Regression (SVR) and MLP-based regressors; for SVR regression configurations, we considered linear, polynomial, and radial-basis-function (RBF) kernels; for MLP regression configurations, we considered a regularization weight *α* ∈ [1e-4, 1e-2, 1, 10]. All other hyperparameters maintained the defaults in `sklearn`.^93^ For the LINCS-based dataset model, the best-performing configuration on the validation data used an SVR regressor with an RBF kernel on the GDSC dataset and polynomial kernel on the CTRP dataset. For the Sagittarius dataset model, the MLP model with *α* = 10 on the GDSC dataset and *α* = 0.01 on the CTRP dataset.

### Cancer gene essentiality prediction

#### Dataset construction

We used the DEMETER^60^ and CERES^61^ versions of the DepMap dataset, which quantify gene essentiality via short hairpin RNAs and CRISPR-Cas9 essentiality screens respectively. For each gene and cell line combination in DepMap, we searched for a drug in the LINCS dataset that listed the given gene as its target, hypothesizing that the drug’s inhibitory effect on a cell line is related to the cell line’s dependency on the target gene.^98^ We constructed a dataset from Sagittarius based on the transformer’s average key representation as in the drug sensitivity prediction dataset, using 78 random doses and a treatment combination in the given cell line and targeting the gene of interest for each DepMap essentiality pair. This resulted in 4,216 and 1,666 datapoints from Sagittarius for the DEMETER and CERES versions respectively. We analogously constructed LINCS-DepMap dataset used the average drug-induced expression across doses for DepMap pairs that matched available LINCS experiments, resulting in 765 and 353 datapoints for the two versions.

#### Dataset evaluation

To evaluate the quality of the Sagittarius- and LINCS-based datasets, we computed the Spearman correlation between the gene essentiality scores measured in the dataset and those predicted by the mode for each test cell line.

#### Model training

We trained a 2-layer MLP regressor with 200- and 100-hidden nodes respectively, ReLU activation functions, MSE loss, and the Adam optimizer^99^ with a learning rate of 1e-3. We used 5-fold cross validation, where 20% of the LINCS-DepMap dataset was used as the test set, and we aligned the Sagittarius dataset’s test set to match the LINCS-DepMap test set. We further held out 10% of the resulting training set for each of the 5 splits to use as a validation set for early model training termination.

### Early cancer patient simulation

#### Mutation profile simulation

To simulate the early-stage sarcoma patient mutation profiles, we trained Sagittarius on all available TCGA mutation data and then predicted mutation probability profiles at 27 survival timepoints, ranging from 203-283 months. Specifically, we selected the longest 27 survival times that appeared somewhere in the initial TCGA mutation dataset, with t ε {203.12, 204.01, 260.70, 208.23, 209.43, 210.51, 210.81, 211.01, 211.73, 212.09, 216.59, 216.75, 225.43, 229.04, 230.72, 232.00, 232.62, 233.44, 234.10, 238.11, 244.32, 244.91, 255.49, 263.07, 275.66, 281.08, 282.69} months (**Supplementary Fig. 23**). We then averaged the mutation profile predictions of the 27 timepoints and identified the 10 genes the model predicted as most likely to be mutated.

#### Differentially expressed gene simulation

To investigate the transcriptional differences between early- and late-stage sarcoma patients, we trained Sagittarius on all available TCGA RNA-seq profiles and then simulated new expression profiles for two synthetic sarcoma patients, one with post-diagnosis survival *t* = 0 and the other with *t* = 244.91, representing late-stage and early-stage tumors respectively. We the identified the *k* = 10 genes that were most overexpressed in the late-stage tumor compared to the early-stage tumor as *PMP2, NDRG1, JUN, SEPT9, PHC2, NCAM1, GFAP, APP, WNK1,* and *RAB10*. Previous work identified *JUN*,^100^ *NCAM1*,^101^ *NDRG1*^102–105^, and *PMP2*^106, 107^ in aggressive sarcomas with poor prognosis and short time-to-metastasis. Furthermore, other work has found *RAB10*,^108^ *SEPT9*,^109^ *PHC2*,^110^ *WNK1*,^111^ and *APP*^112^ overexpression in sarcomas.

## Data availability

The data necessary for reproducing the paper are available in the figshare repository at https://figshare.com/projects/Sagittarius/144771. We provide a more detailed note describing the datasets provided in the figshare repository. Namely, the main experiments can be run with the pre-processed data files provided, and we also include data for the cross-dataset analyses such as the application in drug repurposing. Finally, we provide large-scale simulated data from Sagittarius for the Evo-devo and TCGA datasets.

## Code availability

A python repository including the Sagittarius implementation and code to reproduce the results in this paper is available at https://github.com/addiewc/Sagittarius, with additional details for reproducing results provided in the repository (DOI: https://doi.org/10.5281/zenodo.7879454). We ran experiments on Linux 8.7 with RTX A4000 GPU. We used Python 3.9.7, pytorch 1.9.1, anndata 0.8.0, cudatoolkit 11.1.74, matplotlib 3.4.3, numpy 1.21.2, pandas 1.3.3, pip 21.3.1, pybiomart 0.2.0, python-louvain 0.15, scanpy 1.8.2, scipy 1.7.1, seaborn 0.11.2, sklearn 0.0, statsmodels 0.13.0, tqdm 4.62.3, umap-learn 0.5.1, yaml 0.2.5, lifelines 0.26.5, BioRender Student Plan, and Adobe Illustrator 25.2.3.

## Acknowledgements

Figures were created with BioRender.

## Author Contributions Statement

AW and SW conceptualized the work and designed the method. AW, SW, and JM designed the experiments. AW and MZ ran the experiments, and AW, MZ, and JC developed computational tools for Sagittarius. AW and SW wrote the manuscript and designed the figures.

