## Supplementary Information for "Extrapolating Heterogeneous Time-Series Gene Expression Data using Sagittarius"

### Supplementary Note 1

#### Sagittarius model

We define the input heterogeneous time-series dataset as $\mathcal{D} = \{(x_i, y_i, t_i)\}_{i=1}^{N}$, which we can think of as a collection of measurements associated with irregularly sampled continuous variables. The $x_i \in \mathbb{R}^{T_i \times M}$ are the measured time series input for sequence $i$, where each measurement is $M$-dimensional and the time series is measured at $T_i$ timepoints. The $y_i \in \{1, \ldots, C_j\}^C$ are the $C$ environmental variables for time series $i$, with $y_{i,j} \in \{1, \ldots, C_j\}$ for $C_j$ possible values for the $j$th environmental variable; in the Evo-devo[1] study, for instance, these indicate the species and organ labels of the time series. The $t_i \in \mathbb{R}^{B \times T}$ are the $B$ continuous variables for time series $i$, with $t_{i,j}[r]$ denoting the value of the $j$th continuous variable associated with the $r$th measurement of time series $i$, $x_i[r]$. In particular, $B = 1$ in the Evo-Devo[1] and TCGA[2] studies, while $B = 2$ in the LINCS[3] study, where we model both dose and time as continuous variables. We further assume that $(x_i, y_i, t_i) \sim \mathcal{X}$, where $\mathcal{X}$ is the space of all possible measurements. Sagittarius then predicts a sample $(\hat{x}|y, t) \sim \mathcal{X}$ for a user-specified combination of environmental and continuous variables that may not be measured in dataset $\mathcal{D}$ (**Supplementary Fig. 1**).

Sagittarius first embeds individual measurements to a generative latent space, disentangling the environmental variables as

$$\mu_i[r], \sigma_i[r] = q_\xi(x_i[r], y_i) \qquad z_i[r] \sim \mathcal{N}(\mu_i[r], \sigma_i[r]),$$

where $z_i[r] \in \mathbb{R}^d$ with $d \ll M$. For brevity, we summarize these two steps as $z(x_i, y_i)[r]$. We regularize this learned Gaussian space by imposing the standard-normal prior, $p(z) = \mathcal{N}(0, I)$.

The second component of the model is a continuous transformer.[4] This transformer parameterizes a mapping to and from the shared reference space, facilitating a comparison based on inferred biological age rather than measured timepoint. Given a temporal basis range of interest $\left(\theta_j^{(0)}, \theta_j^{(1)}\right)$ for each of the $j = \{1, \ldots, B\}$ continuous variables and the number of reference timepoints $S + 1 < T$ as model hyperparameters, Sagittarius defines the fixed temporal grid

$$t_{ref,j} \in \mathbb{R}^{S+1}: t_{ref,j}[\tau] = \theta_j^{(0)} + \tau \frac{\theta_j^{(1)} - \theta_j^{(0)}}{S}.$$

We further define the continuous attention embedding function

$$\psi_{h,b}^{enc}(t_b)[v] = sin\left(\omega_{h,b,v}^{enc} t_b + \alpha_{h,b,v}^{enc}\right)$$

for the $b$th continuous variable for each dimension $v \in V$ and attention head $h \in H$, where $V$ and $H$ are model hyperparameters, and $\omega$ and $\alpha$ are fixed scaling and shifting terms. We choose a larger $\omega$ to use higher-frequency embeddings, where the number of cycles of the sine wave for the same input domain increases. This high-frequency sinusoidal embedding both limits the range of embedded timepoints to $\psi_{h,b}^{enc}(t)[v] \in [0,1]$ for all $h, b, t, v$, mitigating potential generalization issues to timepoints not contained in the training set, and explicitly encodes more high-frequency behavior in the data, which neural networks have been shown to learn more slowly than low-frequency patterns.[4,5] We further combine the embeddings for each of the continuous variables to the complete continuous embeddings

$$\psi_h^{enc}(t_i[r]) = \oplus_{b=1}^{B} \psi_{h,b}^{enc}(t_{i,b}[r]) \qquad \psi_h^{enc}(t_{ref}[\tau]) = \oplus_{b=1}^{B} \psi_{h,b}^{enc}(t_{ref,b}[\tau]),$$

where $\oplus$ indicates vector concatenation. Finally, in order to align measured timepoints across environmental conditions, we define the transformer's $h$th attention head key and query for time series $i$ as

$$k_{h,i}^{enc}[r] = f_{h,v}^{enc}(y_i, \psi_h^{enc}(t_i[r])) \qquad q_h^{enc}[\tau] = g_{h,v}^{enc}\left(\psi_h^{enc}(t_{ref}[\tau])\right),$$

where both $k_{h,i}^{enc}[r]$ and $q_h^{enc}[\tau]$ are $d_k$-dimensional vectors, with $d_k$ a hyperparameter. Finally, we use the transformer to map the embedded time series to the reference space as

$$z_{ref}(x_i, y_i, t_i)[\tau] = \sum_{h=1}^{H} \sum_{r=1}^{T} z(x_i, y_i)[r] \frac{exp(\langle k_{h,i}^{enc}[r], q_h^{enc}[\tau]\rangle / \sqrt{d_k})}{\sum_{r'=1}^{T} exp(\langle k_{h,i}^{enc}[r'], q_h^{enc}[\tau]\rangle / \sqrt{d_k})}.$$

An overview of Sagittarius's encoder module including implementation details for $q_\xi, f_{h,v}^{enc}$, and $g_{h,v}^{enc}$ are provided in **Supplementary Fig. 2**.

The decoder layer of our continuous transformer follows a very similar framework, decoding from the regular reference space back to the timepoints of interest. Specifically, we compute

$$\psi_{h,b}^{dec}(t_{j,b}[r])[v] = sin\left(\omega_{h,b,v}^{dec} t_{j,b}[r] + \alpha_{h,b,v}^{dec}\right)$$

$$\psi_h^{dec}(t_j[r]) = \oplus_{b=1}^{B} \psi_{h,b}^{dec}(t_{j,b}[r]) \qquad \psi_h^{dec}(t_{ref}[\tau]) = \oplus_{b=1}^{B} \psi_{h,b}^{dec}(t_{ref,b}[\tau]).$$

We further define

$$k_h^{dec}[\tau] = f_{h,v'}^{dec}\left(\psi_h^{dec}(t_{ref}[\tau])\right) \qquad q_{h,j}^{dec}[r] = g_{h,v'}^{dec}\left(y_j, \psi_h^{dec}(t_j[r])\right)$$

to be the decoding layer's key and query values, respectively. Finally, we convert from the regular time series in the reference space back to the irregular time series with

$$\hat{z}_j(x_i, y_i, t_i, y_j, t_j)[r] = \sum_{h=1}^{H} \sum_{\tau=0}^{S} z_{ref}(x_i, y_i, t_i)[\tau] \frac{exp(\langle k_h^{dec}[\tau], q_{h,j}^{dec}[r]\rangle / \sqrt{d_k})}{\sum_{\tau'=0}^{S} exp(\langle k_h^{dec}[\tau'], q_{h,j}^{dec}[r]\rangle / \sqrt{d_k})}.$$

If we take $j = i$, then this reconstructs the input time series; if we take $j \neq i$, then this generates a new time series from an initial input.

Finally, we convert our time series $\hat{z}_j(x_i, y_i, t_i, y_j, t_j)$ back from the latent embedding space to the data space, with

$$\hat{x}_j(x_i, y_i, t_i, y_j, t_j) = p_\theta(\hat{z}_j(x_i, y_i, t_i, y_j, t_j), y_j).$$

An overview of Sagittarius's decoder module including implementation details for $p_\theta, f^{dec}_{h,v'}$, and $g^{dec}_{h,v'}$ are provided in **Supplementary Fig. 3**.

#### Sagittarius objective

We train our model end-to-end with the Adam optimizer[6] and the loss function $\mathcal{L}(\xi, \nu, \upsilon, \nu', \upsilon', \theta)$, denoted $\mathcal{L}_{Sag}(.)$ for brevity, as

$$\mathcal{L}_{Sag}(.) = \mathbb{E}_{(x_j, y_j, t_j)\in\mathcal{X}}\left[\mathbb{E}_{(x_i, y_i, t_i)\sim\mathcal{D}}\left[\mathbb{E}_{q_\xi(z_i|x_i, y_i)}\left[log\ p_\theta\left(x_j|z_i, y_i, t_i, y_j, t_j\right)\right] - \beta D_{KL}\left(q_\xi(z_i|x_i, y_i) \parallel p(z)\right)\right]\right],$$

where $D_{KL}$ denotes the Kullback-Leibler divergence and $\beta$ is a regularization weighting hyperparameter. During training, we jointly optimized a reconstruction and generation objective. For the reconstruction objective, we set $j = i$ for $i = 1, \cdots, N$ for $N$ total training time series. For the generation objective, we selected a generation target time series $j$ for a source time series $i$, constructing generation examples $(i, j)$ for each task as follows.

- *Evo-devo training*: randomly select five batches of 12 time series from the training dataset. We considered the following four objectives, resulting in $N_{gen} = 48$ generative examples:
  1. mask three observations from each time series and treat the 3 observations as the generation target;
  2. randomly select a second time series with the same species label but different organ label to treat as the generation target;
  3. repeat (2), but use the same organ and different species;
  4. randomly pair the fourth and fifth batches to treat one as the generation target of the other.
- *LINCS training:* randomly select 32 drugs, 32 time series, and 16 pairs of treatment combinations from the training dataset. We considered the following three objectives, resulting in $N_{gen} = 80$ generative examples:
  1. for each drug, identify two training treatment combinations with that drug and treat one as the generation target;
  2. repeat (1), but select treatments for the same cell line;
  3. for each pair of treatment combinations, treat one time series as the generation target of the other.
- *TCGA training*: randomly select three batches of 12 time series from the training dataset. We considered the following two objectives, resulting in $N_{gen} = 24$ generative examples:
  1. mask three observations from each time series and treat the 3 observations as the generation target;

2. randomly pair time series from batches 2 and 3, treating one as the generation target.

We trained Sagittarius with the Adam optimizer[6] with a batch size of 8 time series in the Evo-devo experiments, a batch size of 16 time series in LINCS, and a batch size of 2 time series in the TCGA experiments.

#### Evaluation metrics

We evaluate model performance with four main metrics: RMSE, correlation comparing genes, correlation comparing timepoints, and model AUROC. Given a set of test timepoints $\widetilde{T_\iota} \in [1, \cdots, T_i]$ for time series $i$ with total number of measurements $T_i$, we define model RMSE as the average of each time series RMSE, or

$$RMSE_i = \sqrt{\frac{1}{|\widetilde{T_\iota}|}\sum_{t\in\widetilde{T_\iota}}(\widehat{x_\iota}[t] - x_i[t])^2} \qquad RMSE = \frac{1}{N}\sum_{i=1}^{N} RMSE_i.$$

We further define a model's correlation (comparing genes) as the average of each time series correlation, or

$$\rho_i^{(genes)} = \frac{1}{|\widetilde{T_\iota}|}\sum_{t\in\widetilde{T_\iota}}\rho(\widehat{x_\iota}[t], x_i[t]) \qquad \rho^{(genes)} = \frac{1}{N}\sum_{i=1}^{N}\rho_i^{(genes)},$$

using the Pearson correlation coefficient for $\rho$ in the Evo-devo experiments and the Spearman correlation coefficient for the LINCS and TCGA experiments. Analogously, we define the correlation (comparing timepoints) as

$$\rho_i^{(times)} = \frac{1}{M}\sum_{g=1}^{M}\rho\left(\widehat{x_\iota}[\widetilde{T_\iota}, g], x_i[\widetilde{T_\iota}, g]\right) \qquad \rho^{(times)} = \frac{1}{N}\sum_{i=1}^{N}\rho_i^{(times)},$$

where $x_i[\widetilde{T_\iota}, g]$ and $\widehat{x_\iota}[\widetilde{T_\iota}, g]$ represent the vectors of measured and predicted expression for gene $g$ at test timepoints $\widetilde{T_\iota}$.

Finally, we evaluated the model prediction AUROC for the binary mutation profiles. For this evaluation, we restricted the evaluation genes to those with Augmented Dickey-Fuller (ADF) test p-value > 0.05 in at least $\delta$ cancer types, denoted $\gamma(\delta)$, with $\delta_{THCA} = 2$ and $\delta_{SARC} = 4$, resulting in 9 and 61 usable test splits respectively. We then define the per-patient AUROC as

$$AUROC_i(t) = AUROC(\widehat{x_\iota}[t, \gamma(\delta)], x_i[t, \gamma(\delta)]) \qquad AUROC_i = \frac{1}{|\widetilde{T_\iota}|}\sum_{t\in\widetilde{T_\iota}} AUROC_i(t),$$

using $AUROC$ to indicate the standard AUROC computation between two vectors and excluding any patients with $x_i[t, \gamma(\delta)] = \vec{0}$. To compare models across all test splits in the TCGA setting, we further define the average model AUROC as the average $AUROC_i$ over all test splits $\widetilde{T_\iota}$.

To assess whether Sagittarius statistically outperforms the comparison approaches, we used the one-sided paired t-test between Sagittarius's performance and the best comparison approach's performance per time series. For $\rho^{(genes)}$ and $\rho^{(times)}$, we first computed the Fisher z-transformation[7] of the correlation values, defined as

$$z(r) = \frac{1}{2} ln\left(\frac{1+r}{1-r}\right)$$

for some correlation coefficient $r$, to normalize the t-test inputs.

#### Comparison approaches

**Mean**: The mean model predicts data as $\hat{x}_i[t] = \frac{1}{T}\sum_{r=1}^{T} x_i[r]$; that is, the predicted expression for each gene at any timepoint of interest is the average of the gene expression across all measured timepoints.

**Linear**: The linear baseline model defines a weight

$$\lambda_{i,t} = 0 \text{ if } t < min(t_i);$$
$$\lambda_{i,t} = 1 \text{ if } t > max(t_i);$$
$$\lambda_{i,t} = \max_{r \in t_i:\, r \le t}\left(\min_{s \in t_i:\, s \ge t}\left(\frac{t-r}{s-r}\right)\right) \text{ otherwise.}$$

Then, the linear model predicts expression at time $t$ as

$$\hat{x}_i[t] = \max_{r \in t_i:\, r \le t}(1-\lambda_{i,t})x_i[r] + \min_{s \in t_i:\, s \ge t}\lambda_{i,t}x_i[s].$$

Note that, in the extrapolation setting, the linear baseline therefore predicts a gene expression vector identical to the expression vector of the nearest temporal measurement.

**Neural ODE**: We trained a set of neural ODE models[8] that each model measurements from a single combination of environmental variables. As the environmental conditions $y_i$ are constant within a single sequence, we reduce the task inputs to $(x\,, t)$. We compute an ODE for both the forward and backward direction of the sequence as

$$\tilde{x}_{\rightarrow}[r] = \max_{s \in t_i:\, s \le r} x[s] + \int_{t=s}^{r} f_\theta(x[t])dt$$
$$\tilde{x}_{\leftarrow}[r] = \min_{s \in t_i:\, s \ge r} x[s] + \int_{t=s}^{r} g_\phi(x[t])dt.$$

In the case where $\widetilde{x_{\rightarrow}}$ or $\widetilde{x_{\leftarrow}}$ requires extrapolation (i.e., there is no such $s$ to satisfy the constraint), we set $\widetilde{x_{\rightarrow}}[r] = \tilde{x}_{\leftarrow}[r]$. In order to empirically compute the integrals we used the python torchdiffeq package,[8,9] and parameterized $f_\theta(.)$ and $g_\phi(.)$ using a multi-layer perceptron (MLP). Finally, we combined the forward and backward results to produce the final estimate

$$\hat{x}[r] = \frac{1}{2}(\tilde{x}_{\rightarrow}[r] + \tilde{x}_{\leftarrow}[r]).$$

We trained the model using the Adam optimizer[6] and a batch size of 1 time series.

**RNN**: We learned a set of single-sequence bidirectional gated recurrent unit (GRU)[10] models to learn the dynamics for a single $(x_i, y_i, t_i)$ sequence, again reducing the problem input to $(x, t)$. We define a time step $\Delta_t = 1$ between observations of interest. At each timepoint, we computed $z_r = q_\phi(x[r])$ for an MLP $q_\phi(.)$ as the embedding for each observation in the time series, and computed

$$\hat{z}_t^{\rightarrow} = q_\phi\big(p_\theta(h_t^{\rightarrow})\big) \qquad h_{t+1}^{\rightarrow} = g_\xi^{(gru\rightarrow)}(z_t, h_t^{\rightarrow}) \text{ if } t \in t_i \text{ ; } h_{t+1}^{\rightarrow} = g_\xi^{(gru\rightarrow)}(\hat{z}_t^{\rightarrow}, h_t^{\rightarrow}) \text{ otherwise,}$$

where $h_0^{\rightarrow} = 0$. Similarly, we define the backward GRU as

$$\hat{z}_t^{\leftarrow} = q_{\phi'}\big(p_{\theta'}(h_t^{\leftarrow})\big) \qquad h_{t+1}^{\leftarrow} = g_{\xi'}^{(gru\leftarrow)}(z_t, h_t^{\leftarrow}) \text{ if } t \in t_i \text{ ; } h_{t+1}^{\leftarrow} = g_{\xi'}^{(gru\leftarrow)}(\hat{z}_t^{\leftarrow}, h_t^{\leftarrow}) \text{ otherwise,}$$

with $h_T^{\leftarrow} = 0$. Finally, we combine the forward- and backward directions to produce the simulated gene expression profile

$$\hat{x}[r] = p_\theta\left(\frac{1}{2}(h_r^{\rightarrow} + h_r^{\leftarrow})\right)$$

for an MLP $p_\theta(.)$. We trained the model end-to-end with the Adam optimizer[6] and a batch size of 1 time series.

**mTAN**: We trained a discretized multi-time attention network (mTAN)[4] using the Adam optimizer.[6] As the mTAN module does not handle environmental variables, for each time series $(x_i, y_i, t_i)$ the model received the reduced input $(x_i, t_i)$.

**cVAE**: We trained a conditional variational autoencoder (cVAE)[11] to learn $p(x_i[r] \mid y_i, t_i[r])$. We trained the model using the Adam optimizer[6] and a batch size of 512 gene expression measurements. This is the only model that can be applied to multiple continuous variables out-of-the box, so we use it as the comparison method for the LINCS experiments.

**CPA**: We trained a compositional perturbation autoencoder (CPA)[12] with the Adam optimizer[6] and a batch size of 128. In the Evo-devo experiments, we considered the time to be independent of the organ label (the covariate) and dependent on the species label (the perturbation).

#### Model hyperparameter selection

**Evo-devo:** For each deep learning model, we randomly select 20% of the measurements available at training time to use as validation data, which we use for hyperparameter tuning and early training termination, where we stop training when the validation loss has not reached a new minimum for 250 epochs. For each comparison approach, we conducted an approximate hyperparameter grid search on the late-timepoint Evo-devo extrapolation task, selecting 50 experimental configurations per model from a set of possible values per hyperparameter. Given some redundant experimental configurations, this resulted in 49 unique experiments for cVAE, 50 for CPA, 50 for mTAN, 42 for RNN, 22 for Neural ODE, and 50 for PRESCIENT. We selected the best model based on Pearson correlation comparing genes in the validation set. We then used the same hyperparameters for all Evo-devo experiments, including extrapolation to early timepoints.

For the cVAE, CPA, mTAN, and RNN models, we considered an embedding dimension $d \in [8, 16, 32, 64, 128]$, eventually selecting $d_{cvae} = 128$, $d_{cpa} = 8$, $d_{mtan} = 64$, and $d_{rnn} = 16$. We also considered learning rates $\eta \in$ [1e-4, 1e-3, 1e-2] for the cVAE, CPA autoencoder, mTAN, RNN, Neural ODE, and PRESCIENT, setting $\eta_{cvae}$ = 1e-3, $\eta_{cpa}$ = 1e-4, $\eta_{mtan}$ = 1e-3, $\eta_{rnn}$ = 1e-4, $\eta_{node}$ = 1e-4, and $\eta_{prescient}$ = 1e-4. For cVAE, CPA, RNN, and PRESCIENT, we considered autoencoder widths $W \in [512, 1024, 2046]$ and depths $D \in [1,2,3]$. Based on validation results, we set $(W, D)_{cvae} = (1024, 1)$, $(W, D)_{cpa} = (1024,3)$, $(W, D)_{rnn} = (1024, 1)$, and $(W, D)_{prescient} = (1024, 1)$. For the Neural ODE and PRESCIENT models, we considered time steps of $\Delta_t \in$ [0.1, 0.5, 1.0] and set $\Delta_{t,node}$ = 0.1 and $\Delta_{t,prescient}$ = 0.1. We also used validation performance to select the cVAE's regularization weight as $\beta = 0 \in [0, \frac{1}{6}, \frac{1}{3}, \frac{1}{2}, \frac{2}{3}, \frac{5}{6}, 1]$. Similarly, we selected CPA's adversarial learning rate as 1e-4 ∈ [1e-4, 1e-3], adversary width and depth as $(256, 1) \in [128, 256, 512] \times [1,2,3]$, adversary steps as $2 \in [2,4,8]$, patience as $5 \in [3,4,5,6]$, doser type as logsigm ∈ [logsigm, sigm, mlp], and doser learning rate as 1e-4 ∈ [1e-4, 1e-3]. We also considered an MLP doser width and depth in $[16, 32, 64] \times [1,2]$, but did not use these in the final configuration as the logsigm doser had stronger validation performance. For the mTAN model, we also determined whether to learn the temporal embedding as true ∈ [true, false], set the number of transformer attention heads to $H = 4 \in [4, 8, 12]$, and used the overall time embedding dimension $d_{temp} = 8 \in [1,2,3] \times H$. For Neural ODE, we used an ODE depth of $3 \in [1,2,3]$. Finally, for the PRESCIENT model we also selected a sinkhorn blur and scaling of (0.5, 0.75) ∈ [0.1, 0.5, 1.0] × [0.25, 0.5, 0.75]. We used $200 \in [50,100,200]$ pretrain epochs and $200 \in [50,100,200]$ train burnin epochs. We set the train $\tau = 0 \in$ [0, 1e-6, 1e-4] and gradient clipping to 0 ∈ [0, 0.25, 0.5], indicating no clipping.

We conducted a reduced hyperparameter grid search for the Sagittarius model, again using the validation dataset for both early stopping and hyperparameter selection. We fixed the embedding dimension to $d = 32$, learning rate to $\eta_{sag}$ = 1e-3, autoencoder width to 1024, number of heads $H$ to 8, number of reference points $S + 1$ to 4, time embedding dimension $d_{temp}$ to 4. We then searched over an autoencoder depth in [2, 3];

regularization weight $\beta$ in $[0, \frac{1}{6}, 1]$; autoencoder categorical embedding dimension $d_{yae}$ in [2, 8]; and transformer categorical embedding dimension $d_{ytr}$ in [4, 8]. Based on the validation performance, we selected an autoencoder depth of 3, regularization $\beta$ of $\frac{1}{6}$, and autoencoder and transformer categorical embedding dimensions $d_{yae} = 2$ and $d_{ytr} = 4$.

We then conducted an ablation study for different configuration results to investigate Sagittarius's robustness to various hyperparameter settings. **Supplementary Fig. 6-8** show Sagittarius's test performance for the late development extrapolation experiment under different model hyperparameter settings. We emphasize that this study was conducted after hyperparameters were fixed for all Evo-Devo Sagittarius experiments, and the test results were never used outside of the ablation. In the ablation study, we tuned one hyperparameter while setting the remainder to our optimal configuration as determined by the validation set. We found that Sagittarius's improvement over existing approaches was robust to several hyperparameter settings, including the number of attention heads $H$, the transformer's environmental variable embedding dimension $d_{ytr}$, and the number of generative datapoints used to train the model.

Unsurprisingly, we found that Sagittarius was sensitive to choice of learning rate with respect to all of the evaluation metrics that we measured. We also found that the regularization weight $\beta$ impacted model performance; in the ablation study, we considered $\beta \in [0.0, 0.1667, 1.0]$, where $0.1667 = \frac{m}{N}$, where $m = 8$ was the batch size and $N = 48$ was the dataset size. This finding suggests that both under-regularizing and over-regularizing the model leads to worse performance. Sagittarius's performance is also impacted by the choice of autoencoder structure and latent embedding dimension, perhaps reflecting the need for a relatively tight information bottleneck ($d < 64$) given the limited size of the dataset, and the importance of disentangling non-linear species- and organ-specific patterns from the gene expression measurements before reaching the reference space.

**LINCS:** For Sagittarius and the cVAE model, we randomly partitioned the data into an 80% training, 10% validation, and 10% test split. We terminated model training when the validation loss had not decreased for at least 100 epochs and returned the model with lowest validation loss. For the Sagittarius and cVAE models, we set the embedding dimension $d_{cvae} = d_{sag} = 16$, learning rate $\eta_{cvae} = \eta_{sag} =$ 1e-3, and the autoencoder width and depth $(W, D)_{cvae} = (W, D)_{sag} = (128,2)$. We set the regularization weight $\beta_{cvae} = 1.0$ and $\beta_{sag} = 0.25$. For Sagittarius, we further set the number of attention heads $H = 8$, total number of reference points as 16 (spanning both dose and time, as a $4 \times 4$ grid in the reference space), time embedding dimension $d_{temp}$ to 8 for dose and 4 for time, and both autoencoder and transformer categorical embedding dimensions as $d_{yae} = d_{ytr} = 8$.

**TCGA:** For each of the deep learning models, we used 20% of the available data at train time as a validation set for training termination and hyperparameter selection where applicable. We set an embedding dim $d_{rnn} = d_{sag} = 16$. We set the learning rate $\eta_{rnn} = \eta_{node} =$ 1e-4, and $\eta_{sag} =$ 1e-3. For the RNN, we used an autoencoder of width 256 and depth 1. For Neural ODE, we set the timestep $\Delta_t = 1$ and ODE depth 1. For Sagittarius, we set the regularization weight $\beta = 1$, number of attention heads $H = 8$, number of reference points $S + 1 = 4$, temporal embedding dimension $d_{temp} = 2$, and the autoencoder and transformer categorical embedding dimensions $d_{yae} = 8$ and $d_{ytr} = 4$ respectively. We further set the autoencoder width to be 256, and then searched over depths in [1, 2, 3], using the validation data to select a depth-3 and depth-2 autoencoder for the THCA and SARC experiments respectively.

#### TCGA mutation time series filtering

Although our TCGA time series formulation enables extrapolation to cancer patients with good prognosis, censored patients may have an event time that differs dramatically from their actual survival time, confounding the temporal ordering of patients. For the TCGA gene expression experiments, we excluded all censored patients. For the mutation experiments, we leveraged recent techniques from Learning with Noisy Labels[13,14] to identify censored patients who likely had a death event shortly after the reported event time while excluding patients more likely to survive long after losing contact with the study. We constructed a neural network $f$ for each cancer type to predict $\widehat{t[r]} = f(x[r])$. Given the patient's binary censoring label $c[r]$, where $c[r] = 0$ indicates that the $r$th patient had a censored death event, we defined the per-patient loss as

$$\mathcal{L}_{individual}(x[r], t[r], \hat{t}[r], c[r]) = \mathbb{1}[c[r] = 0]\, |\hat{t} - t|_1 + \mathbb{1}[c[r] = 1]\, max(t - \hat{t}, 0),$$

thereby not penalizing the model for overestimating the survival time of a censored patient, and trained $f(\cdot)$ with the sum of all individual patients' losses and a regularization term. We used the stochastic gradient descent (SGD) optimizer with learning rate 0.1, a single hidden layer with 32 neurons and regularization penalty weight of 0.3 for the L2 norm of the weights. We trained the model for 2,500 epochs and selected the best model from the epoch with maximal concordance index among the observed patients. We then computed the absolute error $|f(x[r]) - t[r]|$ for each patient and fit a beta distribution $\beta_{obs}$ and $\beta_{cens}$ to the absolute errors for the observed and censored patient error observations respectively using the scipy[15] python package. Finally, for each censored patient, if the absolute error associated with their event time was more likely to be generated by $\beta_{obs}$ than $\beta_{cens}$, or if the absolute error was smaller than the error associated with one or more observed patients, we retained that patient; otherwise, we excluded the patient from further analysis (**Supplementary Fig. 21**). After filtering for all cancers, our mutation dataset contained 2,297 cancer patients.

##### Comparison with natural language processing transformer architecture

Sagittarius's transformer-based architecture is built around a continuous transformer formulation,[4] which is inspired by the traditional transformer architecture[16] used in natural language processing (NLP). In the NLP context, the transformer *attends* to tokens (words) in the input sentence, comparing token similarity to encode the input (**Supplementary Fig. 28a)**. Importantly, the query and (key, value) pair are embeddings computed to represent the tokens (words). In the NLP transformer encoder, then, the attention mechanism functions within a sequence-to-sequence framework by comparing key- and query- embeddings representing different words. More similar keys and queries then have a larger contribution for the computed value to be associated with the query; keys that are dissimilar to the query will have a smaller contribution to the query's computed value.

The continuous transformer[4] builds upon this idea by using timepoint embeddings as the keys and queries in the transformer architecture (**Supplementary Fig. 28b**). While the value still represents the measurement associated with the key's timepoint, this incorporates a continuous notion of time, including enabling a query during inference for a timepoint not used during model training. Sagittarius further extends the continuous transformer to include timepoint keys and queries that are also conditioned on the environmental variables, enabling implicit time series alignment and accounting for varying temporal rates in the input trajectories.

#### Supplementary Tables

##### Supplementary Table 1

| species | fisher statistic | fisher p-value |
|---|---|---|
| Chicken | 0.2807991494 | 0.0002050198533 |
| Human | 0.08345916698 | 3.85E-17 |
| Mouse | 0.4646453882 | 0.003192069078 |
| Opossum | 0.1269005312 | 2.91E-13 |
| Rabbit | 0.4279736999 | 1.12E-05 |

| species | fisher statistic | fisher p-value |
|---|---|---|
| Rat | 0.07055946967 | 9.35E-18 |
| RhesusMacaque | 0.3590132827 | 1.03E-05 |

**Supplementary Table 1** Comparison of Sagittarius's extrapolation performance for house-keeping genes (HKGs) and non-HKGs. After identifying HKGs for each species based on the standard deviation of gene expression and generating a set of genes for each species that Sagittarius predicts with at least 0.4 test Pearson correlation comparing timepoints, we report the two-sided Fisher exact test statistic and p-value for whether HKGs have different performance than non-HKGs.

**Supplementary Table 2**

| GO term | Number of species enriched | Species | Organ | p-value |
|---|---|---|---|---|
| transmembrane receptor protein tyrosine kinase signaling pathway | 7 | Chicken | Brain | 1.915271E-06 |
| transmembrane receptor protein tyrosine kinase signaling pathway | 7 | Chicken | Kidney | 1.003212E-05 |
| transmembrane receptor protein tyrosine kinase signaling pathway | 7 | Human | Brain | 3.335650E-05 |
| transmembrane receptor protein tyrosine kinase signaling pathway | 7 | Human | Cerebellum | 6.871604E-03 |
| transmembrane receptor protein tyrosine kinase signaling pathway | 7 | Human | Kidney | 6.995938E-06 |

| GO term | Number of species enriched | Species | Organ | p-value |
|---|---|---|---|---|
| transmembrane receptor protein tyrosine kinase signaling pathway | 7 | Human | Liver | 6.437116E-04 |
| transmembrane receptor protein tyrosine kinase signaling pathway | 7 | Mouse | Ovary | 2.927464E-02 |
| transmembrane receptor protein tyrosine kinase signaling pathway | 7 | Mouse | Testis | 4.091004E-03 |
| transmembrane receptor protein tyrosine kinase signaling pathway | 7 | Opossum | Kidney | 2.895958E-06 |
| transmembrane receptor protein tyrosine kinase signaling pathway | 7 | Opossum | Liver | 1.386478E-05 |
| transmembrane receptor protein tyrosine kinase signaling pathway | 7 | Opossum | Ovary | 5.505152E-04 |
| transmembrane receptor protein tyrosine kinase signaling pathway | 7 | Opossum | Testis | 6.172717E-03 |

| GO term | Number of species enriched | Species | Organ | p-value |
|---|---|---|---|---|
| transmembrane receptor protein tyrosine kinase signaling pathway | 7 | Rabbit | Liver | 2.760365E-04 |
| transmembrane receptor protein tyrosine kinase signaling pathway | 7 | Rabbit | Testis | 2.105139E-04 |
| transmembrane receptor protein tyrosine kinase signaling pathway | 7 | Rat | Brain | 2.061978E-05 |
| transmembrane receptor protein tyrosine kinase signaling pathway | 7 | Rat | Liver | 3.226703E-02 |
| transmembrane receptor protein tyrosine kinase signaling pathway | 7 | Rat | Testis | 2.060156E-02 |
| transmembrane receptor protein tyrosine kinase signaling pathway | 7 | RhesusMacaque | Brain | 3.244361E-10 |
| transmembrane receptor protein tyrosine kinase signaling pathway | 7 | RhesusMacaque | Kidney | 3.951913E-05 |
| small GTPase binding | 7 | Chicken | Brain | 4.054573E-02 |
| small GTPase binding | 7 | Chicken | Kidney | 1.893029E-03 |

| GO term | Number of species enriched | Species | Organ | p-value |
|---|---|---|---|---|
| small GTPase binding | 7 | Human | Brain | 2.901160E-07 |
| small GTPase binding | 7 | Human | Kidney | 4.213821E-05 |
| small GTPase binding | 7 | Human | Liver | 1.468640E-02 |
| small GTPase binding | 7 | Mouse | Cerebellum | 2.396363E-02 |
| small GTPase binding | 7 | Mouse | Ovary | 4.343957E-04 |
| small GTPase binding | 7 | Mouse | Testis | 1.739027E-06 |
| small GTPase binding | 7 | Opossum | Kidney | 1.296523E-06 |
| small GTPase binding | 7 | Opossum | Liver | 1.412536E-03 |
| small GTPase binding | 7 | Rabbit | Liver | 2.884628E-03 |
| small GTPase binding | 7 | Rabbit | Testis | 1.581520E-02 |
| small GTPase binding | 7 | Rat | Brain | 5.239981E-06 |
| small GTPase binding | 7 | Rat | Testis | 5.744777E-05 |
| small GTPase binding | 7 | RhesusMacaque | Testis | 5.160330E-04 |
| response to organonitrogen compound | 7 | Chicken | Kidney | 2.245133E-03 |
| response to organonitrogen compound | 7 | Human | Brain | 4.765489E-02 |
| response to organonitrogen compound | 7 | Human | Testis | 2.643214E-02 |
| response to organonitrogen compound | 7 | Mouse | Ovary | 3.462941E-03 |

| GO term | Number of species enriched | Species | Organ | p-value |
|---|---|---|---|---|
| response to organonitrogen compound | 7 | Opossum | Liver | 1.478138E-03 |
| response to organonitrogen compound | 7 | Opossum | Ovary | 1.799559E-02 |
| response to organonitrogen compound | 7 | Rabbit | Cerebellum | 4.240515E-02 |
| response to organonitrogen compound | 7 | Rat | Brain | 8.047883E-03 |
| response to organonitrogen compound | 7 | Rat | Liver | 1.888863E-02 |
| response to organonitrogen compound | 7 | RhesusMacaque | Liver | 1.001987E-03 |
| regulation of protein kinase activity | 7 | Chicken | Brain | 1.849025E-06 |
| regulation of protein kinase activity | 7 | Human | Brain | 1.199655E-02 |
| regulation of protein kinase activity | 7 | Human | Kidney | 2.360261E-02 |
| regulation of protein kinase activity | 7 | Mouse | Kidney | 1.584154E-03 |
| regulation of protein kinase activity | 7 | Mouse | Ovary | 2.040221E-03 |
| regulation of protein kinase activity | 7 | Mouse | Testis | 2.377961E-03 |
| regulation of protein kinase activity | 7 | Opossum | Heart | 3.144693E-02 |
| regulation of protein kinase activity | 7 | Opossum | Kidney | 2.550958E-03 |

| GO term | Number of species enriched | Species | Organ | p-value |
|---|---|---|---|---|
| regulation of protein kinase activity | 7 | Opossum | Liver | 3.093968E-02 |
| regulation of protein kinase activity | 7 | Opossum | Ovary | 1.912599E-03 |
| regulation of protein kinase activity | 7 | Rabbit | Kidney | 1.247950E-02 |
| regulation of protein kinase activity | 7 | Rabbit | Testis | 4.184600E-02 |
| regulation of protein kinase activity | 7 | Rat | Liver | 1.781173E-03 |
| regulation of protein kinase activity | 7 | RhesusMacaque | Brain | 4.471173E-04 |
| regulation of neuron projection development | 7 | Chicken | Brain | 2.026943E-03 |
| regulation of neuron projection development | 7 | Chicken | Kidney | 8.235158E-04 |
| regulation of neuron projection development | 7 | Human | Brain | 1.349509E-03 |
| regulation of neuron projection development | 7 | Human | Kidney | 2.849987E-04 |
| regulation of neuron projection development | 7 | Human | Liver | 2.242675E-07 |
| regulation of neuron projection development | 7 | Human | Ovary | 1.070618E-02 |

| GO term | Number of species enriched | Species | Organ | p-value |
|---|---|---|---|---|
| regulation of neuron projection development | 7 | Human | Testis | 7.234747E-03 |
| regulation of neuron projection development | 7 | Mouse | Testis | 2.376188E-03 |
| regulation of neuron projection development | 7 | Opossum | Kidney | 1.373725E-02 |
| regulation of neuron projection development | 7 | Opossum | Liver | 2.100523E-02 |
| regulation of neuron projection development | 7 | Rabbit | Liver | 3.067226E-04 |
| regulation of neuron projection development | 7 | Rat | Brain | 6.987796E-07 |
| regulation of neuron projection development | 7 | RhesusMacaque | Brain | 1.497547E-03 |
| regulation of neuron projection development | 7 | RhesusMacaque | Kidney | 1.678636E-03 |
| regulation of neuron projection development | 7 | RhesusMacaque | Liver | 5.064254E-03 |
| regulation of kinase activity | 7 | Chicken | Brain | 4.579910E-06 |
| regulation of kinase activity | 7 | Human | Brain | 1.672572E-02 |
| regulation of kinase activity | 7 | Human | Kidney | 2.556435E-02 |

| GO term | Number of species enriched | Species | Organ | p-value |
|---|---|---|---|---|
| regulation of kinase activity | 7 | Mouse | Kidney | 6.825682E-03 |
| regulation of kinase activity | 7 | Mouse | Ovary | 6.474059E-03 |
| regulation of kinase activity | 7 | Mouse | Testis | 4.311476E-03 |
| regulation of kinase activity | 7 | Opossum | Heart | 2.195761E-02 |
| regulation of kinase activity | 7 | Opossum | Kidney | 4.201631E-03 |
| regulation of kinase activity | 7 | Opossum | Ovary | 1.203891E-03 |
| regulation of kinase activity | 7 | Rabbit | Kidney | 9.471528E-03 |
| regulation of kinase activity | 7 | Rat | Liver | 3.222481E-03 |
| regulation of kinase activity | 7 | RhesusMacaque | Brain | 5.956894E-03 |
| regulation of kinase activity | 7 | RhesusMacaque | Kidney | 2.643671E-02 |
| regulation of cell projection organization | 7 | Chicken | Brain | 4.460439E-06 |
| regulation of cell projection organization | 7 | Chicken | Kidney | 8.615754E-04 |
| regulation of cell projection organization | 7 | Chicken | Ovary | 2.578387E-02 |
| regulation of cell projection organization | 7 | Human | Brain | 4.886149E-05 |
| regulation of cell projection organization | 7 | Human | Kidney | 2.810985E-05 |
| regulation of cell projection organization | 7 | Human | Liver | 5.109937E-08 |

| GO term | Number of species enriched | Species | Organ | p-value |
|---|---|---|---|---|
| regulation of cell projection organization | 7 | Human | Ovary | 2.081702E-02 |
| regulation of cell projection organization | 7 | Mouse | Testis | 4.185357E-04 |
| regulation of cell projection organization | 7 | Opossum | Kidney | 1.223778E-05 |
| regulation of cell projection organization | 7 | Opossum | Liver | 1.334726E-03 |
| regulation of cell projection organization | 7 | Opossum | Ovary | 9.371669E-04 |
| regulation of cell projection organization | 7 | Rabbit | Liver | 1.480143E-05 |
| regulation of cell projection organization | 7 | Rabbit | Testis | 2.398474E-02 |
| regulation of cell projection organization | 7 | Rat | Brain | 1.570184E-08 |
| regulation of cell projection organization | 7 | RhesusMacaque | Brain | 1.003567E-05 |
| regulation of cell projection organization | 7 | RhesusMacaque | Heart | 4.867821E-02 |
| regulation of cell projection organization | 7 | RhesusMacaque | Kidney | 8.628221E-05 |
| regulation of cell projection organization | 7 | RhesusMacaque | Liver | 1.139356E-04 |
| regulation of cell morphogenesis | 7 | Chicken | Brain | 2.568745E-04 |
| regulation of cell morphogenesis | 7 | Chicken | Kidney | 6.451065E-04 |

| GO term | Number of species enriched | Species | Organ | p-value |
|---|---|---|---|---|
| regulation of cell morphogenesis | 7 | Human | Brain | 1.120621E-03 |
| regulation of cell morphogenesis | 7 | Human | Kidney | 1.196184E-03 |
| regulation of cell morphogenesis | 7 | Human | Testis | 4.951151E-02 |
| regulation of cell morphogenesis | 7 | Mouse | Testis | 3.628094E-02 |
| regulation of cell morphogenesis | 7 | Opossum | Kidney | 4.260492E-04 |
| regulation of cell morphogenesis | 7 | Opossum | Liver | 9.724616E-03 |
| regulation of cell morphogenesis | 7 | Rabbit | Liver | 2.026743E-02 |
| regulation of cell morphogenesis | 7 | Rat | Brain | 1.011501E-02 |
| regulation of cell morphogenesis | 7 | RhesusMacaque | Brain | 2.558424E-03 |
| regulation of cell morphogenesis | 7 | RhesusMacaque | Kidney | 4.405979E-04 |
| regulation of GTPase activity | 7 | Chicken | Brain | 1.247288E-02 |
| regulation of GTPase activity | 7 | Human | Brain | 2.844199E-03 |
| regulation of GTPase activity | 7 | Human | Kidney | 2.863261E-03 |
| regulation of GTPase activity | 7 | Mouse | Testis | 1.103345E-03 |

| GO term | Number of species enriched | Species | Organ | p-value |
|---|---|---|---|---|
| regulation of GTPase activity | 7 | Opossum | Kidney | 7.905775E-06 |
| regulation of GTPase activity | 7 | Rabbit | Liver | 2.415217E-02 |
| regulation of GTPase activity | 7 | Rat | Brain | 2.168258E-07 |
| regulation of GTPase activity | 7 | RhesusMacaque | Liver | 1.014225E-03 |
| regulation of GTPase activity | 7 | RhesusMacaque | Testis | 4.043296E-02 |
| protein serine/threonine kinase activity | 7 | Chicken | Ovary | 3.099745E-05 |
| protein serine/threonine kinase activity | 7 | Human | Brain | 3.546568E-07 |
| protein serine/threonine kinase activity | 7 | Human | Cerebellum | 4.980028E-02 |
| protein serine/threonine kinase activity | 7 | Human | Kidney | 1.841681E-06 |
| protein serine/threonine kinase activity | 7 | Human | Liver | 3.553961E-03 |
| protein serine/threonine kinase activity | 7 | Mouse | Testis | 1.126599E-06 |
| protein serine/threonine kinase activity | 7 | Opossum | Brain | 3.291208E-02 |
| protein serine/threonine kinase activity | 7 | Opossum | Kidney | 2.104955E-10 |
| protein serine/threonine kinase activity | 7 | Opossum | Liver | 1.269783E-04 |

| GO term | Number of species enriched | Species | Organ | p-value |
|---|---|---|---|---|
| protein serine/threonine kinase activity | 7 | Opossum | Ovary | 3.198137E-04 |
| protein serine/threonine kinase activity | 7 | Opossum | Testis | 2.244006E-03 |
| protein serine/threonine kinase activity | 7 | Rabbit | Liver | 1.303101E-03 |
| protein serine/threonine kinase activity | 7 | Rabbit | Testis | 8.660366E-05 |
| protein serine/threonine kinase activity | 7 | Rat | Brain | 6.698093E-04 |
| protein serine/threonine kinase activity | 7 | Rat | Cerebellum | 8.197034E-05 |
| protein serine/threonine kinase activity | 7 | Rat | Kidney | 9.151189E-04 |
| protein serine/threonine kinase activity | 7 | Rat | Testis | 1.073770E-05 |
| protein serine/threonine kinase activity | 7 | RhesusMacaque | Brain | 3.460661E-05 |
| protein serine/threonine kinase activity | 7 | RhesusMacaque | Testis | 1.770104E-08 |
| protein phosphorylation | 7 | Chicken | Brain | 2.572473E-04 |
| protein phosphorylation | 7 | Chicken | Kidney | 1.363805E-05 |
| protein phosphorylation | 7 | Chicken | Liver | 7.715129E-03 |
| protein phosphorylation | 7 | Human | Brain | 7.547553E-08 |
| protein phosphorylation | 7 | Human | Cerebellum | 7.595363E-04 |

| GO term | Number of species enriched | Species | Organ | p-value |
|---|---|---|---|---|
| protein phosphorylation | 7 | Human | Kidney | 1.195098E-11 |
| protein phosphorylation | 7 | Human | Liver | 9.562360E-05 |
| protein phosphorylation | 7 | Human | Ovary | 4.726091E-02 |
| protein phosphorylation | 7 | Mouse | Brain | 1.771195E-02 |
| protein phosphorylation | 7 | Mouse | Cerebellum | 7.175638E-03 |
| protein phosphorylation | 7 | Mouse | Ovary | 3.886044E-03 |
| protein phosphorylation | 7 | Mouse | Testis | 7.640980E-07 |
| protein phosphorylation | 7 | Opossum | Brain | 1.937407E-03 |
| protein phosphorylation | 7 | Opossum | Heart | 4.534296E-03 |
| protein phosphorylation | 7 | Opossum | Kidney | 6.634613E-11 |
| protein phosphorylation | 7 | Opossum | Liver | 3.853969E-09 |
| protein phosphorylation | 7 | Opossum | Ovary | 1.078387E-05 |
| protein phosphorylation | 7 | Opossum | Testis | 7.647868E-05 |
| protein phosphorylation | 7 | Rabbit | Brain | 2.008274E-02 |
| protein phosphorylation | 7 | Rabbit | Cerebellum | 7.904741E-04 |
| protein phosphorylation | 7 | Rabbit | Kidney | 4.525550E-02 |
| protein phosphorylation | 7 | Rabbit | Liver | 1.017211E-07 |
| protein phosphorylation | 7 | Rabbit | Testis | 3.456342E-04 |
| protein phosphorylation | 7 | Rat | Brain | 6.660134E-07 |

| GO term | Number of species enriched | Species | Organ | p-value |
|---|---|---|---|---|
| protein phosphorylation | 7 | Rat | Cerebellum | 1.080871E-05 |
| protein phosphorylation | 7 | Rat | Kidney | 4.220425E-04 |
| protein phosphorylation | 7 | Rat | Liver | 2.699376E-04 |
| protein phosphorylation | 7 | Rat | Testis | 4.710772E-07 |
| protein phosphorylation | 7 | RhesusMacaque | Brain | 1.006441E-08 |
| protein phosphorylation | 7 | RhesusMacaque | Heart | 9.367290E-03 |
| protein phosphorylation | 7 | RhesusMacaque | Kidney | 6.716673E-04 |
| protein phosphorylation | 7 | RhesusMacaque | Testis | 9.466368E-07 |
| protein kinase activity | 7 | Chicken | Brain | 1.241348E-04 |
| protein kinase activity | 7 | Chicken | Kidney | 4.580190E-04 |
| protein kinase activity | 7 | Chicken | Liver | 9.057696E-04 |
| protein kinase activity | 7 | Chicken | Ovary | 4.786726E-05 |
| protein kinase activity | 7 | Human | Brain | 5.930706E-08 |
| protein kinase activity | 7 | Human | Cerebellum | 1.537176E-04 |
| protein kinase activity | 7 | Human | Kidney | 4.139872E-10 |
| protein kinase activity | 7 | Human | Liver | 8.744322E-05 |
| protein kinase activity | 7 | Mouse | Cerebellum | 3.240626E-03 |
| protein kinase activity | 7 | Mouse | Ovary | 2.879856E-02 |
| protein kinase activity | 7 | Mouse | Testis | 6.787577E-07 |

| GO term | Number of species enriched | Species | Organ | p-value |
|---|---|---|---|---|
| protein kinase activity | 7 | Opossum | Brain | 2.006127E-02 |
| protein kinase activity | 7 | Opossum | Kidney | 4.462216E-11 |
| protein kinase activity | 7 | Opossum | Liver | 1.161402E-07 |
| protein kinase activity | 7 | Opossum | Ovary | 5.027879E-05 |
| protein kinase activity | 7 | Opossum | Testis | 1.804546E-06 |
| protein kinase activity | 7 | Rabbit | Kidney | 2.254575E-02 |
| protein kinase activity | 7 | Rabbit | Liver | 8.841757E-06 |
| protein kinase activity | 7 | Rabbit | Testis | 4.096903E-05 |
| protein kinase activity | 7 | Rat | Brain | 3.142638E-06 |
| protein kinase activity | 7 | Rat | Cerebellum | 7.384666E-05 |
| protein kinase activity | 7 | Rat | Heart | 5.537376E-03 |
| protein kinase activity | 7 | Rat | Kidney | 5.470656E-05 |
| protein kinase activity | 7 | Rat | Liver | 1.843457E-02 |
| protein kinase activity | 7 | Rat | Testis | 1.573216E-05 |
| protein kinase activity | 7 | RhesusMacaque | Brain | 1.739749E-07 |
| protein kinase activity | 7 | RhesusMacaque | Heart | 7.046782E-03 |
| protein kinase activity | 7 | RhesusMacaque | Kidney | 1.658321E-02 |
| protein kinase activity | 7 | RhesusMacaque | Liver | 2.048943E-02 |
| protein kinase activity | 7 | RhesusMacaque | Testis | 1.263545E-06 |

| GO term | Number of species enriched | Species | Organ | p-value |
|---|---|---|---|---|
| positive regulation of protein kinase activity | 7 | Chicken | Brain | 1.647257E-06 |
| positive regulation of protein kinase activity | 7 | Human | Brain | 1.329134E-02 |
| positive regulation of protein kinase activity | 7 | Human | Kidney | 2.866411E-03 |
| positive regulation of protein kinase activity | 7 | Mouse | Kidney | 9.592319E-03 |
| positive regulation of protein kinase activity | 7 | Mouse | Ovary | 1.580516E-02 |
| positive regulation of protein kinase activity | 7 | Mouse | Testis | 4.715463E-03 |
| positive regulation of protein kinase activity | 7 | Opossum | Kidney | 3.876692E-04 |
| positive regulation of protein kinase activity | 7 | Opossum | Liver | 3.346217E-02 |
| positive regulation of protein kinase activity | 7 | Opossum | Ovary | 2.484980E-02 |
| positive regulation of protein kinase activity | 7 | Rabbit | Liver | 3.319102E-02 |
| positive regulation of protein kinase activity | 7 | Rat | Liver | 3.728906E-03 |

| GO term | Number of species enriched | Species | Organ | p-value |
|---|---|---|---|---|
| positive regulation of protein kinase activity | 7 | RhesusMacaque | Brain | 3.507275E-03 |
| positive regulation of GTPase activity | 7 | Chicken | Brain | 4.255670E-02 |
| positive regulation of GTPase activity | 7 | Human | Brain | 4.320545E-03 |
| positive regulation of GTPase activity | 7 | Human | Kidney | 3.706406E-03 |
| positive regulation of GTPase activity | 7 | Mouse | Testis | 4.034894E-04 |
| positive regulation of GTPase activity | 7 | Opossum | Kidney | 4.885450E-06 |
| positive regulation of GTPase activity | 7 | Rabbit | Liver | 2.923723E-02 |
| positive regulation of GTPase activity | 7 | Rat | Brain | 2.154760E-06 |
| positive regulation of GTPase activity | 7 | RhesusMacaque | Liver | 1.320050E-03 |
| positive regulation of GTPase activity | 7 | RhesusMacaque | Testis | 3.958740E-02 |
| phosphotransferase activity, alcohol group as acceptor | 7 | Chicken | Brain | 5.877922E-07 |

| GO term | Number of species enriched | Species | Organ | p-value |
|---|---|---|---|---|
| phosphotransferase activity, alcohol group as acceptor | 7 | Chicken | Kidney | 7.347460E-06 |
| phosphotransferase activity, alcohol group as acceptor | 7 | Chicken | Liver | 5.977017E-04 |
| phosphotransferase activity, alcohol group as acceptor | 7 | Chicken | Ovary | 2.028924E-04 |
| phosphotransferase activity, alcohol group as acceptor | 7 | Human | Brain | 2.844424E-10 |
| phosphotransferase activity, alcohol group as acceptor | 7 | Human | Cerebellum | 6.209356E-06 |
| phosphotransferase activity, alcohol group as acceptor | 7 | Human | Kidney | 2.098348E-12 |
| phosphotransferase activity, alcohol group as acceptor | 7 | Human | Liver | 1.857324E-04 |
| phosphotransferase activity, alcohol group as acceptor | 7 | Human | Ovary | 9.751635E-03 |
| phosphotransferase activity, alcohol group as acceptor | 7 | Human | Testis | 1.881821E-02 |
| phosphotransferase activity, alcohol group as acceptor | 7 | Mouse | Brain | 9.672515E-04 |
| phosphotransferase activity, alcohol group as acceptor | 7 | Mouse | Cerebellum | 7.386004E-05 |

| GO term | Number of species enriched | Species | Organ | p-value |
|---|---|---|---|---|
| phosphotransferase activity, alcohol group as acceptor | 7 | Mouse | Testis | 6.168425E-09 |
| phosphotransferase activity, alcohol group as acceptor | 7 | Opossum | Brain | 1.995401E-03 |
| phosphotransferase activity, alcohol group as acceptor | 7 | Opossum | Heart | 8.029263E-03 |
| phosphotransferase activity, alcohol group as acceptor | 7 | Opossum | Kidney | 3.070931E-11 |
| phosphotransferase activity, alcohol group as acceptor | 7 | Opossum | Liver | 4.210508E-09 |
| phosphotransferase activity, alcohol group as acceptor | 7 | Opossum | Ovary | 5.497368E-07 |
| phosphotransferase activity, alcohol group as acceptor | 7 | Opossum | Testis | 2.906873E-09 |
| phosphotransferase activity, alcohol group as acceptor | 7 | Rabbit | Kidney | 9.984209E-04 |
| phosphotransferase activity, alcohol group as acceptor | 7 | Rabbit | Liver | 7.675810E-07 |
| phosphotransferase activity, alcohol group as acceptor | 7 | Rabbit | Testis | 1.507014E-06 |
| phosphotransferase activity, alcohol group as acceptor | 7 | Rat | Brain | 6.108005E-09 |

| GO term | Number of species enriched | Species | Organ | p-value |
|---|---|---|---|---|
| phosphotransferase activity, alcohol group as acceptor | 7 | Rat | Cerebellum | 3.972323E-04 |
| phosphotransferase activity, alcohol group as acceptor | 7 | Rat | Heart | 4.805418E-04 |
| phosphotransferase activity, alcohol group as acceptor | 7 | Rat | Kidney | 8.292033E-06 |
| phosphotransferase activity, alcohol group as acceptor | 7 | Rat | Liver | 1.063413E-03 |
| phosphotransferase activity, alcohol group as acceptor | 7 | Rat | Testis | 6.904498E-07 |
| phosphotransferase activity, alcohol group as acceptor | 7 | RhesusMacaque | Brain | 3.941270E-09 |
| phosphotransferase activity, alcohol group as acceptor | 7 | RhesusMacaque | Cerebellum | 6.123734E-03 |
| phosphotransferase activity, alcohol group as acceptor | 7 | RhesusMacaque | Heart | 1.638384E-03 |
| phosphotransferase activity, alcohol group as acceptor | 7 | RhesusMacaque | Kidney | 4.489712E-04 |
| phosphotransferase activity, alcohol group as acceptor | 7 | RhesusMacaque | Liver | 3.002856E-03 |
| phosphotransferase activity, alcohol group as acceptor | 7 | RhesusMacaque | Testis | 4.055598E-07 |

| GO term | Number of species enriched | Species | Organ | p-value |
|---|---|---|---|---|
| organophosphate metabolic process | 7 | Chicken | Brain | 7.620738E-03 |
| organophosphate metabolic process | 7 | Chicken | Kidney | 1.034205E-04 |
| organophosphate metabolic process | 7 | Human | Brain | 5.820413E-03 |
| organophosphate metabolic process | 7 | Human | Cerebellum | 5.900296E-04 |
| organophosphate metabolic process | 7 | Human | Ovary | 1.947816E-03 |
| organophosphate metabolic process | 7 | Mouse | Brain | 1.296476E-04 |
| organophosphate metabolic process | 7 | Mouse | Ovary | 4.233715E-02 |
| organophosphate metabolic process | 7 | Mouse | Testis | 1.392117E-02 |
| organophosphate metabolic process | 7 | Opossum | Cerebellum | 8.789060E-03 |
| organophosphate metabolic process | 7 | Opossum | Liver | 2.893564E-03 |
| organophosphate metabolic process | 7 | Rabbit | Brain | 8.351446E-05 |
| organophosphate metabolic process | 7 | Rabbit | Kidney | 1.836520E-03 |
| organophosphate metabolic process | 7 | Rabbit | Testis | 6.709083E-04 |
| organophosphate metabolic process | 7 | Rat | Kidney | 3.989979E-02 |

| GO term | Number of species enriched | Species | Organ | p-value |
|---|---|---|---|---|
| organophosphate metabolic process | 7 | Rat | Liver | 1.808681E-06 |
| organophosphate metabolic process | 7 | Rat | Ovary | 1.356851E-03 |
| organophosphate metabolic process | 7 | RhesusMacaque | Cerebellum | 6.895509E-03 |
| organophosphate metabolic process | 7 | RhesusMacaque | Liver | 9.928799E-04 |
| nucleolus | 7 | Chicken | Cerebellum | 4.649031E-14 |
| nucleolus | 7 | Chicken | Kidney | 1.443376E-08 |
| nucleolus | 7 | Chicken | Testis | 3.236238E-05 |
| nucleolus | 7 | Human | Brain | 1.046159E-05 |
| nucleolus | 7 | Human | Cerebellum | 1.286527E-03 |
| nucleolus | 7 | Human | Liver | 5.743246E-03 |
| nucleolus | 7 | Human | Ovary | 7.009480E-07 |
| nucleolus | 7 | Mouse | Brain | 3.385917E-08 |
| nucleolus | 7 | Mouse | Cerebellum | 3.212285E-03 |
| nucleolus | 7 | Mouse | Heart | 7.674515E-07 |
| nucleolus | 7 | Mouse | Ovary | 5.595974E-04 |
| nucleolus | 7 | Opossum | Brain | 2.123131E-09 |
| nucleolus | 7 | Opossum | Liver | 3.392219E-05 |
| nucleolus | 7 | Rabbit | Brain | 4.592077E-07 |
| nucleolus | 7 | Rabbit | Cerebellum | 2.848907E-07 |
| nucleolus | 7 | Rabbit | Heart | 4.659361E-08 |
| nucleolus | 7 | Rabbit | Kidney | 2.078293E-04 |
| nucleolus | 7 | Rabbit | Liver | 2.895637E-05 |
| nucleolus | 7 | Rabbit | Ovary | 4.410804E-11 |
| nucleolus | 7 | Rabbit | Testis | 4.311261E-06 |
| nucleolus | 7 | Rat | Cerebellum | 5.034846E-03 |

| GO term | Number of species enriched | Species | Organ | p-value |
|---|---|---|---|---|
| nucleolus | 7 | Rat | Kidney | 1.161593E-10 |
| nucleolus | 7 | Rat | Testis | 3.879232E-04 |
| nucleolus | 7 | RhesusMacaque | Heart | 2.120845E-02 |
| nuclear membrane | 7 | Chicken | Cerebellum | 7.802244E-06 |
| nuclear membrane | 7 | Human | Brain | 3.728824E-02 |
| nuclear membrane | 7 | Human | Cerebellum | 2.101536E-03 |
| nuclear membrane | 7 | Human | Liver | 3.310409E-03 |
| nuclear membrane | 7 | Mouse | Heart | 3.763977E-02 |
| nuclear membrane | 7 | Opossum | Brain | 9.349223E-04 |
| nuclear membrane | 7 | Opossum | Liver | 3.276380E-03 |
| nuclear membrane | 7 | Rabbit | Cerebellum | 1.386762E-03 |
| nuclear membrane | 7 | Rabbit | Kidney | 2.310675E-03 |
| nuclear membrane | 7 | Rabbit | Ovary | 2.461980E-02 |
| nuclear membrane | 7 | Rat | Cerebellum | 3.685624E-03 |
| nuclear membrane | 7 | Rat | Kidney | 3.852801E-05 |
| nuclear membrane | 7 | Rat | Testis | 7.242447E-03 |
| nuclear membrane | 7 | RhesusMacaque | Testis | 3.888868E-02 |
| neuron projection guidance | 7 | Chicken | Brain | 7.224890E-05 |
| neuron projection guidance | 7 | Chicken | Kidney | 2.377626E-03 |

| GO term | Number of species enriched | Species | Organ | p-value |
|---|---|---|---|---|
| neuron projection guidance | 7 | Human | Brain | 2.324266E-03 |
| neuron projection guidance | 7 | Human | Kidney | 1.909929E-02 |
| neuron projection guidance | 7 | Human | Liver | 3.906340E-07 |
| neuron projection guidance | 7 | Human | Ovary | 2.468000E-03 |
| neuron projection guidance | 7 | Mouse | Ovary | 2.179078E-04 |
| neuron projection guidance | 7 | Opossum | Kidney | 1.941807E-05 |
| neuron projection guidance | 7 | Opossum | Liver | 1.459979E-04 |
| neuron projection guidance | 7 | Opossum | Ovary | 3.972078E-02 |
| neuron projection guidance | 7 | Rabbit | Liver | 9.024715E-07 |
| neuron projection guidance | 7 | Rat | Brain | 1.685131E-02 |
| neuron projection guidance | 7 | Rat | Liver | 3.223755E-06 |
| neuron projection guidance | 7 | Rat | Testis | 3.331650E-02 |
| neuron projection guidance | 7 | RhesusMacaque | Brain | 3.185710E-05 |
| neuron projection guidance | 7 | RhesusMacaque | Heart | 1.248585E-02 |

| GO term | Number of species enriched | Species | Organ | p-value |
|---|---|---|---|---|
| neuron projection guidance | 7 | RhesusMacaque | Kidney | 2.892582E-03 |
| neuron projection | 7 | Chicken | Kidney | 9.070257E-04 |
| neuron projection | 7 | Human | Brain | 4.565291E-07 |
| neuron projection | 7 | Human | Cerebellum | 1.625130E-04 |
| neuron projection | 7 | Human | Kidney | 5.749157E-03 |
| neuron projection | 7 | Human | Liver | 4.427333E-08 |
| neuron projection | 7 | Human | Testis | 3.182853E-06 |
| neuron projection | 7 | Mouse | Cerebellum | 9.378547E-03 |
| neuron projection | 7 | Mouse | Kidney | 1.352348E-04 |
| neuron projection | 7 | Mouse | Ovary | 3.645048E-03 |
| neuron projection | 7 | Mouse | Testis | 1.172701E-03 |
| neuron projection | 7 | Opossum | Cerebellum | 2.952293E-02 |
| neuron projection | 7 | Opossum | Kidney | 3.731590E-05 |
| neuron projection | 7 | Opossum | Liver | 4.163914E-04 |
| neuron projection | 7 | Opossum | Ovary | 4.976462E-03 |
| neuron projection | 7 | Rabbit | Brain | 2.249977E-02 |
| neuron projection | 7 | Rabbit | Liver | 1.373218E-07 |
| neuron projection | 7 | Rabbit | Ovary | 3.300111E-02 |
| neuron projection | 7 | Rabbit | Testis | 2.922196E-04 |

| GO term | Number of species enriched | Species | Organ | p-value |
|---|---|---|---|---|
| neuron projection | 7 | Rat | Brain | 1.314477E-08 |
| neuron projection | 7 | Rat | Cerebellum | 4.821143E-04 |
| neuron projection | 7 | Rat | Ovary | 1.481364E-04 |
| neuron projection | 7 | Rat | Testis | 2.409480E-02 |
| neuron projection | 7 | RhesusMacaque | Brain | 1.585386E-09 |
| neuron projection | 7 | RhesusMacaque | Heart | 1.080769E-03 |
| neuron projection | 7 | RhesusMacaque | Kidney | 4.172523E-09 |
| neuron projection | 7 | RhesusMacaque | Liver | 3.883572E-03 |
| negative regulation of cell cycle | 7 | Chicken | Cerebellum | 9.503540E-05 |
| negative regulation of cell cycle | 7 | Chicken | Kidney | 9.969764E-06 |
| negative regulation of cell cycle | 7 | Chicken | Ovary | 2.023288E-02 |
| negative regulation of cell cycle | 7 | Human | Brain | 2.364929E-06 |
| negative regulation of cell cycle | 7 | Human | Cerebellum | 2.493383E-05 |
| negative regulation of cell cycle | 7 | Human | Ovary | 8.831448E-04 |
| negative regulation of cell cycle | 7 | Mouse | Brain | 3.305867E-04 |
| negative regulation of cell cycle | 7 | Mouse | Cerebellum | 3.977966E-04 |

| GO term | Number of species enriched | Species | Organ | p-value |
| --- | --- | --- | --- | --- |
| negative regulation of cell cycle | 7 | Mouse | Heart | 2.962266E-02 |
| negative regulation of cell cycle | 7 | Mouse | Ovary | 9.338755E-06 |
| negative regulation of cell cycle | 7 | Mouse | Testis | 1.835092E-05 |
| negative regulation of cell cycle | 7 | Opossum | Brain | 4.706488E-07 |
| negative regulation of cell cycle | 7 | Opossum | Cerebellum | 1.571851E-03 |
| negative regulation of cell cycle | 7 | Opossum | Heart | 8.528556E-03 |
| negative regulation of cell cycle | 7 | Opossum | Liver | 1.530200E-07 |
| negative regulation of cell cycle | 7 | Opossum | Ovary | 3.696722E-02 |
| negative regulation of cell cycle | 7 | Rabbit | Brain | 1.616290E-05 |
| negative regulation of cell cycle | 7 | Rabbit | Cerebellum | 1.420333E-09 |
| negative regulation of cell cycle | 7 | Rabbit | Heart | 7.221687E-07 |
| negative regulation of cell cycle | 7 | Rabbit | Kidney | 2.895035E-05 |
| negative regulation of cell cycle | 7 | Rabbit | Liver | 1.096614E-03 |
| negative regulation of cell cycle | 7 | Rabbit | Ovary | 3.556922E-07 |

| GO term | Number of species enriched | Species | Organ | p-value |
|---|---|---|---|---|
| negative regulation of cell cycle | 7 | Rabbit | Testis | 8.611601E-06 |
| negative regulation of cell cycle | 7 | Rat | Cerebellum | 2.952748E-06 |
| negative regulation of cell cycle | 7 | Rat | Kidney | 1.441190E-04 |
| negative regulation of cell cycle | 7 | Rat | Liver | 2.195877E-03 |
| negative regulation of cell cycle | 7 | Rat | Testis | 3.546816E-03 |
| negative regulation of cell cycle | 7 | RhesusMacaque | Cerebellum | 1.276728E-03 |
| mitotic cell cycle process | 7 | Chicken | Cerebellum | 4.512436E-17 |
| mitotic cell cycle process | 7 | Chicken | Heart | 8.493634E-04 |
| mitotic cell cycle process | 7 | Chicken | Kidney | 3.323412E-16 |
| mitotic cell cycle process | 7 | Chicken | Ovary | 7.251037E-06 |
| mitotic cell cycle process | 7 | Chicken | Testis | 6.853257E-03 |
| mitotic cell cycle process | 7 | Human | Brain | 2.298532E-14 |
| mitotic cell cycle process | 7 | Human | Cerebellum | 5.122411E-11 |
| mitotic cell cycle process | 7 | Human | Ovary | 1.952558E-12 |
| mitotic cell cycle process | 7 | Human | Testis | 8.686666E-05 |
| mitotic cell cycle process | 7 | Mouse | Brain | 1.112023E-13 |
| mitotic cell cycle process | 7 | Mouse | Cerebellum | 1.084557E-15 |

| GO term | Number of species enriched | Species | Organ | p-value |
|---|---|---|---|---|
| mitotic cell cycle process | 7 | Mouse | Heart | 9.632778E-08 |
| mitotic cell cycle process | 7 | Mouse | Ovary | 2.066835E-10 |
| mitotic cell cycle process | 7 | Mouse | Testis | 3.068569E-10 |
| mitotic cell cycle process | 7 | Opossum | Brain | 3.098548E-19 |
| mitotic cell cycle process | 7 | Opossum | Cerebellum | 1.100323E-04 |
| mitotic cell cycle process | 7 | Opossum | Heart | 2.259167E-03 |
| mitotic cell cycle process | 7 | Opossum | Kidney | 4.383256E-04 |
| mitotic cell cycle process | 7 | Opossum | Liver | 4.801817E-10 |
| mitotic cell cycle process | 7 | Rabbit | Brain | 2.198396E-17 |
| mitotic cell cycle process | 7 | Rabbit | Cerebellum | 3.462544E-20 |
| mitotic cell cycle process | 7 | Rabbit | Heart | 1.346306E-12 |
| mitotic cell cycle process | 7 | Rabbit | Kidney | 6.323046E-11 |
| mitotic cell cycle process | 7 | Rabbit | Liver | 3.840709E-05 |
| mitotic cell cycle process | 7 | Rabbit | Ovary | 1.213055E-17 |
| mitotic cell cycle process | 7 | Rabbit | Testis | 1.590272E-10 |
| mitotic cell cycle process | 7 | Rat | Cerebellum | 5.855330E-16 |
| mitotic cell cycle process | 7 | Rat | Kidney | 3.112403E-11 |
| mitotic cell cycle process | 7 | Rat | Liver | 2.100650E-03 |
| mitotic cell cycle process | 7 | Rat | Testis | 5.687297E-07 |

| GO term | Number of species enriched | Species | Organ | p-value |
|---|---|---|---|---|
| mitotic cell cycle process | 7 | RhesusMacaque | Cerebellum | 1.977741E-05 |
| mitotic cell cycle | 7 | Chicken | Cerebellum | 5.659791E-28 |
| mitotic cell cycle | 7 | Chicken | Heart | 2.235707E-05 |
| mitotic cell cycle | 7 | Chicken | Kidney | 1.111372E-22 |
| mitotic cell cycle | 7 | Chicken | Ovary | 1.963601E-05 |
| mitotic cell cycle | 7 | Chicken | Testis | 1.767045E-08 |
| mitotic cell cycle | 7 | Human | Brain | 9.975533E-20 |
| mitotic cell cycle | 7 | Human | Cerebellum | 6.799599E-20 |
| mitotic cell cycle | 7 | Human | Liver | 6.835672E-05 |
| mitotic cell cycle | 7 | Human | Ovary | 2.398560E-20 |
| mitotic cell cycle | 7 | Human | Testis | 3.206414E-05 |
| mitotic cell cycle | 7 | Mouse | Brain | 1.249311E-20 |
| mitotic cell cycle | 7 | Mouse | Cerebellum | 1.413843E-21 |
| mitotic cell cycle | 7 | Mouse | Heart | 8.236958E-14 |
| mitotic cell cycle | 7 | Mouse | Ovary | 9.831517E-19 |
| mitotic cell cycle | 7 | Mouse | Testis | 2.495460E-15 |
| mitotic cell cycle | 7 | Opossum | Brain | 9.885533E-32 |
| mitotic cell cycle | 7 | Opossum | Cerebellum | 8.392591E-07 |
| mitotic cell cycle | 7 | Opossum | Heart | 1.151056E-04 |

| GO term | Number of species enriched | Species | Organ | p-value |
|---|---|---|---|---|
| mitotic cell cycle | 7 | Opossum | Kidney | 7.159273E-03 |
| mitotic cell cycle | 7 | Opossum | Liver | 9.599013E-14 |
| mitotic cell cycle | 7 | Opossum | Ovary | 6.449992E-03 |
| mitotic cell cycle | 7 | Rabbit | Brain | 1.603819E-25 |
| mitotic cell cycle | 7 | Rabbit | Cerebellum | 3.029661E-30 |
| mitotic cell cycle | 7 | Rabbit | Heart | 1.275367E-20 |
| mitotic cell cycle | 7 | Rabbit | Kidney | 5.262535E-16 |
| mitotic cell cycle | 7 | Rabbit | Liver | 5.062580E-11 |
| mitotic cell cycle | 7 | Rabbit | Ovary | 1.698281E-26 |
| mitotic cell cycle | 7 | Rabbit | Testis | 1.127946E-15 |
| mitotic cell cycle | 7 | Rat | Cerebellum | 2.295967E-24 |
| mitotic cell cycle | 7 | Rat | Kidney | 9.065585E-15 |
| mitotic cell cycle | 7 | Rat | Liver | 2.003627E-10 |
| mitotic cell cycle | 7 | Rat | Testis | 5.154927E-11 |
| mitotic cell cycle | 7 | RhesusMacaque | Cerebellum | 1.053488E-08 |
| microtubule organizing center | 7 | Chicken | Cerebellum | 1.101529E-06 |
| microtubule organizing center | 7 | Chicken | Kidney | 1.629337E-08 |
| microtubule organizing center | 7 | Chicken | Ovary | 1.852278E-08 |

| GO term | Number of species enriched | Species | Organ | p-value |
|---|---|---|---|---|
| microtubule organizing center | 7 | Chicken | Testis | 2.922900E-03 |
| microtubule organizing center | 7 | Human | Brain | 2.427590E-14 |
| microtubule organizing center | 7 | Human | Cerebellum | 4.932157E-02 |
| microtubule organizing center | 7 | Human | Kidney | 9.047131E-05 |
| microtubule organizing center | 7 | Human | Liver | 3.389713E-06 |
| microtubule organizing center | 7 | Human | Ovary | 3.325311E-03 |
| microtubule organizing center | 7 | Mouse | Cerebellum | 3.146534E-09 |
| microtubule organizing center | 7 | Mouse | Ovary | 2.570971E-07 |
| microtubule organizing center | 7 | Mouse | Testis | 2.548683E-09 |
| microtubule organizing center | 7 | Opossum | Brain | 2.843531E-03 |
| microtubule organizing center | 7 | Opossum | Heart | 3.809015E-02 |
| microtubule organizing center | 7 | Opossum | Kidney | 2.722511E-10 |
| microtubule organizing center | 7 | Opossum | Liver | 6.280380E-12 |
| microtubule organizing center | 7 | Opossum | Ovary | 1.046128E-07 |

| GO term | Number of species enriched | Species | Organ | p-value |
|---|---|---|---|---|
| microtubule organizing center | 7 | Opossum | Testis | 1.561141E-03 |
| microtubule organizing center | 7 | Rabbit | Brain | 9.887462E-07 |
| microtubule organizing center | 7 | Rabbit | Cerebellum | 1.816113E-04 |
| microtubule organizing center | 7 | Rabbit | Heart | 3.122591E-02 |
| microtubule organizing center | 7 | Rabbit | Kidney | 2.323955E-04 |
| microtubule organizing center | 7 | Rabbit | Liver | 2.085630E-08 |
| microtubule organizing center | 7 | Rabbit | Ovary | 1.388289E-02 |
| microtubule organizing center | 7 | Rabbit | Testis | 8.949438E-09 |
| microtubule organizing center | 7 | Rat | Brain | 1.597222E-02 |
| microtubule organizing center | 7 | Rat | Cerebellum | 1.738238E-07 |
| microtubule organizing center | 7 | Rat | Kidney | 3.719926E-08 |
| microtubule organizing center | 7 | Rat | Testis | 5.556563E-07 |
| microtubule organizing center | 7 | RhesusMacaque | Heart | 1.879926E-02 |
| microtubule organizing center | 7 | RhesusMacaque | Kidney | 7.591682E-05 |

| GO term | Number of species enriched | Species | Organ | p-value |
|---|---|---|---|---|
| microtubule organizing center | 7 | RhesusMacaque | Testis | 4.487229E-04 |
| microtubule | 7 | Chicken | Kidney | 1.802678E-03 |
| microtubule | 7 | Human | Brain | 7.292825E-05 |
| microtubule | 7 | Human | Ovary | 2.529047E-03 |
| microtubule | 7 | Human | Testis | 2.088811E-02 |
| microtubule | 7 | Mouse | Cerebellum | 4.091737E-03 |
| microtubule | 7 | Mouse | Ovary | 2.202348E-02 |
| microtubule | 7 | Mouse | Testis | 2.836038E-03 |
| microtubule | 7 | Opossum | Heart | 3.109391E-02 |
| microtubule | 7 | Rabbit | Brain | 3.812166E-05 |
| microtubule | 7 | Rabbit | Ovary | 2.529970E-03 |
| microtubule | 7 | Rabbit | Testis | 2.609649E-04 |
| microtubule | 7 | Rat | Kidney | 2.457777E-03 |
| microtubule | 7 | Rat | Testis | 3.638354E-02 |
| microtubule | 7 | RhesusMacaque | Brain | 9.734275E-03 |
| kinetochore | 7 | Chicken | Cerebellum | 3.933247E-05 |
| kinetochore | 7 | Chicken | Kidney | 1.569894E-03 |
| kinetochore | 7 | Human | Cerebellum | 9.551367E-03 |
| kinetochore | 7 | Human | Ovary | 4.018412E-04 |
| kinetochore | 7 | Mouse | Brain | 3.962748E-02 |
| kinetochore | 7 | Mouse | Cerebellum | 5.301375E-06 |
| kinetochore | 7 | Mouse | Ovary | 4.060486E-02 |
| kinetochore | 7 | Mouse | Testis | 1.933303E-02 |
| kinetochore | 7 | Opossum | Brain | 2.919338E-06 |
| kinetochore | 7 | Rabbit | Brain | 1.442292E-07 |
| kinetochore | 7 | Rabbit | Cerebellum | 6.645770E-06 |
| kinetochore | 7 | Rabbit | Heart | 1.601580E-02 |
| kinetochore | 7 | Rabbit | Ovary | 1.394723E-03 |

| GO term | Number of species enriched | Species | Organ | p-value |
|---|---|---|---|---|
| kinetochore | 7 | Rat | Cerebellum | 1.258810E-05 |
| kinetochore | 7 | Rat | Kidney | 7.838599E-05 |
| kinetochore | 7 | RhesusMacaque | Cerebellum | 8.102242E-03 |
| kinase activity | 7 | Chicken | Brain | 6.395023E-06 |
| kinase activity | 7 | Chicken | Kidney | 6.548200E-05 |
| kinase activity | 7 | Chicken | Liver | 1.537669E-03 |
| kinase activity | 7 | Chicken | Ovary | 9.286156E-05 |
| kinase activity | 7 | Human | Brain | 1.065436E-08 |
| kinase activity | 7 | Human | Cerebellum | 4.993975E-05 |
| kinase activity | 7 | Human | Kidney | 2.934524E-10 |
| kinase activity | 7 | Human | Liver | 1.547051E-04 |
| kinase activity | 7 | Human | Ovary | 3.656713E-02 |
| kinase activity | 7 | Mouse | Brain | 4.932305E-03 |
| kinase activity | 7 | Mouse | Cerebellum | 9.260115E-04 |
| kinase activity | 7 | Mouse | Liver | 4.368286E-02 |
| kinase activity | 7 | Mouse | Testis | 8.210072E-08 |
| kinase activity | 7 | Opossum | Brain | 2.570157E-03 |
| kinase activity | 7 | Opossum | Heart | 1.339160E-02 |
| kinase activity | 7 | Opossum | Kidney | 3.117368E-10 |
| kinase activity | 7 | Opossum | Liver | 4.196031E-08 |
| kinase activity | 7 | Opossum | Ovary | 1.149334E-05 |
| kinase activity | 7 | Opossum | Testis | 9.968027E-08 |
| kinase activity | 7 | Rabbit | Kidney | 5.083610E-03 |
| kinase activity | 7 | Rabbit | Liver | 1.514406E-06 |
| kinase activity | 7 | Rabbit | Testis | 4.174937E-06 |
| kinase activity | 7 | Rat | Brain | 5.597614E-07 |
| kinase activity | 7 | Rat | Cerebellum | 2.345016E-03 |
| kinase activity | 7 | Rat | Heart | 1.987698E-03 |
| kinase activity | 7 | Rat | Kidney | 4.391884E-05 |

| GO term | Number of species enriched | Species | Organ | p-value |
|---|---|---|---|---|
| kinase activity | 7 | Rat | Liver | 4.708015E-03 |
| kinase activity | 7 | Rat | Testis | 3.536137E-06 |
| kinase activity | 7 | RhesusMacaque | Brain | 2.115737E-07 |
| kinase activity | 7 | RhesusMacaque | Cerebellum | 9.735309E-03 |
| kinase activity | 7 | RhesusMacaque | Heart | 4.919732E-03 |
| kinase activity | 7 | RhesusMacaque | Kidney | 4.575126E-03 |
| kinase activity | 7 | RhesusMacaque | Liver | 4.619411E-03 |
| kinase activity | 7 | RhesusMacaque | Testis | 1.092270E-05 |
| gene expression | 7 | Chicken | Cerebellum | 1.530305E-17 |
| gene expression | 7 | Chicken | Kidney | 4.751067E-06 |
| gene expression | 7 | Chicken | Testis | 5.230875E-08 |
| gene expression | 7 | Human | Brain | 1.129761E-06 |
| gene expression | 7 | Human | Cerebellum | 7.932664E-04 |
| gene expression | 7 | Human | Ovary | 2.573917E-10 |
| gene expression | 7 | Mouse | Brain | 7.790319E-13 |
| gene expression | 7 | Mouse | Cerebellum | 4.968032E-06 |
| gene expression | 7 | Mouse | Heart | 3.461195E-15 |
| gene expression | 7 | Mouse | Ovary | 8.566487E-07 |
| gene expression | 7 | Opossum | Brain | 3.686241E-17 |
| gene expression | 7 | Opossum | Heart | 4.664229E-03 |
| gene expression | 7 | Opossum | Liver | 5.287903E-07 |
| gene expression | 7 | Rabbit | Brain | 3.355319E-11 |

| GO term | Number of species enriched | Species | Organ | p-value |
| --- | --- | --- | --- | --- |
| gene expression | 7 | Rabbit | Cerebellum | 2.321957E-14 |
| gene expression | 7 | Rabbit | Heart | 7.184885E-10 |
| gene expression | 7 | Rabbit | Ovary | 5.191445E-16 |
| gene expression | 7 | Rabbit | Testis | 1.716424E-05 |
| gene expression | 7 | Rat | Cerebellum | 2.349349E-06 |
| gene expression | 7 | Rat | Kidney | 5.687937E-03 |
| gene expression | 7 | Rat | Liver | 2.653322E-02 |
| gene expression | 7 | Rat | Testis | 1.194467E-11 |
| gene expression | 7 | RhesusMacaque | Heart | 3.258485E-02 |
| cytoskeleton organization | 7 | Chicken | Brain | 2.556251E-03 |
| cytoskeleton organization | 7 | Chicken | Cerebellum | 2.290099E-02 |
| cytoskeleton organization | 7 | Chicken | Kidney | 2.605769E-04 |
| cytoskeleton organization | 7 | Chicken | Testis | 1.275763E-03 |
| cytoskeleton organization | 7 | Human | Brain | 3.219530E-10 |
| cytoskeleton organization | 7 | Human | Cerebellum | 1.276005E-03 |
| cytoskeleton organization | 7 | Human | Kidney | 9.689263E-05 |
| cytoskeleton organization | 7 | Human | Liver | 7.083989E-08 |
| cytoskeleton organization | 7 | Human | Ovary | 1.371051E-03 |
| cytoskeleton organization | 7 | Human | Testis | 1.122640E-03 |

| GO term | Number of species enriched | Species | Organ | p-value |
|---|---|---|---|---|
| cytoskeleton organization | 7 | Mouse | Cerebellum | 3.752323E-03 |
| cytoskeleton organization | 7 | Mouse | Liver | 4.666591E-02 |
| cytoskeleton organization | 7 | Mouse | Ovary | 2.064489E-03 |
| cytoskeleton organization | 7 | Mouse | Testis | 9.099250E-11 |
| cytoskeleton organization | 7 | Opossum | Kidney | 1.105362E-10 |
| cytoskeleton organization | 7 | Opossum | Liver | 3.761543E-07 |
| cytoskeleton organization | 7 | Opossum | Ovary | 6.778500E-05 |
| cytoskeleton organization | 7 | Opossum | Testis | 8.899299E-03 |
| cytoskeleton organization | 7 | Rabbit | Brain | 1.486343E-02 |
| cytoskeleton organization | 7 | Rabbit | Liver | 8.852884E-04 |
| cytoskeleton organization | 7 | Rabbit | Ovary | 3.016287E-02 |
| cytoskeleton organization | 7 | Rabbit | Testis | 3.744730E-04 |
| cytoskeleton organization | 7 | Rat | Brain | 2.763106E-02 |
| cytoskeleton organization | 7 | Rat | Cerebellum | 6.836958E-07 |
| cytoskeleton organization | 7 | Rat | Kidney | 3.048967E-07 |
| cytoskeleton organization | 7 | Rat | Testis | 6.120940E-04 |
| cytoskeleton organization | 7 | RhesusMacaque | Brain | 1.824785E-07 |
| cytoskeleton organization | 7 | RhesusMacaque | Heart | 1.973054E-04 |
| cytoskeleton organization | 7 | RhesusMacaque | Kidney | 5.775554E-07 |

| GO term | Number of species enriched | Species | Organ | p-value |
|---|---|---|---|---|
| cytoskeleton organization | 7 | RhesusMacaque | Liver | 3.284218E-03 |
| cytoskeleton organization | 7 | RhesusMacaque | Testis | 2.520300E-07 |
| cytoskeleton | 7 | Chicken | Brain | 3.034177E-03 |
| cytoskeleton | 7 | Chicken | Testis | 6.474395E-04 |
| cytoskeleton | 7 | Human | Brain | 5.982912E-05 |
| cytoskeleton | 7 | Human | Cerebellum | 1.599929E-02 |
| cytoskeleton | 7 | Human | Liver | 3.643901E-06 |
| cytoskeleton | 7 | Human | Testis | 2.233825E-02 |
| cytoskeleton | 7 | Mouse | Testis | 2.311793E-03 |
| cytoskeleton | 7 | Opossum | Kidney | 2.361224E-07 |
| cytoskeleton | 7 | Opossum | Liver | 4.698293E-03 |
| cytoskeleton | 7 | Rabbit | Liver | 1.968866E-02 |
| cytoskeleton | 7 | Rat | Brain | 2.094242E-04 |
| cytoskeleton | 7 | Rat | Cerebellum | 3.701705E-04 |
| cytoskeleton | 7 | Rat | Kidney | 1.095577E-03 |
| cytoskeleton | 7 | Rat | Testis | 3.266103E-02 |
| cytoskeleton | 7 | RhesusMacaque | Cerebellum | 1.146654E-03 |
| cytoskeleton | 7 | RhesusMacaque | Kidney | 1.068901E-03 |
| cytoskeleton | 7 | RhesusMacaque | Testis | 1.597053E-02 |
| centrosome | 7 | Chicken | Cerebellum | 1.959350E-05 |
| centrosome | 7 | Chicken | Kidney | 2.207138E-06 |
| centrosome | 7 | Chicken | Ovary | 3.868088E-05 |
| centrosome | 7 | Human | Brain | 2.484136E-10 |
| centrosome | 7 | Human | Liver | 6.169238E-04 |
| centrosome | 7 | Mouse | Cerebellum | 1.603534E-07 |
| centrosome | 7 | Mouse | Ovary | 2.021674E-07 |
| centrosome | 7 | Mouse | Testis | 3.041897E-06 |
| centrosome | 7 | Opossum | Kidney | 2.507759E-05 |

| GO term | Number of species enriched | Species | Organ | p-value |
|---|---|---|---|---|
| centrosome | 7 | Opossum | Liver | 1.145497E-07 |
| centrosome | 7 | Opossum | Ovary | 2.567070E-04 |
| centrosome | 7 | Rabbit | Brain | 1.327594E-04 |
| centrosome | 7 | Rabbit | Cerebellum | 2.871956E-04 |
| centrosome | 7 | Rabbit | Kidney | 2.542843E-02 |
| centrosome | 7 | Rabbit | Liver | 1.429747E-05 |
| centrosome | 7 | Rabbit | Ovary | 8.629719E-03 |
| centrosome | 7 | Rabbit | Testis | 4.846592E-07 |
| centrosome | 7 | Rat | Cerebellum | 6.532941E-05 |
| centrosome | 7 | Rat | Kidney | 2.683574E-06 |
| centrosome | 7 | Rat | Testis | 3.830566E-05 |
| centrosome | 7 | RhesusMacaque | Heart | 1.076976E-02 |
| centrosome | 7 | RhesusMacaque | Kidney | 4.531551E-02 |
| cellular lipid metabolic process | 7 | Chicken | Brain | 3.516599E-03 |
| cellular lipid metabolic process | 7 | Chicken | Heart | 2.727817E-02 |
| cellular lipid metabolic process | 7 | Chicken | Kidney | 4.300198E-03 |
| cellular lipid metabolic process | 7 | Human | Cerebellum | 2.503372E-05 |
| cellular lipid metabolic process | 7 | Mouse | Cerebellum | 8.442307E-03 |
| cellular lipid metabolic process | 7 | Mouse | Testis | 2.351080E-02 |
| cellular lipid metabolic process | 7 | Opossum | Liver | 1.214154E-03 |
| cellular lipid metabolic process | 7 | Opossum | Ovary | 1.130244E-03 |

| GO term | Number of species enriched | Species | Organ | p-value |
|---|---|---|---|---|
| cellular lipid metabolic process | 7 | Opossum | Testis | 2.651591E-02 |
| cellular lipid metabolic process | 7 | Rabbit | Kidney | 9.472469E-04 |
| cellular lipid metabolic process | 7 | Rabbit | Testis | 1.083784E-03 |
| cellular lipid metabolic process | 7 | Rat | Kidney | 1.583157E-02 |
| cellular lipid metabolic process | 7 | Rat | Liver | 4.760913E-06 |
| cellular lipid metabolic process | 7 | Rat | Ovary | 7.636012E-04 |
| cellular lipid metabolic process | 7 | RhesusMacaque | Cerebellum | 1.859533E-04 |
| cellular lipid metabolic process | 7 | RhesusMacaque | Liver | 1.271363E-05 |
| cell division | 7 | Chicken | Cerebellum | 1.278288E-12 |
| cell division | 7 | Chicken | Kidney | 1.755462E-08 |
| cell division | 7 | Human | Brain | 2.199814E-07 |
| cell division | 7 | Human | Cerebellum | 1.429781E-08 |
| cell division | 7 | Human | Ovary | 4.326172E-08 |
| cell division | 7 | Mouse | Brain | 3.407628E-03 |
| cell division | 7 | Mouse | Cerebellum | 8.300431E-09 |
| cell division | 7 | Mouse | Heart | 2.109375E-02 |
| cell division | 7 | Mouse | Ovary | 3.927316E-08 |
| cell division | 7 | Mouse | Testis | 4.751196E-05 |
| cell division | 7 | Opossum | Brain | 1.597163E-07 |
| cell division | 7 | Opossum | Heart | 1.017299E-02 |
| cell division | 7 | Opossum | Kidney | 1.216348E-04 |

| GO term | Number of species enriched | Species | Organ | p-value |
|---|---|---|---|---|
| cell division | 7 | Opossum | Liver | 1.095682E-02 |
| cell division | 7 | Rabbit | Brain | 4.278471E-07 |
| cell division | 7 | Rabbit | Cerebellum | 5.216762E-12 |
| cell division | 7 | Rabbit | Heart | 4.779129E-06 |
| cell division | 7 | Rabbit | Kidney | 1.148056E-06 |
| cell division | 7 | Rabbit | Ovary | 6.135173E-08 |
| cell division | 7 | Rabbit | Testis | 5.404434E-07 |
| cell division | 7 | Rat | Cerebellum | 6.204651E-07 |
| cell division | 7 | Rat | Kidney | 7.701412E-07 |
| cell division | 7 | Rat | Testis | 7.551913E-03 |
| cell division | 7 | RhesusMacaque | Cerebellum | 7.340968E-04 |
| cell cycle process | 7 | Chicken | Cerebellum | 1.064070E-19 |
| cell cycle process | 7 | Chicken | Heart | 5.049198E-04 |
| cell cycle process | 7 | Chicken | Kidney | 1.689374E-16 |
| cell cycle process | 7 | Chicken | Ovary | 3.522999E-08 |
| cell cycle process | 7 | Chicken | Testis | 2.068926E-04 |
| cell cycle process | 7 | Human | Brain | 5.661483E-18 |
| cell cycle process | 7 | Human | Cerebellum | 9.592378E-13 |
| cell cycle process | 7 | Human | Liver | 6.832185E-04 |
| cell cycle process | 7 | Human | Ovary | 3.380728E-13 |
| cell cycle process | 7 | Human | Testis | 4.468517E-03 |
| cell cycle process | 7 | Mouse | Brain | 3.353740E-11 |
| cell cycle process | 7 | Mouse | Cerebellum | 3.165613E-13 |

| GO term | Number of species enriched | Species | Organ | p-value |
|---|---|---|---|---|
| cell cycle process | 7 | Mouse | Heart | 6.248771E-07 |
| cell cycle process | 7 | Mouse | Ovary | 2.059616E-13 |
| cell cycle process | 7 | Mouse | Testis | 4.145799E-15 |
| cell cycle process | 7 | Opossum | Brain | 1.263279E-18 |
| cell cycle process | 7 | Opossum | Cerebellum | 2.370673E-05 |
| cell cycle process | 7 | Opossum | Heart | 3.149094E-05 |
| cell cycle process | 7 | Opossum | Kidney | 1.956145E-08 |
| cell cycle process | 7 | Opossum | Liver | 1.714714E-12 |
| cell cycle process | 7 | Opossum | Ovary | 8.084597E-04 |
| cell cycle process | 7 | Rabbit | Brain | 7.257045E-15 |
| cell cycle process | 7 | Rabbit | Cerebellum | 1.450520E-20 |
| cell cycle process | 7 | Rabbit | Heart | 1.861440E-14 |
| cell cycle process | 7 | Rabbit | Kidney | 6.408024E-13 |
| cell cycle process | 7 | Rabbit | Liver | 1.239731E-06 |
| cell cycle process | 7 | Rabbit | Ovary | 1.599195E-17 |
| cell cycle process | 7 | Rabbit | Testis | 1.741148E-13 |
| cell cycle process | 7 | Rat | Cerebellum | 1.277368E-17 |
| cell cycle process | 7 | Rat | Kidney | 5.465290E-12 |
| cell cycle process | 7 | Rat | Liver | 1.337306E-04 |

| GO term | Number of species enriched | Species | Organ | p-value |
|---|---|---|---|---|
| cell cycle process | 7 | Rat | Testis | 2.334480E-10 |
| cell cycle process | 7 | RhesusMacaque | Cerebellum | 3.975942E-07 |
| cell cycle process | 7 | RhesusMacaque | Heart | 1.586207E-04 |
| cell cycle process | 7 | RhesusMacaque | Testis | 1.210027E-04 |
| cell cycle | 7 | Chicken | Cerebellum | 2.329545E-24 |
| cell cycle | 7 | Chicken | Heart | 8.943694E-04 |
| cell cycle | 7 | Chicken | Kidney | 1.988646E-18 |
| cell cycle | 7 | Chicken | Ovary | 1.756320E-03 |
| cell cycle | 7 | Chicken | Testis | 5.597259E-07 |
| cell cycle | 7 | Human | Brain | 1.124982E-16 |
| cell cycle | 7 | Human | Cerebellum | 9.594305E-16 |
| cell cycle | 7 | Human | Liver | 1.325212E-03 |
| cell cycle | 7 | Human | Ovary | 2.091636E-16 |
| cell cycle | 7 | Human | Testis | 4.144964E-02 |
| cell cycle | 7 | Mouse | Brain | 4.862750E-15 |
| cell cycle | 7 | Mouse | Cerebellum | 3.639909E-18 |
| cell cycle | 7 | Mouse | Heart | 1.776477E-09 |
| cell cycle | 7 | Mouse | Ovary | 1.103343E-17 |
| cell cycle | 7 | Mouse | Testis | 1.737578E-13 |
| cell cycle | 7 | Opossum | Brain | 9.969578E-26 |
| cell cycle | 7 | Opossum | Cerebellum | 3.406712E-04 |
| cell cycle | 7 | Opossum | Heart | 2.531968E-02 |
| cell cycle | 7 | Opossum | Kidney | 5.676887E-05 |
| cell cycle | 7 | Opossum | Liver | 5.080961E-10 |
| cell cycle | 7 | Rabbit | Brain | 2.411341E-20 |
| cell cycle | 7 | Rabbit | Cerebellum | 4.528752E-27 |
| cell cycle | 7 | Rabbit | Heart | 5.612168E-16 |

| GO term | Number of species enriched | Species | Organ | p-value |
|---|---|---|---|---|
| cell cycle | 7 | Rabbit | Kidney | 2.381347E-11 |
| cell cycle | 7 | Rabbit | Liver | 2.364305E-10 |
| cell cycle | 7 | Rabbit | Ovary | 5.331695E-19 |
| cell cycle | 7 | Rabbit | Testis | 1.308708E-13 |
| cell cycle | 7 | Rat | Cerebellum | 2.525538E-19 |
| cell cycle | 7 | Rat | Kidney | 1.678135E-13 |
| cell cycle | 7 | Rat | Liver | 7.283382E-07 |
| cell cycle | 7 | Rat | Testis | 5.570600E-08 |
| cell cycle | 7 | RhesusMacaque | Cerebellum | 2.313066E-07 |
| cell cycle | 7 | RhesusMacaque | Testis | 5.531825E-03 |
| cell body | 7 | Chicken | Kidney | 2.059525E-03 |
| cell body | 7 | Human | Brain | 4.052754E-02 |
| cell body | 7 | Human | Liver | 2.652511E-04 |
| cell body | 7 | Human | Ovary | 4.719925E-02 |
| cell body | 7 | Mouse | Kidney | 8.882718E-03 |
| cell body | 7 | Opossum | Liver | 5.898215E-03 |
| cell body | 7 | Rabbit | Liver | 1.961514E-04 |
| cell body | 7 | Rabbit | Testis | 1.404049E-02 |
| cell body | 7 | Rat | Brain | 3.563311E-02 |
| cell body | 7 | Rat | Cerebellum | 1.651845E-02 |
| cell body | 7 | Rat | Testis | 4.064134E-02 |
| cell body | 7 | RhesusMacaque | Kidney | 1.571049E-04 |
| axon guidance | 7 | Chicken | Brain | 6.641830E-05 |
| axon guidance | 7 | Chicken | Kidney | 3.732914E-03 |
| axon guidance | 7 | Human | Brain | 3.453116E-03 |
| axon guidance | 7 | Human | Kidney | 3.018990E-02 |
| axon guidance | 7 | Human | Liver | 7.250320E-07 |
| axon guidance | 7 | Human | Ovary | 4.259498E-03 |
| axon guidance | 7 | Mouse | Ovary | 3.527938E-04 |

| GO term | Number of species enriched | Species | Organ | p-value |
|---|---|---|---|---|
| axon guidance | 7 | Opossum | Kidney | 1.731436E-05 |
| axon guidance | 7 | Opossum | Liver | 2.233841E-04 |
| axon guidance | 7 | Opossum | Ovary | 3.619472E-02 |
| axon guidance | 7 | Rabbit | Liver | 1.555297E-06 |
| axon guidance | 7 | Rat | Brain | 1.539105E-02 |
| axon guidance | 7 | Rat | Liver | 5.526978E-06 |
| axon guidance | 7 | RhesusMacaque | Brain | 6.310710E-05 |
| axon guidance | 7 | RhesusMacaque | Heart | 2.035647E-02 |
| axon guidance | 7 | RhesusMacaque | Kidney | 4.695783E-03 |
| Ras GTPase binding | 7 | Chicken | Kidney | 8.299257E-03 |
| Ras GTPase binding | 7 | Human | Brain | 3.905819E-06 |
| Ras GTPase binding | 7 | Human | Kidney | 1.318787E-04 |
| Ras GTPase binding | 7 | Human | Liver | 2.398545E-02 |
| Ras GTPase binding | 7 | Mouse | Ovary | 1.985583E-03 |
| Ras GTPase binding | 7 | Mouse | Testis | 4.204699E-06 |
| Ras GTPase binding | 7 | Opossum | Kidney | 1.072100E-05 |
| Ras GTPase binding | 7 | Opossum | Liver | 6.918023E-03 |
| Ras GTPase binding | 7 | Rabbit | Liver | 6.426747E-03 |
| Ras GTPase binding | 7 | Rat | Brain | 1.763914E-05 |
| Ras GTPase binding | 7 | Rat | Testis | 1.600511E-04 |
| Ras GTPase binding | 7 | RhesusMacaque | Testis | 1.719580E-03 |
| GTPase binding | 7 | Chicken | Kidney | 4.378600E-04 |

| GO term | Number of species enriched | Species | Organ | p-value |
|---|---|---|---|---|
| GTPase binding | 7 | Human | Brain | 9.456672E-09 |
| GTPase binding | 7 | Human | Cerebellum | 2.666969E-02 |
| GTPase binding | 7 | Human | Kidney | 3.814235E-05 |
| GTPase binding | 7 | Human | Liver | 1.506923E-02 |
| GTPase binding | 7 | Mouse | Ovary | 1.087442E-04 |
| GTPase binding | 7 | Mouse | Testis | 4.358373E-07 |
| GTPase binding | 7 | Opossum | Kidney | 4.923957E-07 |
| GTPase binding | 7 | Opossum | Liver | 5.345537E-03 |
| GTPase binding | 7 | Opossum | Ovary | 1.403758E-02 |
| GTPase binding | 7 | Rabbit | Brain | 4.778201E-02 |
| GTPase binding | 7 | Rabbit | Kidney | 2.971187E-02 |
| GTPase binding | 7 | Rabbit | Liver | 2.025186E-05 |
| GTPase binding | 7 | Rabbit | Testis | 1.695021E-02 |
| GTPase binding | 7 | Rat | Brain | 1.790889E-07 |
| GTPase binding | 7 | Rat | Testis | 2.247806E-06 |
| GTPase binding | 7 | RhesusMacaque | Heart | 8.634156E-03 |
| GTPase binding | 7 | RhesusMacaque | Testis | 4.694768E-04 |
| transferase complex | 6 | Chicken | Cerebellum | 2.715063E-02 |
| transferase complex | 6 | Chicken | Kidney | 7.244913E-04 |

| GO term | Number of species enriched | Species | Organ | p-value |
|---|---|---|---|---|
| transferase complex | 6 | Human | Brain | 1.408709E-05 |
| transferase complex | 6 | Human | Kidney | 5.091246E-03 |
| transferase complex | 6 | Human | Liver | 5.265119E-04 |
| transferase complex | 6 | Human | Ovary | 8.371257E-03 |
| transferase complex | 6 | Mouse | Cerebellum | 1.301526E-02 |
| transferase complex | 6 | Mouse | Liver | 2.994292E-02 |
| transferase complex | 6 | Mouse | Ovary | 2.199918E-02 |
| transferase complex | 6 | Opossum | Brain | 1.556568E-04 |
| transferase complex | 6 | Opossum | Kidney | 3.099497E-03 |
| transferase complex | 6 | Opossum | Liver | 5.088644E-08 |
| transferase complex | 6 | Opossum | Ovary | 4.729454E-02 |
| transferase complex | 6 | Rabbit | Kidney | 1.594661E-02 |
| transferase complex | 6 | Rabbit | Liver | 1.642330E-04 |
| transferase complex | 6 | Rabbit | Ovary | 5.906528E-04 |
| transferase complex | 6 | Rabbit | Testis | 2.163597E-03 |
| transferase complex | 6 | RhesusMacaque | Heart | 9.525127E-04 |
| transferase complex | 6 | RhesusMacaque | Testis | 8.352049E-04 |
| tRNA metabolic process | 6 | Chicken | Cerebellum | 1.092534E-02 |
| tRNA metabolic process | 6 | Human | Ovary | 4.003834E-03 |

| GO term | Number of species enriched | Species | Organ | p-value |
|---|---|---|---|---|
| tRNA metabolic process | 6 | Mouse | Brain | 3.313281E-08 |
| tRNA metabolic process | 6 | Mouse | Heart | 2.292828E-05 |
| tRNA metabolic process | 6 | Opossum | Brain | 2.210967E-04 |
| tRNA metabolic process | 6 | Opossum | Liver | 3.068610E-02 |
| tRNA metabolic process | 6 | Rabbit | Brain | 1.809936E-07 |
| tRNA metabolic process | 6 | Rabbit | Cerebellum | 6.635890E-03 |
| tRNA metabolic process | 6 | Rabbit | Heart | 5.179638E-06 |
| tRNA metabolic process | 6 | Rabbit | Liver | 3.817197E-02 |
| tRNA metabolic process | 6 | Rabbit | Ovary | 1.398576E-04 |
| tRNA metabolic process | 6 | Rat | Testis | 1.398075E-03 |
| regulation of mitotic cell cycle | 6 | Chicken | Cerebellum | 1.230454E-04 |
| regulation of mitotic cell cycle | 6 | Chicken | Kidney | 1.553512E-04 |
| regulation of mitotic cell cycle | 6 | Human | Brain | 1.026074E-04 |
| regulation of mitotic cell cycle | 6 | Human | Cerebellum | 1.352549E-03 |
| regulation of mitotic cell cycle | 6 | Human | Ovary | 1.863593E-02 |
| regulation of mitotic cell cycle | 6 | Mouse | Brain | 1.038809E-07 |

| GO term | Number of species enriched | Species | Organ | p-value |
|---|---|---|---|---|
| regulation of mitotic cell cycle | 6 | Mouse | Cerebellum | 2.934121E-04 |
| regulation of mitotic cell cycle | 6 | Mouse | Ovary | 1.749678E-03 |
| regulation of mitotic cell cycle | 6 | Mouse | Testis | 1.631839E-04 |
| regulation of mitotic cell cycle | 6 | Opossum | Brain | 7.842998E-05 |
| regulation of mitotic cell cycle | 6 | Opossum | Liver | 5.425916E-04 |
| regulation of mitotic cell cycle | 6 | Rabbit | Brain | 2.301954E-04 |
| regulation of mitotic cell cycle | 6 | Rabbit | Cerebellum | 1.304063E-08 |
| regulation of mitotic cell cycle | 6 | Rabbit | Heart | 2.007866E-02 |
| regulation of mitotic cell cycle | 6 | Rabbit | Kidney | 8.947821E-03 |
| regulation of mitotic cell cycle | 6 | Rabbit | Ovary | 2.628465E-04 |
| regulation of mitotic cell cycle | 6 | Rabbit | Testis | 1.920454E-03 |
| regulation of mitotic cell cycle | 6 | Rat | Cerebellum | 8.201162E-06 |
| regulation of mitotic cell cycle | 6 | Rat | Kidney | 1.268278E-04 |
| regulation of mitotic cell cycle | 6 | Rat | Testis | 1.639912E-03 |

| GO term | Number of species enriched | Species | Organ | p-value |
|---|---|---|---|---|
| regulation of locomotion | 6 | Chicken | Brain | 1.083404E-04 |
| regulation of locomotion | 6 | Human | Brain | 3.253056E-02 |
| regulation of locomotion | 6 | Human | Kidney | 4.761415E-03 |
| regulation of locomotion | 6 | Mouse | Testis | 9.015299E-03 |
| regulation of locomotion | 6 | Opossum | Kidney | 8.720790E-04 |
| regulation of locomotion | 6 | Opossum | Testis | 3.303526E-02 |
| regulation of locomotion | 6 | Rat | Brain | 2.810669E-03 |
| regulation of locomotion | 6 | RhesusMacaque | Heart | 1.014160E-03 |
| regulation of locomotion | 6 | RhesusMacaque | Kidney | 4.597168E-03 |
| regulation of locomotion | 6 | RhesusMacaque | Liver | 7.082156E-03 |
| regulation of cellular response to heat | 6 | Chicken | Cerebellum | 7.785325E-03 |
| regulation of cellular response to heat | 6 | Chicken | Kidney | 2.289092E-02 |
| regulation of cellular response to heat | 6 | Human | Liver | 2.615858E-03 |
| regulation of cellular response to heat | 6 | Human | Testis | 3.947530E-02 |
| regulation of cellular response to heat | 6 | Mouse | Brain | 3.326471E-03 |

| GO term | Number of species enriched | Species | Organ | p-value |
|---|---|---|---|---|
| regulation of cellular response to heat | 6 | Mouse | Heart | 3.666393E-03 |
| regulation of cellular response to heat | 6 | Opossum | Brain | 2.769424E-04 |
| regulation of cellular response to heat | 6 | Rabbit | Brain | 1.168526E-04 |
| regulation of cellular response to heat | 6 | Rabbit | Cerebellum | 2.358258E-02 |
| regulation of cellular response to heat | 6 | Rabbit | Heart | 1.642274E-04 |
| regulation of cellular response to heat | 6 | Rabbit | Liver | 1.493291E-03 |
| regulation of cellular response to heat | 6 | Rabbit | Testis | 5.121612E-03 |
| regulation of cellular response to heat | 6 | Rat | Testis | 6.312333E-04 |
| regulation of cellular component movement | 6 | Chicken | Brain | 9.862214E-06 |
| regulation of cellular component movement | 6 | Human | Brain | 1.455935E-02 |
| regulation of cellular component movement | 6 | Human | Cerebellum | 2.635160E-02 |

| GO term | Number of species enriched | Species | Organ | p-value |
|---|---|---|---|---|
| regulation of cellular component movement | 6 | Human | Kidney | 2.053174E-03 |
| regulation of cellular component movement | 6 | Human | Testis | 2.927668E-02 |
| regulation of cellular component movement | 6 | Mouse | Kidney | 4.887697E-02 |
| regulation of cellular component movement | 6 | Mouse | Testis | 1.029096E-02 |
| regulation of cellular component movement | 6 | Opossum | Kidney | 6.196964E-04 |
| regulation of cellular component movement | 6 | Opossum | Ovary | 2.793042E-02 |
| regulation of cellular component movement | 6 | Opossum | Testis | 8.918589E-03 |
| regulation of cellular component movement | 6 | Rat | Brain | 3.397238E-03 |
| regulation of cellular component movement | 6 | RhesusMacaque | Heart | 4.121422E-05 |
| regulation of cellular component movement | 6 | RhesusMacaque | Kidney | 9.010429E-03 |
| regulation of cellular component movement | 6 | RhesusMacaque | Liver | 3.082656E-03 |

| GO term | Number of species enriched | Species | Organ | p-value |
|---|---|---|---|---|
| regulation of cell motility | 6 | Chicken | Brain | 4.208354E-05 |
| regulation of cell motility | 6 | Human | Brain | 2.944178E-02 |
| regulation of cell motility | 6 | Human | Kidney | 1.305358E-02 |
| regulation of cell motility | 6 | Mouse | Testis | 3.020670E-03 |
| regulation of cell motility | 6 | Opossum | Kidney | 9.561096E-04 |
| regulation of cell motility | 6 | Opossum | Testis | 2.754169E-02 |
| regulation of cell motility | 6 | Rat | Brain | 1.780797E-03 |
| regulation of cell motility | 6 | RhesusMacaque | Heart | 2.751715E-04 |
| regulation of cell motility | 6 | RhesusMacaque | Kidney | 4.352587E-03 |
| regulation of cell motility | 6 | RhesusMacaque | Liver | 6.739199E-03 |
| regulation of cell migration | 6 | Chicken | Brain | 8.446238E-05 |
| regulation of cell migration | 6 | Human | Kidney | 1.859370E-02 |
| regulation of cell migration | 6 | Mouse | Testis | 3.410088E-02 |
| regulation of cell migration | 6 | Opossum | Kidney | 1.476327E-03 |
| regulation of cell migration | 6 | Rat | Brain | 7.881653E-03 |
| regulation of cell migration | 6 | RhesusMacaque | Heart | 5.225279E-04 |
| regulation of cell migration | 6 | RhesusMacaque | Kidney | 2.801282E-02 |
| regulation of cell migration | 6 | RhesusMacaque | Liver | 1.097711E-02 |
| regulation of cell cycle process | 6 | Chicken | Cerebellum | 6.896947E-04 |

| GO term | Number of species enriched | Species | Organ | p-value |
|---|---|---|---|---|
| regulation of cell cycle process | 6 | Chicken | Kidney | 2.763064E-02 |
| regulation of cell cycle process | 6 | Human | Brain | 2.824499E-02 |
| regulation of cell cycle process | 6 | Mouse | Brain | 6.533891E-04 |
| regulation of cell cycle process | 6 | Mouse | Cerebellum | 7.901881E-04 |
| regulation of cell cycle process | 6 | Opossum | Brain | 2.079002E-05 |
| regulation of cell cycle process | 6 | Rabbit | Brain | 6.527450E-05 |
| regulation of cell cycle process | 6 | Rabbit | Cerebellum | 1.828767E-07 |
| regulation of cell cycle process | 6 | Rabbit | Heart | 2.794359E-03 |
| regulation of cell cycle process | 6 | Rabbit | Ovary | 1.229418E-03 |
| regulation of cell cycle process | 6 | Rat | Cerebellum | 2.120861E-03 |
| protein kinase binding | 6 | Human | Brain | 1.335230E-03 |
| protein kinase binding | 6 | Mouse | Cerebellum | 1.760900E-02 |
| protein kinase binding | 6 | Opossum | Kidney | 5.246481E-05 |
| protein kinase binding | 6 | Opossum | Liver | 4.453792E-02 |
| protein kinase binding | 6 | Opossum | Ovary | 1.586847E-03 |

| GO term | Number of species enriched | Species | Organ | p-value |
|---|---|---|---|---|
| protein kinase binding | 6 | Rabbit | Testis | 1.076373E-02 |
| protein kinase binding | 6 | Rat | Kidney | 1.229594E-02 |
| protein kinase binding | 6 | RhesusMacaque | Brain | 4.212714E-03 |
| post-translational protein modification | 6 | Chicken | Kidney | 7.243600E-03 |
| post-translational protein modification | 6 | Human | Brain | 6.472678E-04 |
| post-translational protein modification | 6 | Human | Ovary | 7.953445E-03 |
| post-translational protein modification | 6 | Mouse | Cerebellum | 9.029298E-03 |
| post-translational protein modification | 6 | Mouse | Heart | 9.969219E-04 |
| post-translational protein modification | 6 | Opossum | Liver | 9.345584E-05 |
| post-translational protein modification | 6 | Opossum | Ovary | 4.388801E-03 |
| post-translational protein modification | 6 | Rabbit | Brain | 5.626926E-03 |
| post-translational protein modification | 6 | Rabbit | Heart | 1.832680E-03 |

| GO term | Number of species enriched | Species | Organ | p-value |
|---|---|---|---|---|
| post-translational protein modification | 6 | Rabbit | Liver | 1.704011E-02 |
| post-translational protein modification | 6 | Rat | Liver | 6.901295E-03 |
| positive regulation of kinase activity | 6 | Chicken | Brain | 3.802221E-06 |
| positive regulation of kinase activity | 6 | Human | Kidney | 1.218164E-02 |
| positive regulation of kinase activity | 6 | Mouse | Kidney | 1.672076E-02 |
| positive regulation of kinase activity | 6 | Mouse | Ovary | 4.836442E-02 |
| positive regulation of kinase activity | 6 | Mouse | Testis | 1.458973E-02 |
| positive regulation of kinase activity | 6 | Opossum | Kidney | 1.754719E-03 |
| positive regulation of kinase activity | 6 | Rat | Liver | 1.159716E-02 |
| positive regulation of kinase activity | 6 | RhesusMacaque | Brain | 2.464865E-02 |
| positive regulation of hydrolase activity | 6 | Chicken | Brain | 6.925917E-03 |
| positive regulation of hydrolase activity | 6 | Human | Brain | 3.325736E-02 |

| GO term | Number of species enriched | Species | Organ | p-value |
|---|---|---|---|---|
| positive regulation of hydrolase activity | 6 | Mouse | Testis | 2.046811E-02 |
| positive regulation of hydrolase activity | 6 | Opossum | Kidney | 1.059429E-03 |
| positive regulation of hydrolase activity | 6 | Rat | Brain | 5.046554E-05 |
| positive regulation of hydrolase activity | 6 | RhesusMacaque | Liver | 3.753607E-02 |
| phosphatidylinositol binding | 6 | Chicken | Brain | 1.291629E-03 |
| phosphatidylinositol binding | 6 | Chicken | Kidney | 1.741545E-02 |
| phosphatidylinositol binding | 6 | Human | Brain | 2.450211E-02 |
| phosphatidylinositol binding | 6 | Opossum | Kidney | 1.083008E-03 |
| phosphatidylinositol binding | 6 | Opossum | Liver | 3.456872E-05 |
| phosphatidylinositol binding | 6 | Opossum | Ovary | 1.771661E-03 |
| phosphatidylinositol binding | 6 | Opossum | Testis | 2.887722E-03 |
| phosphatidylinositol binding | 6 | Rabbit | Liver | 3.291813E-02 |
| phosphatidylinositol binding | 6 | Rat | Brain | 3.858046E-03 |
| phosphatidylinositol binding | 6 | RhesusMacaque | Heart | 1.452566E-02 |
| phosphatidylinositol binding | 6 | RhesusMacaque | Liver | 2.360068E-02 |
| peptidyl-lysine modification | 6 | Chicken | Cerebellum | 2.044918E-03 |

| GO term | Number of species enriched | Species | Organ | p-value |
|---|---|---|---|---|
| peptidyl-lysine modification | 6 | Chicken | Kidney | 1.756840E-03 |
| peptidyl-lysine modification | 6 | Human | Liver | 1.244497E-03 |
| peptidyl-lysine modification | 6 | Mouse | Cerebellum | 1.405675E-02 |
| peptidyl-lysine modification | 6 | Opossum | Brain | 1.092505E-04 |
| peptidyl-lysine modification | 6 | Opossum | Liver | 4.969648E-03 |
| peptidyl-lysine modification | 6 | Rabbit | Cerebellum | 6.579179E-05 |
| peptidyl-lysine modification | 6 | Rabbit | Ovary | 5.581866E-04 |
| peptidyl-lysine modification | 6 | Rat | Cerebellum | 1.538434E-02 |
| peptidyl-lysine modification | 6 | Rat | Testis | 2.438604E-02 |
| nucleotide metabolic process | 6 | Chicken | Kidney | 3.543961E-02 |
| nucleotide metabolic process | 6 | Human | Cerebellum | 4.972711E-03 |
| nucleotide metabolic process | 6 | Human | Ovary | 7.487364E-04 |
| nucleotide metabolic process | 6 | Mouse | Brain | 1.035163E-03 |
| nucleotide metabolic process | 6 | Opossum | Brain | 2.115121E-02 |
| nucleotide metabolic process | 6 | Opossum | Liver | 1.636568E-02 |
| nucleotide metabolic process | 6 | Rabbit | Brain | 9.565703E-04 |

| GO term | Number of species enriched | Species | Organ | p-value |
|---|---|---|---|---|
| nucleotide metabolic process | 6 | Rabbit | Heart | 3.958944E-02 |
| nucleotide metabolic process | 6 | Rabbit | Kidney | 3.134592E-03 |
| nucleotide metabolic process | 6 | Rabbit | Testis | 3.844067E-03 |
| nucleotide metabolic process | 6 | Rat | Liver | 9.997461E-06 |
| nucleobase-containing small molecule metabolic process | 6 | Chicken | Kidney | 4.022341E-02 |
| nucleobase-containing small molecule metabolic process | 6 | Human | Cerebellum | 3.337147E-03 |
| nucleobase-containing small molecule metabolic process | 6 | Human | Ovary | 6.254476E-04 |
| nucleobase-containing small molecule metabolic process | 6 | Mouse | Brain | 2.029045E-02 |
| nucleobase-containing small molecule metabolic process | 6 | Opossum | Liver | 5.310242E-03 |
| nucleobase-containing small molecule metabolic process | 6 | Rabbit | Brain | 3.484008E-03 |

| GO term | Number of species enriched | Species | Organ | p-value |
|---|---|---|---|---|
| nucleobase-containing small molecule metabolic process | 6 | Rabbit | Heart | 2.429437E-02 |
| nucleobase-containing small molecule metabolic process | 6 | Rabbit | Kidney | 6.556935E-03 |
| nucleobase-containing small molecule metabolic process | 6 | Rabbit | Testis | 4.356465E-03 |
| nucleobase-containing small molecule metabolic process | 6 | Rat | Liver | 4.317301E-06 |
| ncRNA metabolic process | 6 | Chicken | Cerebellum | 7.301979E-08 |
| ncRNA metabolic process | 6 | Chicken | Testis | 5.092375E-03 |
| ncRNA metabolic process | 6 | Human | Ovary | 2.807464E-03 |
| ncRNA metabolic process | 6 | Mouse | Brain | 2.034433E-09 |
| ncRNA metabolic process | 6 | Mouse | Heart | 3.245384E-08 |
| ncRNA metabolic process | 6 | Opossum | Brain | 1.723023E-05 |
| ncRNA metabolic process | 6 | Rabbit | Brain | 1.250117E-07 |
| ncRNA metabolic process | 6 | Rabbit | Cerebellum | 1.137051E-04 |

| GO term | Number of species enriched | Species | Organ | p-value |
|---|---|---|---|---|
| ncRNA metabolic process | 6 | Rabbit | Heart | 3.090677E-07 |
| ncRNA metabolic process | 6 | Rabbit | Ovary | 2.418498E-05 |
| ncRNA metabolic process | 6 | Rat | Testis | 2.313795E-05 |
| mitotic cell cycle phase transition | 6 | Chicken | Cerebellum | 7.447771E-07 |
| mitotic cell cycle phase transition | 6 | Chicken | Kidney | 1.030291E-07 |
| mitotic cell cycle phase transition | 6 | Human | Brain | 1.057672E-07 |
| mitotic cell cycle phase transition | 6 | Human | Cerebellum | 1.201248E-04 |
| mitotic cell cycle phase transition | 6 | Human | Ovary | 2.739810E-07 |
| mitotic cell cycle phase transition | 6 | Mouse | Brain | 3.317057E-06 |
| mitotic cell cycle phase transition | 6 | Mouse | Cerebellum | 7.712239E-07 |
| mitotic cell cycle phase transition | 6 | Mouse | Heart | 2.467420E-05 |
| mitotic cell cycle phase transition | 6 | Mouse | Ovary | 8.904303E-06 |
| mitotic cell cycle phase transition | 6 | Mouse | Testis | 3.028419E-04 |
| mitotic cell cycle phase transition | 6 | Opossum | Brain | 2.726870E-10 |

| GO term | Number of species enriched | Species | Organ | p-value |
|---|---|---|---|---|
| mitotic cell cycle phase transition | 6 | Opossum | Cerebellum | 1.562430E-03 |
| mitotic cell cycle phase transition | 6 | Opossum | Liver | 5.499623E-05 |
| mitotic cell cycle phase transition | 6 | Rabbit | Brain | 2.943837E-07 |
| mitotic cell cycle phase transition | 6 | Rabbit | Cerebellum | 2.551515E-09 |
| mitotic cell cycle phase transition | 6 | Rabbit | Heart | 1.890115E-05 |
| mitotic cell cycle phase transition | 6 | Rabbit | Kidney | 5.891593E-05 |
| mitotic cell cycle phase transition | 6 | Rabbit | Liver | 3.573334E-02 |
| mitotic cell cycle phase transition | 6 | Rabbit | Ovary | 1.794327E-10 |
| mitotic cell cycle phase transition | 6 | Rabbit | Testis | 3.119594E-04 |
| mitotic cell cycle phase transition | 6 | Rat | Cerebellum | 2.631097E-09 |
| mitotic cell cycle phase transition | 6 | Rat | Kidney | 2.462566E-03 |
| mitotic cell cycle phase transition | 6 | Rat | Liver | 2.607119E-03 |
| mitotic cell cycle phase transition | 6 | Rat | Testis | 7.210522E-03 |
| microtubule-based process | 6 | Chicken | Cerebellum | 7.828171E-03 |

| GO term | Number of species enriched | Species | Organ | p-value |
|---|---|---|---|---|
| microtubule-based process | 6 | Chicken | Kidney | 1.430412E-02 |
| microtubule-based process | 6 | Chicken | Ovary | 6.715058E-04 |
| microtubule-based process | 6 | Chicken | Testis | 1.653247E-04 |
| microtubule-based process | 6 | Human | Brain | 7.846525E-08 |
| microtubule-based process | 6 | Human | Liver | 5.148246E-05 |
| microtubule-based process | 6 | Human | Ovary | 5.546171E-03 |
| microtubule-based process | 6 | Mouse | Cerebellum | 7.538854E-04 |
| microtubule-based process | 6 | Mouse | Ovary | 2.358177E-02 |
| microtubule-based process | 6 | Mouse | Testis | 4.226573E-06 |
| microtubule-based process | 6 | Opossum | Kidney | 5.590922E-05 |
| microtubule-based process | 6 | Opossum | Liver | 9.166214E-05 |
| microtubule-based process | 6 | Rabbit | Brain | 7.636552E-05 |
| microtubule-based process | 6 | Rabbit | Liver | 1.837846E-02 |
| microtubule-based process | 6 | Rabbit | Testis | 5.032929E-04 |
| microtubule-based process | 6 | Rat | Cerebellum | 2.128602E-03 |
| microtubule-based process | 6 | Rat | Kidney | 1.315852E-02 |
| microtubule cytoskeleton organization | 6 | Chicken | Cerebellum | 7.801272E-03 |
| microtubule cytoskeleton organization | 6 | Human | Brain | 1.664808E-04 |

| GO term | Number of species enriched | Species | Organ | p-value |
|---|---|---|---|---|
| microtubule cytoskeleton organization | 6 | Human | Liver | 1.485774E-02 |
| microtubule cytoskeleton organization | 6 | Mouse | Testis | 1.545140E-04 |
| microtubule cytoskeleton organization | 6 | Opossum | Kidney | 6.003246E-03 |
| microtubule cytoskeleton organization | 6 | Opossum | Liver | 1.666864E-02 |
| microtubule cytoskeleton organization | 6 | Rabbit | Brain | 4.856503E-04 |
| microtubule cytoskeleton organization | 6 | Rat | Kidney | 2.887443E-02 |
| ligase activity | 6 | Human | Brain | 1.718982E-04 |
| ligase activity | 6 | Human | Kidney | 2.854556E-02 |
| ligase activity | 6 | Human | Ovary | 5.173059E-04 |
| ligase activity | 6 | Mouse | Cerebellum | 3.568146E-02 |
| ligase activity | 6 | Mouse | Ovary | 9.100439E-03 |
| ligase activity | 6 | Mouse | Testis | 3.209191E-05 |
| ligase activity | 6 | Opossum | Liver | 1.751533E-03 |
| ligase activity | 6 | Opossum | Ovary | 9.287930E-03 |
| ligase activity | 6 | Rabbit | Brain | 7.051953E-03 |
| ligase activity | 6 | Rabbit | Heart | 3.031081E-03 |
| ligase activity | 6 | Rabbit | Ovary | 1.209737E-02 |
| ligase activity | 6 | Rabbit | Testis | 2.442208E-04 |
| ligase activity | 6 | Rat | Testis | 1.802774E-03 |
| ligase activity | 6 | RhesusMacaque | Testis | 1.387290E-02 |
| kinase binding | 6 | Human | Brain | 2.181882E-04 |
| kinase binding | 6 | Mouse | Cerebellum | 8.546326E-03 |
| kinase binding | 6 | Opossum | Kidney | 1.279839E-05 |

| GO term | Number of species enriched | Species | Organ | p-value |
|---|---|---|---|---|
| kinase binding | 6 | Opossum | Liver | 3.255335E-02 |
| kinase binding | 6 | Opossum | Ovary | 4.530825E-04 |
| kinase binding | 6 | Rabbit | Testis | 1.072742E-03 |
| kinase binding | 6 | Rat | Kidney | 3.681912E-03 |
| kinase binding | 6 | RhesusMacaque | Brain | 2.985274E-04 |
| extrinsic component of membrane | 6 | Chicken | Brain | 3.374748E-03 |
| extrinsic component of membrane | 6 | Human | Brain | 1.898846E-05 |
| extrinsic component of membrane | 6 | Human | Kidney | 1.732436E-02 |
| extrinsic component of membrane | 6 | Human | Testis | 2.439427E-02 |
| extrinsic component of membrane | 6 | Opossum | Kidney | 1.756318E-06 |
| extrinsic component of membrane | 6 | Opossum | Liver | 5.276464E-03 |
| extrinsic component of membrane | 6 | Rabbit | Kidney | 4.829025E-02 |
| extrinsic component of membrane | 6 | Rabbit | Testis | 3.220397E-03 |
| extrinsic component of membrane | 6 | Rat | Brain | 1.967000E-02 |
| extrinsic component of membrane | 6 | Rat | Liver | 1.821765E-03 |
| extrinsic component of membrane | 6 | RhesusMacaque | Kidney | 2.175604E-02 |

| GO term | Number of species enriched | Species | Organ | p-value |
|---|---|---|---|---|
| extrinsic component of membrane | 6 | RhesusMacaque | Liver | 3.502370E-03 |
| extrinsic component of membrane | 6 | RhesusMacaque | Testis | 5.896510E-04 |
| developmental growth | 6 | Chicken | Brain | 2.982929E-02 |
| developmental growth | 6 | Chicken | Kidney | 2.370872E-02 |
| developmental growth | 6 | Human | Testis | 2.055683E-03 |
| developmental growth | 6 | Opossum | Kidney | 1.385888E-03 |
| developmental growth | 6 | Opossum | Liver | 1.495771E-03 |
| developmental growth | 6 | Opossum | Testis | 4.016085E-02 |
| developmental growth | 6 | Rabbit | Testis | 3.367075E-02 |
| developmental growth | 6 | Rat | Kidney | 3.248649E-02 |
| developmental growth | 6 | RhesusMacaque | Kidney | 3.203579E-02 |
| developmental growth | 6 | RhesusMacaque | Liver | 1.649011E-02 |
| condensed chromosome kinetochore | 6 | Chicken | Cerebellum | 2.013938E-03 |
| condensed chromosome kinetochore | 6 | Chicken | Kidney | 9.286761E-03 |
| condensed chromosome kinetochore | 6 | Human | Cerebellum | 2.222360E-02 |
| condensed chromosome kinetochore | 6 | Human | Ovary | 2.206545E-03 |

| GO term | Number of species enriched | Species | Organ | p-value |
|---|---|---|---|---|
| condensed chromosome kinetochore | 6 | Mouse | Cerebellum | 1.347752E-04 |
| condensed chromosome kinetochore | 6 | Opossum | Brain | 6.028149E-05 |
| condensed chromosome kinetochore | 6 | Rabbit | Brain | 3.623035E-05 |
| condensed chromosome kinetochore | 6 | Rabbit | Cerebellum | 4.847762E-04 |
| condensed chromosome kinetochore | 6 | Rabbit | Heart | 4.583003E-02 |
| condensed chromosome kinetochore | 6 | Rabbit | Ovary | 4.475512E-03 |
| condensed chromosome kinetochore | 6 | Rat | Cerebellum | 9.281708E-05 |
| condensed chromosome kinetochore | 6 | Rat | Kidney | 1.176427E-03 |
| ciliary tip | 6 | Chicken | Ovary | 3.875133E-05 |
| ciliary tip | 6 | Human | Brain | 1.539999E-02 |
| ciliary tip | 6 | Human | Liver | 1.910235E-02 |
| ciliary tip | 6 | Mouse | Cerebellum | 9.149745E-03 |
| ciliary tip | 6 | Opossum | Liver | 6.556481E-04 |
| ciliary tip | 6 | Rabbit | Liver | 8.401799E-04 |
| ciliary tip | 6 | Rat | Cerebellum | 3.669940E-02 |
| chromosome organization | 6 | Chicken | Cerebellum | 6.395679E-13 |
| chromosome organization | 6 | Chicken | Kidney | 1.070700E-04 |
| chromosome organization | 6 | Chicken | Ovary | 1.761649E-02 |

| GO term | Number of species enriched | Species | Organ | p-value |
|---|---|---|---|---|
| chromosome organization | 6 | Human | Brain | 1.728065E-04 |
| chromosome organization | 6 | Human | Cerebellum | 9.169969E-06 |
| chromosome organization | 6 | Human | Ovary | 7.912271E-08 |
| chromosome organization | 6 | Mouse | Brain | 2.995115E-02 |
| chromosome organization | 6 | Mouse | Cerebellum | 2.112327E-05 |
| chromosome organization | 6 | Mouse | Ovary | 8.738855E-03 |
| chromosome organization | 6 | Opossum | Brain | 2.364454E-10 |
| chromosome organization | 6 | Rabbit | Brain | 6.790481E-06 |
| chromosome organization | 6 | Rabbit | Cerebellum | 3.545740E-06 |
| chromosome organization | 6 | Rabbit | Heart | 8.776488E-06 |
| chromosome organization | 6 | Rabbit | Ovary | 2.557506E-06 |
| chromosome organization | 6 | Rabbit | Testis | 3.925063E-04 |
| chromosome organization | 6 | Rat | Cerebellum | 7.324681E-07 |
| chromosome organization | 6 | Rat | Kidney | 1.345211E-03 |
| chromosome organization | 6 | Rat | Testis | 1.850181E-02 |
| chromatin binding | 6 | Chicken | Cerebellum | 4.827048E-05 |
| chromatin binding | 6 | Chicken | Kidney | 2.589422E-02 |
| chromatin binding | 6 | Human | Liver | 1.178949E-02 |
| chromatin binding | 6 | Mouse | Cerebellum | 2.460414E-04 |

| GO term | Number of species enriched | Species | Organ | p-value |
|---|---|---|---|---|
| chromatin binding | 6 | Mouse | Heart | 4.542242E-02 |
| chromatin binding | 6 | Opossum | Brain | 4.296623E-02 |
| chromatin binding | 6 | Opossum | Kidney | 3.523288E-02 |
| chromatin binding | 6 | Rabbit | Cerebellum | 7.261572E-07 |
| chromatin binding | 6 | Rabbit | Ovary | 3.105181E-03 |
| chromatin binding | 6 | Rat | Cerebellum | 1.685445E-02 |
| cellular response to DNA damage stimulus | 6 | Chicken | Cerebellum | 2.766467E-11 |
| cellular response to DNA damage stimulus | 6 | Chicken | Kidney | 1.383647E-05 |
| cellular response to DNA damage stimulus | 6 | Human | Brain | 8.957823E-08 |
| cellular response to DNA damage stimulus | 6 | Human | Cerebellum | 1.583507E-03 |
| cellular response to DNA damage stimulus | 6 | Human | Ovary | 1.168626E-11 |
| cellular response to DNA damage stimulus | 6 | Human | Testis | 1.345098E-02 |
| cellular response to DNA damage stimulus | 6 | Mouse | Brain | 1.849922E-07 |

| GO term | Number of species enriched | Species | Organ | p-value |
|---|---|---|---|---|
| cellular response to DNA damage stimulus | 6 | Mouse | Cerebellum | 5.247329E-10 |
| cellular response to DNA damage stimulus | 6 | Mouse | Heart | 1.086874E-02 |
| cellular response to DNA damage stimulus | 6 | Mouse | Ovary | 4.009340E-06 |
| cellular response to DNA damage stimulus | 6 | Mouse | Testis | 1.849912E-02 |
| cellular response to DNA damage stimulus | 6 | Opossum | Brain | 2.025070E-12 |
| cellular response to DNA damage stimulus | 6 | Opossum | Liver | 9.211465E-06 |
| cellular response to DNA damage stimulus | 6 | Rabbit | Brain | 5.647658E-06 |
| cellular response to DNA damage stimulus | 6 | Rabbit | Cerebellum | 6.263268E-12 |
| cellular response to DNA damage stimulus | 6 | Rabbit | Heart | 2.439977E-07 |
| cellular response to DNA damage stimulus | 6 | Rabbit | Liver | 1.477793E-06 |
| cellular response to DNA damage stimulus | 6 | Rabbit | Ovary | 1.007572E-10 |

| GO term | Number of species enriched | Species | Organ | p-value |
|---|---|---|---|---|
| cellular response to DNA damage stimulus | 6 | Rabbit | Testis | 3.897514E-05 |
| cellular response to DNA damage stimulus | 6 | Rat | Cerebellum | 2.168173E-09 |
| cellular response to DNA damage stimulus | 6 | Rat | Testis | 1.762937E-05 |
| cell projection organization | 6 | Human | Brain | 1.857124E-02 |
| cell projection organization | 6 | Human | Liver | 1.408987E-05 |
| cell projection organization | 6 | Mouse | Ovary | 2.108047E-02 |
| cell projection organization | 6 | Opossum | Kidney | 1.722166E-03 |
| cell projection organization | 6 | Rabbit | Liver | 1.766166E-05 |
| cell projection organization | 6 | Rat | Brain | 9.380442E-04 |
| cell projection organization | 6 | Rat | Cerebellum | 2.853307E-02 |
| cell projection organization | 6 | RhesusMacaque | Kidney | 3.142852E-04 |
| cell cycle phase transition | 6 | Chicken | Cerebellum | 2.016098E-07 |
| cell cycle phase transition | 6 | Chicken | Kidney | 9.415487E-08 |
| cell cycle phase transition | 6 | Human | Brain | 4.756674E-08 |
| cell cycle phase transition | 6 | Human | Cerebellum | 4.661351E-05 |

| GO term | Number of species enriched | Species | Organ | p-value |
|---|---|---|---|---|
| cell cycle phase transition | 6 | Human | Ovary | 1.931856E-07 |
| cell cycle phase transition | 6 | Mouse | Brain | 1.135883E-06 |
| cell cycle phase transition | 6 | Mouse | Cerebellum | 6.686629E-07 |
| cell cycle phase transition | 6 | Mouse | Heart | 1.620913E-05 |
| cell cycle phase transition | 6 | Mouse | Ovary | 3.357009E-06 |
| cell cycle phase transition | 6 | Mouse | Testis | 2.641101E-04 |
| cell cycle phase transition | 6 | Opossum | Brain | 1.883552E-10 |
| cell cycle phase transition | 6 | Opossum | Cerebellum | 8.872915E-04 |
| cell cycle phase transition | 6 | Opossum | Liver | 5.773640E-05 |
| cell cycle phase transition | 6 | Rabbit | Brain | 2.376336E-07 |
| cell cycle phase transition | 6 | Rabbit | Cerebellum | 7.163837E-10 |
| cell cycle phase transition | 6 | Rabbit | Heart | 1.471722E-05 |
| cell cycle phase transition | 6 | Rabbit | Kidney | 2.262932E-05 |
| cell cycle phase transition | 6 | Rabbit | Liver | 3.134994E-02 |

| GO term | Number of species enriched | Species | Organ | p-value |
|---|---|---|---|---|
| cell cycle phase transition | 6 | Rabbit | Ovary | 5.401810E-11 |
| cell cycle phase transition | 6 | Rabbit | Testis | 2.950759E-04 |
| cell cycle phase transition | 6 | Rat | Cerebellum | 8.250004E-10 |
| cell cycle phase transition | 6 | Rat | Kidney | 2.003698E-03 |
| cell cycle phase transition | 6 | Rat | Liver | 1.056415E-03 |
| cell cycle phase transition | 6 | Rat | Testis | 2.835232E-03 |
| cell cycle checkpoint | 6 | Chicken | Cerebellum | 2.630717E-07 |
| cell cycle checkpoint | 6 | Chicken | Kidney | 1.924268E-04 |
| cell cycle checkpoint | 6 | Chicken | Ovary | 1.808174E-03 |
| cell cycle checkpoint | 6 | Human | Brain | 2.789292E-03 |
| cell cycle checkpoint | 6 | Human | Cerebellum | 1.584118E-02 |
| cell cycle checkpoint | 6 | Human | Ovary | 4.865058E-03 |
| cell cycle checkpoint | 6 | Mouse | Brain | 1.698017E-03 |
| cell cycle checkpoint | 6 | Mouse | Cerebellum | 5.698712E-05 |
| cell cycle checkpoint | 6 | Mouse | Ovary | 3.819684E-03 |
| cell cycle checkpoint | 6 | Opossum | Brain | 1.776163E-06 |
| cell cycle checkpoint | 6 | Opossum | Liver | 4.926658E-02 |

| GO term | Number of species enriched | Species | Organ | p-value |
|---|---|---|---|---|
| cell cycle checkpoint | 6 | Rabbit | Brain | 3.595941E-06 |
| cell cycle checkpoint | 6 | Rabbit | Cerebellum | 6.539374E-07 |
| cell cycle checkpoint | 6 | Rabbit | Heart | 2.122577E-04 |
| cell cycle checkpoint | 6 | Rabbit | Kidney | 1.695206E-02 |
| cell cycle checkpoint | 6 | Rabbit | Ovary | 6.158613E-04 |
| cell cycle checkpoint | 6 | Rabbit | Testis | 9.270817E-04 |
| cell cycle checkpoint | 6 | Rat | Cerebellum | 2.330141E-03 |
| cell cycle checkpoint | 6 | Rat | Kidney | 5.828283E-03 |
| carboxylic acid metabolic process | 6 | Chicken | Brain | 1.154380E-02 |
| carboxylic acid metabolic process | 6 | Chicken | Kidney | 3.028091E-02 |
| carboxylic acid metabolic process | 6 | Human | Cerebellum | 3.977122E-03 |
| carboxylic acid metabolic process | 6 | Human | Ovary | 1.376367E-03 |
| carboxylic acid metabolic process | 6 | Mouse | Brain | 2.103323E-03 |
| carboxylic acid metabolic process | 6 | Opossum | Ovary | 6.964023E-03 |
| carboxylic acid metabolic process | 6 | Rabbit | Kidney | 1.693314E-05 |
| carboxylic acid metabolic process | 6 | Rabbit | Testis | 2.859000E-02 |

| GO term | Number of species enriched | Species | Organ | p-value |
|---|---|---|---|---|
| carboxylic acid metabolic process | 6 | Rat | Liver | 4.766473E-09 |
| carboxylic acid metabolic process | 6 | Rat | Ovary | 9.305553E-04 |
| axon | 6 | Chicken | Kidney | 2.384023E-04 |
| axon | 6 | Human | Brain | 1.840326E-02 |
| axon | 6 | Human | Kidney | 3.919564E-02 |
| axon | 6 | Human | Liver | 1.324852E-03 |
| axon | 6 | Human | Testis | 2.743163E-02 |
| axon | 6 | Opossum | Ovary | 9.600147E-03 |
| axon | 6 | Rabbit | Liver | 7.834043E-05 |
| axon | 6 | Rat | Brain | 4.379190E-05 |
| axon | 6 | RhesusMacaque | Brain | 6.499734E-04 |
| RNA processing | 6 | Chicken | Cerebellum | 8.237590E-20 |
| RNA processing | 6 | Chicken | Kidney | 1.351697E-04 |
| RNA processing | 6 | Chicken | Testis | 7.577432E-10 |
| RNA processing | 6 | Human | Brain | 1.047086E-03 |
| RNA processing | 6 | Human | Cerebellum | 9.703456E-04 |
| RNA processing | 6 | Human | Liver | 2.127377E-03 |
| RNA processing | 6 | Human | Ovary | 4.259265E-04 |
| RNA processing | 6 | Mouse | Brain | 2.623146E-08 |
| RNA processing | 6 | Mouse | Cerebellum | 2.096276E-03 |
| RNA processing | 6 | Mouse | Heart | 5.361633E-05 |

| GO term | Number of species enriched | Species | Organ | p-value |
|---|---|---|---|---|
| RNA processing | 6 | Mouse | Ovary | 4.216052E-05 |
| RNA processing | 6 | Opossum | Brain | 4.518373E-18 |
| RNA processing | 6 | Opossum | Liver | 1.302220E-05 |
| RNA processing | 6 | Rabbit | Brain | 2.004826E-10 |
| RNA processing | 6 | Rabbit | Cerebellum | 4.058408E-15 |
| RNA processing | 6 | Rabbit | Heart | 3.801276E-12 |
| RNA processing | 6 | Rabbit | Kidney | 1.398885E-04 |
| RNA processing | 6 | Rabbit | Ovary | 1.172281E-11 |
| RNA processing | 6 | Rabbit | Testis | 1.559699E-05 |
| RNA processing | 6 | Rat | Cerebellum | 1.510256E-06 |
| RNA processing | 6 | Rat | Kidney | 4.509501E-07 |
| RNA processing | 6 | Rat | Testis | 1.695834E-08 |
| G1/S transition of mitotic cell cycle | 6 | Chicken | Kidney | 1.543112E-02 |
| G1/S transition of mitotic cell cycle | 6 | Human | Cerebellum | 2.372715E-02 |
| G1/S transition of mitotic cell cycle | 6 | Human | Ovary | 6.915629E-03 |
| G1/S transition of mitotic cell cycle | 6 | Mouse | Brain | 3.374395E-04 |
| G1/S transition of mitotic cell cycle | 6 | Mouse | Heart | 6.537038E-03 |

| GO term | Number of species enriched | Species | Organ | p-value |
|---|---|---|---|---|
| G1/S transition of mitotic cell cycle | 6 | Opossum | Brain | 2.814530E-06 |
| G1/S transition of mitotic cell cycle | 6 | Rabbit | Brain | 5.949679E-05 |
| G1/S transition of mitotic cell cycle | 6 | Rabbit | Cerebellum | 7.695069E-03 |
| G1/S transition of mitotic cell cycle | 6 | Rabbit | Heart | 3.270520E-04 |
| G1/S transition of mitotic cell cycle | 6 | Rabbit | Ovary | 1.286325E-05 |
| G1/S transition of mitotic cell cycle | 6 | Rat | Cerebellum | 3.564360E-03 |
| G1/S transition of mitotic cell cycle | 6 | Rat | Liver | 2.628611E-04 |
| DNA strand elongation involved in DNA replication | 6 | Chicken | Cerebellum | 6.765780E-05 |
| DNA strand elongation involved in DNA replication | 6 | Chicken | Kidney | 1.196310E-02 |
| DNA strand elongation involved in DNA replication | 6 | Human | Cerebellum | 1.545430E-03 |
| DNA strand elongation involved in DNA replication | 6 | Human | Ovary | 1.362707E-03 |
| DNA strand elongation involved in DNA replication | 6 | Mouse | Brain | 1.662692E-02 |

| GO term | Number of species enriched | Species | Organ | p-value |
|---|---|---|---|---|
| DNA strand elongation involved in DNA replication | 6 | Mouse | Cerebellum | 2.412737E-02 |
| DNA strand elongation involved in DNA replication | 6 | Opossum | Brain | 1.156146E-04 |
| DNA strand elongation involved in DNA replication | 6 | Rabbit | Brain | 7.101125E-04 |
| DNA strand elongation involved in DNA replication | 6 | Rabbit | Cerebellum | 5.712741E-04 |
| DNA strand elongation involved in DNA replication | 6 | Rabbit | Heart | 2.136469E-03 |
| DNA strand elongation involved in DNA replication | 6 | Rabbit | Ovary | 2.842948E-03 |
| DNA strand elongation involved in DNA replication | 6 | Rabbit | Testis | 1.153272E-02 |
| DNA strand elongation involved in DNA replication | 6 | Rat | Cerebellum | 1.938490E-02 |
| DNA replication | 6 | Chicken | Cerebellum | 2.807875E-13 |
| DNA replication | 6 | Chicken | Kidney | 4.440222E-05 |
| DNA replication | 6 | Chicken | Ovary | 1.276614E-03 |
| DNA replication | 6 | Human | Brain | 2.928687E-03 |
| DNA replication | 6 | Human | Cerebellum | 2.642954E-06 |
| DNA replication | 6 | Human | Ovary | 1.554050E-05 |
| DNA replication | 6 | Mouse | Brain | 1.304974E-05 |
| DNA replication | 6 | Mouse | Cerebellum | 1.070796E-07 |

| GO term | Number of species enriched | Species | Organ | p-value |
|---|---|---|---|---|
| DNA replication | 6 | Mouse | Heart | 3.456692E-04 |
| DNA replication | 6 | Mouse | Ovary | 6.061123E-03 |
| DNA replication | 6 | Mouse | Testis | 1.471783E-02 |
| DNA replication | 6 | Opossum | Brain | 1.326901E-07 |
| DNA replication | 6 | Opossum | Liver | 2.119607E-02 |
| DNA replication | 6 | Rabbit | Brain | 1.205737E-08 |
| DNA replication | 6 | Rabbit | Cerebellum | 6.851967E-09 |
| DNA replication | 6 | Rabbit | Heart | 5.317607E-09 |
| DNA replication | 6 | Rabbit | Kidney | 3.989408E-03 |
| DNA replication | 6 | Rabbit | Ovary | 2.972021E-09 |
| DNA replication | 6 | Rabbit | Testis | 7.383536E-04 |
| DNA replication | 6 | Rat | Cerebellum | 4.549834E-04 |
| DNA repair | 6 | Chicken | Cerebellum | 3.609816E-16 |
| DNA repair | 6 | Chicken | Kidney | 4.592631E-06 |
| DNA repair | 6 | Human | Brain | 5.763925E-07 |
| DNA repair | 6 | Human | Cerebellum | 5.085068E-05 |
| DNA repair | 6 | Human | Ovary | 1.494034E-12 |
| DNA repair | 6 | Human | Testis | 1.144202E-02 |
| DNA repair | 6 | Mouse | Brain | 2.097171E-07 |
| DNA repair | 6 | Mouse | Cerebellum | 2.087381E-09 |
| DNA repair | 6 | Mouse | Ovary | 2.941448E-03 |
| DNA repair | 6 | Mouse | Testis | 3.513022E-03 |
| DNA repair | 6 | Opossum | Brain | 9.542162E-12 |
| DNA repair | 6 | Opossum | Liver | 2.685317E-05 |
| DNA repair | 6 | Rabbit | Brain | 3.886736E-07 |
| DNA repair | 6 | Rabbit | Cerebellum | 3.231315E-11 |
| DNA repair | 6 | Rabbit | Heart | 1.839556E-06 |
| DNA repair | 6 | Rabbit | Kidney | 1.531564E-02 |
| DNA repair | 6 | Rabbit | Liver | 1.269786E-05 |

| GO term | Number of species enriched | Species | Organ | p-value |
|---|---|---|---|---|
| DNA repair | 6 | Rabbit | Ovary | 5.621238E-11 |
| DNA repair | 6 | Rabbit | Testis | 6.776870E-05 |
| DNA repair | 6 | Rat | Cerebellum | 1.844364E-08 |
| DNA repair | 6 | Rat | Kidney | 2.004487E-03 |
| DNA repair | 6 | Rat | Testis | 6.700638E-05 |
| DNA metabolic process | 6 | Chicken | Cerebellum | 2.462976E-17 |
| DNA metabolic process | 6 | Chicken | Kidney | 9.880197E-07 |
| DNA metabolic process | 6 | Human | Brain | 9.897544E-05 |
| DNA metabolic process | 6 | Human | Cerebellum | 2.047209E-04 |
| DNA metabolic process | 6 | Human | Ovary | 1.199943E-10 |
| DNA metabolic process | 6 | Mouse | Brain | 5.063531E-04 |
| DNA metabolic process | 6 | Mouse | Cerebellum | 1.755285E-08 |
| DNA metabolic process | 6 | Mouse | Ovary | 2.819894E-02 |
| DNA metabolic process | 6 | Opossum | Brain | 7.639706E-13 |
| DNA metabolic process | 6 | Opossum | Liver | 1.090607E-02 |
| DNA metabolic process | 6 | Rabbit | Brain | 1.285625E-07 |
| DNA metabolic process | 6 | Rabbit | Cerebellum | 8.510760E-12 |
| DNA metabolic process | 6 | Rabbit | Heart | 1.997789E-07 |
| DNA metabolic process | 6 | Rabbit | Liver | 1.314207E-02 |
| DNA metabolic process | 6 | Rabbit | Ovary | 1.098283E-10 |
| DNA metabolic process | 6 | Rabbit | Testis | 2.143685E-04 |

| GO term | Number of species enriched | Species | Organ | p-value |
|---|---|---|---|---|
| DNA metabolic process | 6 | Rat | Cerebellum | 2.122615E-07 |
| DNA metabolic process | 6 | Rat | Kidney | 1.203981E-02 |
| DNA metabolic process | 6 | Rat | Testis | 1.680453E-03 |
| ATPase activity, coupled | 6 | Chicken | Cerebellum | 5.361193E-03 |
| ATPase activity, coupled | 6 | Chicken | Kidney | 2.113306E-02 |
| ATPase activity, coupled | 6 | Human | Brain | 4.697823E-04 |
| ATPase activity, coupled | 6 | Human | Cerebellum | 2.571029E-06 |
| ATPase activity, coupled | 6 | Human | Ovary | 4.467406E-02 |
| ATPase activity, coupled | 6 | Mouse | Brain | 1.429374E-02 |
| ATPase activity, coupled | 6 | Mouse | Cerebellum | 5.095316E-03 |
| ATPase activity, coupled | 6 | Mouse | Ovary | 3.877400E-02 |
| ATPase activity, coupled | 6 | Opossum | Brain | 7.690678E-03 |
| ATPase activity, coupled | 6 | Rabbit | Brain | 1.543735E-02 |
| ATPase activity, coupled | 6 | Rabbit | Cerebellum | 2.111841E-04 |
| ATPase activity, coupled | 6 | Rabbit | Kidney | 1.165822E-02 |

| GO term | Number of species enriched | Species | Organ | p-value |
|---|---|---|---|---|
| ATPase activity, coupled | 6 | Rabbit | Ovary | 7.278081E-03 |
| ATPase activity, coupled | 6 | Rabbit | Testis | 1.993731E-02 |
| ATPase activity, coupled | 6 | Rat | Cerebellum | 1.098337E-02 |
| ATPase activity, coupled | 6 | Rat | Kidney | 5.154937E-05 |
| ATPase activity, coupled | 6 | Rat | Liver | 3.493199E-02 |
| ATPase activity | 6 | Chicken | Cerebellum | 8.990724E-05 |
| ATPase activity | 6 | Chicken | Kidney | 1.084118E-03 |
| ATPase activity | 6 | Human | Brain | 4.716640E-06 |
| ATPase activity | 6 | Human | Cerebellum | 3.333226E-06 |
| ATPase activity | 6 | Human | Ovary | 3.624602E-04 |
| ATPase activity | 6 | Mouse | Brain | 8.805795E-04 |
| ATPase activity | 6 | Mouse | Cerebellum | 4.442524E-04 |
| ATPase activity | 6 | Mouse | Heart | 3.669068E-02 |
| ATPase activity | 6 | Mouse | Ovary | 3.611096E-03 |
| ATPase activity | 6 | Mouse | Testis | 1.643693E-02 |
| ATPase activity | 6 | Opossum | Brain | 1.889248E-05 |
| ATPase activity | 6 | Opossum | Liver | 7.042771E-03 |
| ATPase activity | 6 | Rabbit | Brain | 1.319404E-03 |
| ATPase activity | 6 | Rabbit | Cerebellum | 1.897392E-05 |
| ATPase activity | 6 | Rabbit | Heart | 1.275263E-02 |
| ATPase activity | 6 | Rabbit | Kidney | 3.424753E-03 |
| ATPase activity | 6 | Rabbit | Liver | 9.499723E-03 |
| ATPase activity | 6 | Rabbit | Ovary | 8.968628E-04 |
| ATPase activity | 6 | Rabbit | Testis | 1.677099E-04 |

| GO term | Number of species enriched | Species | Organ | p-value |
|---|---|---|---|---|
| ATPase activity | 6 | Rat | Cerebellum | 1.489189E-03 |
| ATPase activity | 6 | Rat | Kidney | 3.907998E-06 |
| ubiquitin-dependent protein catabolic process | 5 | Chicken | Kidney | 2.593843E-02 |
| ubiquitin-dependent protein catabolic process | 5 | Human | Brain | 7.255602E-03 |
| ubiquitin-dependent protein catabolic process | 5 | Mouse | Ovary | 1.006519E-03 |
| ubiquitin-dependent protein catabolic process | 5 | Opossum | Liver | 9.404363E-04 |
| ubiquitin-dependent protein catabolic process | 5 | Rabbit | Cerebellum | 1.916714E-03 |
| ribose phosphate metabolic process | 5 | Human | Cerebellum | 3.489565E-03 |
| ribose phosphate metabolic process | 5 | Human | Ovary | 1.787638E-02 |
| ribose phosphate metabolic process | 5 | Mouse | Brain | 2.220617E-02 |
| ribose phosphate metabolic process | 5 | Opossum | Liver | 8.039426E-03 |

| GO term | Number of species enriched | Species | Organ | p-value |
|---|---|---|---|---|
| ribose phosphate metabolic process | 5 | Rabbit | Brain | 3.098049E-02 |
| ribose phosphate metabolic process | 5 | Rabbit | Kidney | 7.460618E-03 |
| ribose phosphate metabolic process | 5 | Rat | Liver | 3.476704E-04 |
| regulation of protein serine/threonine kinase activity | 5 | Chicken | Brain | 8.469255E-04 |
| regulation of protein serine/threonine kinase activity | 5 | Mouse | Ovary | 1.844617E-02 |
| regulation of protein serine/threonine kinase activity | 5 | Opossum | Liver | 3.168675E-02 |
| regulation of protein serine/threonine kinase activity | 5 | Opossum | Ovary | 6.252525E-03 |
| regulation of protein serine/threonine kinase activity | 5 | Rat | Liver | 4.475558E-03 |
| regulation of protein serine/threonine kinase activity | 5 | RhesusMacaque | Brain | 3.536226E-03 |
| pyrophosphatase activity | 5 | Chicken | Cerebellum | 2.472611E-02 |
| pyrophosphatase activity | 5 | Chicken | Kidney | 1.798102E-02 |
| pyrophosphatase activity | 5 | Human | Brain | 6.388778E-04 |

| GO term | Number of species enriched | Species | Organ | p-value |
|---|---|---|---|---|
| pyrophosphatase activity | 5 | Human | Cerebellum | 8.881242E-04 |
| pyrophosphatase activity | 5 | Human | Ovary | 5.342761E-03 |
| pyrophosphatase activity | 5 | Opossum | Brain | 2.494670E-02 |
| pyrophosphatase activity | 5 | Rabbit | Brain | 3.289862E-02 |
| pyrophosphatase activity | 5 | Rabbit | Testis | 2.757548E-03 |
| pyrophosphatase activity | 5 | Rat | Kidney | 3.354246E-03 |
| purine nucleotide metabolic process | 5 | Human | Cerebellum | 1.302256E-02 |
| purine nucleotide metabolic process | 5 | Human | Ovary | 6.378386E-03 |
| purine nucleotide metabolic process | 5 | Human | Testis | 3.891843E-02 |
| purine nucleotide metabolic process | 5 | Mouse | Brain | 2.477256E-02 |
| purine nucleotide metabolic process | 5 | Opossum | Liver | 1.095682E-02 |
| purine nucleotide metabolic process | 5 | Rabbit | Brain | 1.964719E-02 |
| purine nucleotide metabolic process | 5 | Rabbit | Heart | 3.862447E-02 |

| GO term | Number of species enriched | Species | Organ | p-value |
|---|---|---|---|---|
| purine nucleotide metabolic process | 5 | Rabbit | Kidney | 1.443201E-02 |
| purine nucleotide metabolic process | 5 | Rabbit | Testis | 3.816806E-02 |
| purine nucleotide metabolic process | 5 | Rat | Liver | 7.126757E-04 |
| proteolysis involved in cellular protein catabolic process | 5 | Chicken | Kidney | 3.693176E-03 |
| proteolysis involved in cellular protein catabolic process | 5 | Human | Brain | 3.697177E-04 |
| proteolysis involved in cellular protein catabolic process | 5 | Mouse | Ovary | 5.383252E-04 |
| proteolysis involved in cellular protein catabolic process | 5 | Opossum | Heart | 2.534228E-02 |
| proteolysis involved in cellular protein catabolic process | 5 | Opossum | Liver | 1.734368E-04 |
| proteolysis involved in cellular protein catabolic process | 5 | Rabbit | Brain | 4.267453E-02 |

| GO term | Number of species enriched | Species | Organ | p-value |
|---|---|---|---|---|
| proteolysis involved in cellular protein catabolic process | 5 | Rabbit | Cerebellum | 1.926496E-03 |
| proteolysis involved in cellular protein catabolic process | 5 | Rabbit | Heart | 1.637122E-02 |
| proteolysis involved in cellular protein catabolic process | 5 | Rabbit | Liver | 2.874799E-02 |
| protein autophosphorylation | 5 | Human | Cerebellum | 4.692973E-02 |
| protein autophosphorylation | 5 | Mouse | Testis | 2.926253E-02 |
| protein autophosphorylation | 5 | Opossum | Kidney | 1.358045E-02 |
| protein autophosphorylation | 5 | Opossum | Testis | 3.438921E-02 |
| protein autophosphorylation | 5 | Rat | Brain | 4.346992E-02 |
| protein autophosphorylation | 5 | RhesusMacaque | Brain | 2.465457E-04 |
| positive regulation of transferase activity | 5 | Chicken | Brain | 8.653703E-07 |
| positive regulation of transferase activity | 5 | Human | Brain | 2.933808E-02 |
| positive regulation of transferase activity | 5 | Human | Kidney | 1.859370E-02 |
| positive regulation of transferase activity | 5 | Mouse | Ovary | 1.099107E-02 |

| GO term | Number of species enriched | Species | Organ | p-value |
|---|---|---|---|---|
| positive regulation of transferase activity | 5 | Mouse | Testis | 2.105577E-02 |
| positive regulation of transferase activity | 5 | Opossum | Heart | 4.395255E-02 |
| positive regulation of transferase activity | 5 | Opossum | Kidney | 2.574575E-03 |
| positive regulation of transferase activity | 5 | Rat | Liver | 1.014600E-02 |
| positive regulation of cell projection organization | 5 | Chicken | Brain | 3.634561E-02 |
| positive regulation of cell projection organization | 5 | Human | Kidney | 1.324541E-02 |
| positive regulation of cell projection organization | 5 | Human | Liver | 8.046080E-03 |
| positive regulation of cell projection organization | 5 | Opossum | Ovary | 4.520881E-02 |
| positive regulation of cell projection organization | 5 | Rat | Brain | 4.833460E-04 |
| positive regulation of cell projection organization | 5 | RhesusMacaque | Brain | 5.196210E-03 |
| positive regulation of cell projection organization | 5 | RhesusMacaque | Liver | 3.000566E-02 |

| GO term | Number of species enriched | Species | Organ | p-value |
|---|---|---|---|---|
| phospholipid binding | 5 | Chicken | Brain | 3.256829E-06 |
| phospholipid binding | 5 | Human | Brain | 4.148380E-03 |
| phospholipid binding | 5 | Human | Liver | 2.588545E-02 |
| phospholipid binding | 5 | Opossum | Kidney | 1.462064E-02 |
| phospholipid binding | 5 | Opossum | Liver | 1.682749E-03 |
| phospholipid binding | 5 | Opossum | Ovary | 1.483507E-04 |
| phospholipid binding | 5 | Opossum | Testis | 2.744052E-03 |
| phospholipid binding | 5 | Rat | Brain | 5.449691E-03 |
| phospholipid binding | 5 | Rat | Liver | 2.207137E-02 |
| phospholipid binding | 5 | RhesusMacaque | Brain | 1.407439E-02 |
| phospholipid binding | 5 | RhesusMacaque | Liver | 3.124562E-02 |
| phospholipid binding | 5 | RhesusMacaque | Testis | 4.336237E-03 |
| nucleoside-triphosphatase activity | 5 | Chicken | Cerebellum | 7.415093E-03 |
| nucleoside-triphosphatase activity | 5 | Chicken | Kidney | 2.301387E-02 |
| nucleoside-triphosphatase activity | 5 | Human | Brain | 7.221664E-04 |
| nucleoside-triphosphatase activity | 5 | Human | Cerebellum | 6.915592E-04 |
| nucleoside-triphosphatase activity | 5 | Human | Ovary | 5.211246E-03 |

| GO term | Number of species enriched | Species | Organ | p-value |
|---|---|---|---|---|
| nucleoside-triphosphatase activity | 5 | Opossum | Brain | 2.742981E-02 |
| nucleoside-triphosphatase activity | 5 | Rabbit | Brain | 3.015250E-02 |
| nucleoside-triphosphatase activity | 5 | Rabbit | Testis | 1.416072E-03 |
| nucleoside-triphosphatase activity | 5 | Rat | Kidney | 1.572176E-03 |
| nuclear transport | 5 | Chicken | Cerebellum | 1.004690E-04 |
| nuclear transport | 5 | Human | Liver | 4.486880E-03 |
| nuclear transport | 5 | Opossum | Brain | 1.074786E-05 |
| nuclear transport | 5 | Rabbit | Cerebellum | 3.966319E-02 |
| nuclear transport | 5 | Rat | Testis | 1.898062E-02 |
| nuclear envelope | 5 | Chicken | Kidney | 2.932812E-03 |
| nuclear envelope | 5 | Human | Cerebellum | 4.825378E-03 |
| nuclear envelope | 5 | Human | Testis | 1.613632E-03 |
| nuclear envelope | 5 | Mouse | Brain | 4.030416E-03 |
| nuclear envelope | 5 | Rabbit | Brain | 1.035983E-04 |
| nuclear envelope | 5 | Rabbit | Cerebellum | 6.793728E-05 |
| nuclear envelope | 5 | Rabbit | Heart | 1.563262E-02 |
| nuclear envelope | 5 | Rabbit | Ovary | 1.455838E-03 |
| nuclear envelope | 5 | Rat | Cerebellum | 3.777669E-02 |

| GO term | Number of species enriched | Species | Organ | p-value |
|---|---|---|---|---|
| nuclear body | 5 | Chicken | Cerebellum | 1.069181E-02 |
| nuclear body | 5 | Chicken | Testis | 4.527609E-02 |
| nuclear body | 5 | Mouse | Cerebellum | 2.194539E-02 |
| nuclear body | 5 | Opossum | Brain | 1.469471E-02 |
| nuclear body | 5 | Rabbit | Cerebellum | 1.479831E-06 |
| nuclear body | 5 | Rat | Cerebellum | 7.021655E-04 |
| modification-dependent protein catabolic process | 5 | Chicken | Kidney | 3.651084E-02 |
| modification-dependent protein catabolic process | 5 | Human | Brain | 7.109978E-03 |
| modification-dependent protein catabolic process | 5 | Mouse | Ovary | 7.777122E-04 |
| modification-dependent protein catabolic process | 5 | Opossum | Liver | 9.451609E-04 |
| modification-dependent protein catabolic process | 5 | Rabbit | Cerebellum | 4.889950E-03 |
| mitotic cell cycle checkpoint | 5 | Chicken | Cerebellum | 4.876221E-05 |
| mitotic cell cycle checkpoint | 5 | Chicken | Kidney | 3.628431E-02 |
| mitotic cell cycle checkpoint | 5 | Mouse | Brain | 4.046209E-02 |

| GO term | Number of species enriched | Species | Organ | p-value |
|---|---|---|---|---|
| mitotic cell cycle checkpoint | 5 | Mouse | Cerebellum | 7.614826E-03 |
| mitotic cell cycle checkpoint | 5 | Opossum | Brain | 1.496933E-02 |
| mitotic cell cycle checkpoint | 5 | Rabbit | Brain | 5.826835E-03 |
| mitotic cell cycle checkpoint | 5 | Rabbit | Cerebellum | 9.375875E-05 |
| mitotic cell cycle checkpoint | 5 | Rabbit | Heart | 2.000644E-02 |
| mitotic cell cycle checkpoint | 5 | Rabbit | Ovary | 4.078413E-02 |
| mitotic cell cycle checkpoint | 5 | Rat | Kidney | 3.307269E-02 |
| membrane organization | 5 | Chicken | Heart | 3.248981E-02 |
| membrane organization | 5 | Human | Brain | 2.426455E-03 |
| membrane organization | 5 | Human | Kidney | 1.886700E-02 |
| membrane organization | 5 | Human | Liver | 4.867503E-04 |
| membrane organization | 5 | Mouse | Liver | 4.262706E-02 |
| membrane organization | 5 | Mouse | Testis | 1.568408E-03 |
| membrane organization | 5 | Opossum | Kidney | 4.812320E-04 |
| membrane organization | 5 | Opossum | Liver | 4.247713E-05 |
| membrane organization | 5 | Opossum | Ovary | 2.921108E-02 |

| GO term | Number of species enriched | Species | Organ | p-value |
|---|---|---|---|---|
| membrane organization | 5 | Rabbit | Testis | 1.909833E-02 |
| lamellipodium | 5 | Chicken | Testis | 1.482102E-02 |
| lamellipodium | 5 | Human | Brain | 4.188757E-02 |
| lamellipodium | 5 | Human | Kidney | 1.387264E-03 |
| lamellipodium | 5 | Human | Liver | 2.424014E-02 |
| lamellipodium | 5 | Opossum | Kidney | 5.251291E-05 |
| lamellipodium | 5 | Opossum | Ovary | 3.964217E-04 |
| lamellipodium | 5 | Rat | Brain | 1.320619E-02 |
| lamellipodium | 5 | RhesusMacaque | Liver | 4.081767E-02 |
| intracellular protein transport | 5 | Chicken | Testis | 8.142040E-04 |
| intracellular protein transport | 5 | Human | Brain | 6.073308E-04 |
| intracellular protein transport | 5 | Opossum | Liver | 7.177241E-06 |
| intracellular protein transport | 5 | Rabbit | Liver | 9.442036E-05 |
| intracellular protein transport | 5 | Rat | Liver | 1.403714E-03 |
| hydrolase activity, acting on acid anhydrides, in phosphorus-containing anhydrides | 5 | Chicken | Cerebellum | 2.829547E-02 |
| hydrolase activity, acting on acid anhydrides, in phosphorus-containing anhydrides | 5 | Chicken | Kidney | 2.116402E-02 |

| GO term | Number of species enriched | Species | Organ | p-value |
| --- | --- | --- | --- | --- |
| hydrolase activity, acting on acid anhydrides, in phosphorus-containing anhydrides | 5 | Human | Brain | 7.911187E-04 |
| hydrolase activity, acting on acid anhydrides, in phosphorus-containing anhydrides | 5 | Human | Cerebellum | 1.064941E-03 |
| hydrolase activity, acting on acid anhydrides, in phosphorus-containing anhydrides | 5 | Human | Ovary | 6.138926E-03 |
| hydrolase activity, acting on acid anhydrides, in phosphorus-containing anhydrides | 5 | Opossum | Brain | 2.835786E-02 |
| hydrolase activity, acting on acid anhydrides, in phosphorus-containing anhydrides | 5 | Rabbit | Brain | 3.798663E-02 |
| hydrolase activity, acting on acid anhydrides, in phosphorus-containing anhydrides | 5 | Rabbit | Testis | 3.323407E-03 |

| GO term | Number of species enriched | Species | Organ | p-value |
|---|---|---|---|---|
| hydrolase activity, acting on acid anhydrides, in phosphorus-containing anhydrides | 5 | Rat | Kidney | 3.937914E-03 |
| hydrolase activity, acting on acid anhydrides | 5 | Chicken | Cerebellum | 2.829547E-02 |
| hydrolase activity, acting on acid anhydrides | 5 | Chicken | Kidney | 2.116402E-02 |
| hydrolase activity, acting on acid anhydrides | 5 | Human | Brain | 7.911187E-04 |
| hydrolase activity, acting on acid anhydrides | 5 | Human | Cerebellum | 1.064941E-03 |
| hydrolase activity, acting on acid anhydrides | 5 | Human | Ovary | 6.138926E-03 |
| hydrolase activity, acting on acid anhydrides | 5 | Opossum | Brain | 2.835786E-02 |
| hydrolase activity, acting on acid anhydrides | 5 | Rabbit | Brain | 3.798663E-02 |
| hydrolase activity, acting on acid anhydrides | 5 | Rabbit | Testis | 3.323407E-03 |
| hydrolase activity, acting on acid anhydrides | 5 | Rat | Kidney | 3.937914E-03 |
| helicase activity | 5 | Chicken | Cerebellum | 5.751763E-07 |

| GO term | Number of species enriched | Species | Organ | p-value |
|---|---|---|---|---|
| helicase activity | 5 | Human | Ovary | 9.808702E-03 |
| helicase activity | 5 | Mouse | Heart | 1.008766E-02 |
| helicase activity | 5 | Opossum | Brain | 5.062843E-04 |
| helicase activity | 5 | Rabbit | Cerebellum | 2.543367E-02 |
| helicase activity | 5 | Rabbit | Ovary | 2.119392E-02 |
| double-strand break repair | 5 | Chicken | Cerebellum | 3.345351E-05 |
| double-strand break repair | 5 | Human | Ovary | 3.430807E-06 |
| double-strand break repair | 5 | Mouse | Brain | 2.016792E-02 |
| double-strand break repair | 5 | Opossum | Brain | 1.960411E-02 |
| double-strand break repair | 5 | Rabbit | Cerebellum | 3.325543E-04 |
| double-strand break repair | 5 | Rabbit | Ovary | 5.249272E-04 |
| cytoskeleton-dependent intracellular transport | 5 | Chicken | Ovary | 4.091251E-02 |
| cytoskeleton-dependent intracellular transport | 5 | Human | Brain | 3.912029E-03 |
| cytoskeleton-dependent intracellular transport | 5 | Human | Liver | 5.924472E-05 |
| cytoskeleton-dependent intracellular transport | 5 | Opossum | Kidney | 1.757586E-02 |

| GO term | Number of species enriched | Species | Organ | p-value |
|---|---|---|---|---|
| cytoskeleton-dependent intracellular transport | 5 | Opossum | Liver | 6.707591E-05 |
| cytoskeleton-dependent intracellular transport | 5 | Rabbit | Liver | 1.043947E-02 |
| cytoskeleton-dependent intracellular transport | 5 | RhesusMacaque | Heart | 3.175391E-02 |
| cytoskeleton-dependent intracellular transport | 5 | RhesusMacaque | Kidney | 1.710950E-02 |
| cytoskeletal protein binding | 5 | Chicken | Ovary | 3.946820E-02 |
| cytoskeletal protein binding | 5 | Human | Brain | 1.072754E-04 |
| cytoskeletal protein binding | 5 | Human | Cerebellum | 6.882710E-03 |
| cytoskeletal protein binding | 5 | Human | Liver | 6.026736E-05 |
| cytoskeletal protein binding | 5 | Human | Testis | 3.248181E-04 |
| cytoskeletal protein binding | 5 | Opossum | Kidney | 1.060651E-08 |
| cytoskeletal protein binding | 5 | Opossum | Liver | 1.652092E-02 |
| cytoskeletal protein binding | 5 | Rat | Brain | 1.005410E-03 |
| cytoskeletal protein binding | 5 | Rat | Kidney | 1.361962E-02 |
| cytoskeletal protein binding | 5 | RhesusMacaque | Brain | 2.439879E-05 |
| cytoskeletal protein binding | 5 | RhesusMacaque | Kidney | 2.121810E-02 |
| cytoskeletal protein binding | 5 | RhesusMacaque | Testis | 3.645870E-04 |

| GO term | Number of species enriched | Species | Organ | p-value |
|---|---|---|---|---|
| centrosome localization | 5 | Chicken | Cerebellum | 7.784892E-03 |
| centrosome localization | 5 | Human | Liver | 1.887230E-03 |
| centrosome localization | 5 | Opossum | Liver | 7.236515E-04 |
| centrosome localization | 5 | Opossum | Testis | 5.632089E-03 |
| centrosome localization | 5 | Rat | Kidney | 4.799615E-02 |
| centrosome localization | 5 | RhesusMacaque | Kidney | 4.780151E-02 |
| centrosome localization | 5 | RhesusMacaque | Liver | 1.742106E-03 |
| G2/M transition of mitotic cell cycle | 5 | Chicken | Cerebellum | 3.161283E-02 |
| G2/M transition of mitotic cell cycle | 5 | Human | Brain | 1.146205E-02 |
| G2/M transition of mitotic cell cycle | 5 | Mouse | Cerebellum | 1.644859E-02 |
| G2/M transition of mitotic cell cycle | 5 | Rabbit | Cerebellum | 3.415631E-02 |
| G2/M transition of mitotic cell cycle | 5 | Rat | Cerebellum | 1.100649E-02 |
| transport along microtubule | 4 | Chicken | Ovary | 3.440789E-02 |
| transport along microtubule | 4 | Human | Brain | 4.524100E-03 |
| transport along microtubule | 4 | Human | Liver | 5.671647E-04 |
| transport along microtubule | 4 | Opossum | Liver | 1.796672E-04 |
| transport along microtubule | 4 | Rabbit | Liver | 4.083569E-02 |

| GO term | Number of species enriched | Species | Organ | p-value |
|---|---|---|---|---|
| tRNA processing | 4 | Chicken | Cerebellum | 1.427667E-02 |
| tRNA processing | 4 | Mouse | Brain | 7.057603E-05 |
| tRNA processing | 4 | Mouse | Heart | 9.499196E-03 |
| tRNA processing | 4 | Opossum | Brain | 3.933927E-03 |
| tRNA processing | 4 | Rabbit | Brain | 1.190145E-03 |
| tRNA processing | 4 | Rabbit | Heart | 1.888497E-02 |
| synaptic membrane | 4 | Human | Cerebellum | 4.051754E-02 |
| synaptic membrane | 4 | Rabbit | Brain | 2.620512E-02 |
| synaptic membrane | 4 | Rat | Brain | 2.037402E-02 |
| synaptic membrane | 4 | RhesusMacaque | Brain | 9.875540E-03 |
| smoothened signaling pathway | 4 | Chicken | Ovary | 7.064442E-03 |
| smoothened signaling pathway | 4 | Human | Brain | 4.980042E-02 |
| smoothened signaling pathway | 4 | Human | Liver | 5.537655E-04 |
| smoothened signaling pathway | 4 | Rabbit | Liver | 4.911080E-02 |
| smoothened signaling pathway | 4 | Rat | Cerebellum | 1.169067E-02 |
| smoothened signaling pathway | 4 | Rat | Liver | 2.790488E-02 |

| GO term | Number of species enriched | Species | Organ | p-value |
|---|---|---|---|---|
| small GTPase mediated signal transduction | 4 | Human | Brain | 7.249438E-03 |
| small GTPase mediated signal transduction | 4 | Mouse | Testis | 1.246185E-02 |
| small GTPase mediated signal transduction | 4 | Opossum | Kidney | 6.770438E-05 |
| small GTPase mediated signal transduction | 4 | Rabbit | Brain | 1.149776E-03 |
| ribonucleotide metabolic process | 4 | Human | Cerebellum | 4.043688E-03 |
| ribonucleotide metabolic process | 4 | Human | Ovary | 1.791629E-02 |
| ribonucleotide metabolic process | 4 | Opossum | Liver | 6.011777E-03 |
| ribonucleotide metabolic process | 4 | Rabbit | Brain | 4.657774E-02 |
| ribonucleotide metabolic process | 4 | Rabbit | Kidney | 4.500837E-03 |
| ribonucleotide metabolic process | 4 | Rat | Liver | 8.245583E-04 |
| regulation of mitotic cell cycle phase transition | 4 | Mouse | Brain | 1.119808E-03 |
| regulation of mitotic cell cycle phase transition | 4 | Mouse | Cerebellum | 2.932671E-02 |

| GO term | Number of species enriched | Species | Organ | p-value |
|---|---|---|---|---|
| regulation of mitotic cell cycle phase transition | 4 | Opossum | Brain | 2.702079E-05 |
| regulation of mitotic cell cycle phase transition | 4 | Rabbit | Cerebellum | 7.136257E-06 |
| regulation of mitotic cell cycle phase transition | 4 | Rabbit | Ovary | 2.543459E-03 |
| regulation of mitotic cell cycle phase transition | 4 | Rat | Cerebellum | 1.728509E-02 |
| regulation of developmental growth | 4 | Human | Kidney | 1.030453E-03 |
| regulation of developmental growth | 4 | Human | Liver | 1.182597E-02 |
| regulation of developmental growth | 4 | Opossum | Testis | 3.882564E-02 |
| regulation of developmental growth | 4 | Rat | Brain | 3.756781E-03 |
| regulation of developmental growth | 4 | RhesusMacaque | Brain | 5.656815E-05 |
| regulation of developmental growth | 4 | RhesusMacaque | Kidney | 1.328439E-02 |
| regulation of cell development | 4 | Chicken | Brain | 3.161184E-02 |
| regulation of cell development | 4 | Human | Kidney | 7.844985E-03 |
| regulation of cell development | 4 | Human | Liver | 9.059395E-03 |

| GO term | Number of species enriched | Species | Organ | p-value |
|---|---|---|---|---|
| regulation of cell development | 4 | Rat | Brain | 2.810669E-03 |
| regulation of cell development | 4 | RhesusMacaque | Brain | 3.639219E-03 |
| regulation of cell cycle phase transition | 4 | Mouse | Brain | 1.711710E-03 |
| regulation of cell cycle phase transition | 4 | Mouse | Cerebellum | 2.040220E-02 |
| regulation of cell cycle phase transition | 4 | Opossum | Brain | 2.266060E-05 |
| regulation of cell cycle phase transition | 4 | Rabbit | Brain | 2.999985E-02 |
| regulation of cell cycle phase transition | 4 | Rabbit | Cerebellum | 2.591228E-05 |
| regulation of cell cycle phase transition | 4 | Rabbit | Heart | 4.927593E-02 |
| regulation of cell cycle phase transition | 4 | Rabbit | Ovary | 8.026320E-03 |
| regulation of cell cycle phase transition | 4 | Rat | Cerebellum | 4.922611E-02 |
| positive regulation of protein phosphorylation | 4 | Chicken | Brain | 3.829248E-05 |

| GO term | Number of species enriched | Species | Organ | p-value |
|---|---|---|---|---|
| positive regulation of protein phosphorylation | 4 | Mouse | Kidney | 1.532608E-03 |
| positive regulation of protein phosphorylation | 4 | Opossum | Kidney | 3.891159E-02 |
| positive regulation of protein phosphorylation | 4 | RhesusMacaque | Brain | 3.474641E-02 |
| positive regulation of cell differentiation | 4 | Chicken | Brain | 8.361970E-03 |
| positive regulation of cell differentiation | 4 | Human | Kidney | 5.993917E-03 |
| positive regulation of cell differentiation | 4 | Rat | Brain | 1.970705E-03 |
| positive regulation of cell differentiation | 4 | RhesusMacaque | Brain | 3.712622E-04 |
| peptidyl-serine phosphorylation | 4 | Human | Kidney | 4.994147E-02 |
| peptidyl-serine phosphorylation | 4 | Mouse | Testis | 2.708405E-02 |
| peptidyl-serine phosphorylation | 4 | Opossum | Kidney | 2.078088E-02 |
| peptidyl-serine phosphorylation | 4 | RhesusMacaque | Brain | 1.347861E-03 |
| peptidyl-serine phosphorylation | 4 | RhesusMacaque | Testis | 4.134668E-02 |
| organic acid transport | 4 | Mouse | Cerebellum | 9.577387E-04 |
| organic acid transport | 4 | Rabbit | Testis | 3.379068E-04 |

| GO term | Number of species enriched | Species | Organ | p-value |
|---|---|---|---|---|
| organic acid transport | 4 | Rat | Brain | 2.814940E-04 |
| organic acid transport | 4 | RhesusMacaque | Brain | 6.311997E-03 |
| organic acid transport | 4 | RhesusMacaque | Kidney | 4.231431E-04 |
| nucleocytoplasmic transport | 4 | Chicken | Cerebellum | 2.336265E-04 |
| nucleocytoplasmic transport | 4 | Human | Liver | 3.804218E-03 |
| nucleocytoplasmic transport | 4 | Opossum | Brain | 2.697309E-05 |
| nucleocytoplasmic transport | 4 | Rat | Testis | 3.695297E-02 |
| nuclear envelope disassembly | 4 | Chicken | Cerebellum | 1.766219E-02 |
| nuclear envelope disassembly | 4 | Mouse | Brain | 4.610675E-03 |
| nuclear envelope disassembly | 4 | Opossum | Brain | 2.394187E-02 |
| nuclear envelope disassembly | 4 | Rabbit | Brain | 5.810251E-03 |
| nuclear envelope disassembly | 4 | Rabbit | Cerebellum | 2.402310E-02 |
| neuronal cell body | 4 | Human | Liver | 1.437234E-03 |
| neuronal cell body | 4 | Mouse | Kidney | 3.590691E-03 |
| neuronal cell body | 4 | Rabbit | Liver | 3.707075E-03 |
| neuronal cell body | 4 | RhesusMacaque | Kidney | 1.012166E-03 |
| myelin sheath | 4 | Chicken | Kidney | 2.817628E-02 |
| myelin sheath | 4 | Human | Brain | 3.646499E-02 |

| GO term | Number of species enriched | Species | Organ | p-value |
|---|---|---|---|---|
| myelin sheath | 4 | Mouse | Brain | 5.339275E-03 |
| myelin sheath | 4 | Mouse | Ovary | 1.545429E-03 |
| myelin sheath | 4 | Opossum | Cerebellum | 2.132704E-03 |
| monocarboxylic acid metabolic process | 4 | Chicken | Brain | 3.924219E-02 |
| monocarboxylic acid metabolic process | 4 | Human | Cerebellum | 1.631572E-02 |
| monocarboxylic acid metabolic process | 4 | Rabbit | Kidney | 8.377301E-05 |
| monocarboxylic acid metabolic process | 4 | Rat | Liver | 5.550880E-04 |
| mitotic nuclear envelope disassembly | 4 | Chicken | Cerebellum | 8.063251E-03 |
| mitotic nuclear envelope disassembly | 4 | Human | Cerebellum | 3.195969E-02 |
| mitotic nuclear envelope disassembly | 4 | Mouse | Brain | 1.780708E-03 |
| mitotic nuclear envelope disassembly | 4 | Rabbit | Brain | 1.294704E-02 |
| mitotic nuclear envelope disassembly | 4 | Rabbit | Kidney | 3.316348E-02 |
| mRNA processing | 4 | Chicken | Cerebellum | 3.810848E-09 |
| mRNA processing | 4 | Chicken | Testis | 8.956988E-04 |
| mRNA processing | 4 | Opossum | Brain | 1.514889E-06 |
| mRNA processing | 4 | Rabbit | Cerebellum | 1.338853E-08 |
| mRNA processing | 4 | Rabbit | Heart | 3.905052E-03 |

| GO term | Number of species enriched | Species | Organ | p-value |
|---|---|---|---|---|
| mRNA processing | 4 | Rabbit | Ovary | 4.279112E-04 |
| mRNA processing | 4 | Rat | Kidney | 2.470946E-02 |
| lipid binding | 4 | Chicken | Brain | 1.384528E-08 |
| lipid binding | 4 | Human | Brain | 4.078347E-02 |
| lipid binding | 4 | Human | Testis | 4.379128E-03 |
| lipid binding | 4 | Opossum | Kidney | 3.217414E-02 |
| lipid binding | 4 | Opossum | Liver | 1.687023E-02 |
| lipid binding | 4 | Opossum | Ovary | 4.976605E-04 |
| lipid binding | 4 | Opossum | Testis | 1.314016E-04 |
| lipid binding | 4 | RhesusMacaque | Brain | 1.176918E-03 |
| histone modification | 4 | Chicken | Kidney | 7.149070E-03 |
| histone modification | 4 | Human | Liver | 2.420050E-02 |
| histone modification | 4 | Opossum | Liver | 4.969648E-03 |
| histone modification | 4 | Rabbit | Cerebellum | 1.577772E-03 |
| growth | 4 | Chicken | Brain | 2.882487E-02 |
| growth | 4 | Chicken | Kidney | 3.249368E-02 |
| growth | 4 | Human | Liver | 3.808167E-02 |
| growth | 4 | Human | Testis | 3.663970E-04 |
| growth | 4 | Opossum | Kidney | 3.602663E-03 |
| growth | 4 | Opossum | Liver | 1.504369E-03 |
| growth | 4 | RhesusMacaque | Kidney | 2.281616E-02 |
| growth | 4 | RhesusMacaque | Liver | 4.593413E-03 |
| cellular response to heat | 4 | Mouse | Brain | 3.137912E-02 |
| cellular response to heat | 4 | Mouse | Heart | 1.033761E-02 |

| GO term | Number of species enriched | Species | Organ | p-value |
|---|---|---|---|---|
| cellular response to heat | 4 | Opossum | Brain | 2.081230E-03 |
| cellular response to heat | 4 | Rabbit | Brain | 3.666139E-04 |
| cellular response to heat | 4 | Rabbit | Heart | 3.434032E-04 |
| cellular response to heat | 4 | Rabbit | Liver | 9.638763E-03 |
| cellular response to heat | 4 | Rabbit | Testis | 1.934104E-02 |
| cellular response to heat | 4 | Rat | Testis | 2.339784E-02 |
| carboxylic acid transport | 4 | Mouse | Cerebellum | 5.733158E-04 |
| carboxylic acid transport | 4 | Rabbit | Brain | 3.364618E-02 |
| carboxylic acid transport | 4 | Rabbit | Kidney | 4.243603E-02 |
| carboxylic acid transport | 4 | Rabbit | Testis | 1.878556E-04 |
| carboxylic acid transport | 4 | Rat | Brain | 1.701845E-04 |
| carboxylic acid transport | 4 | Rat | Kidney | 3.629367E-02 |
| carboxylic acid transport | 4 | RhesusMacaque | Brain | 4.432579E-03 |
| carboxylic acid transport | 4 | RhesusMacaque | Kidney | 2.554235E-04 |
| RNA splicing | 4 | Chicken | Cerebellum | 6.174093E-07 |
| RNA splicing | 4 | Chicken | Testis | 1.768093E-03 |
| RNA splicing | 4 | Opossum | Brain | 5.479311E-06 |
| RNA splicing | 4 | Rabbit | Cerebellum | 8.864233E-06 |
| RNA splicing | 4 | Rabbit | Heart | 2.396241E-02 |

| GO term | Number of species enriched | Species | Organ | p-value |
|---|---|---|---|---|
| RNA splicing | 4 | Rabbit | Ovary | 3.880981E-03 |
| RNA splicing | 4 | Rat | Cerebellum | 4.590026E-02 |
| Golgi membrane | 4 | Human | Brain | 3.888180E-02 |
| Golgi membrane | 4 | Mouse | Liver | 8.361501E-03 |
| Golgi membrane | 4 | Opossum | Liver | 9.752290E-03 |
| Golgi membrane | 4 | Rat | Liver | 4.759188E-02 |
| DNA strand elongation | 4 | Chicken | Cerebellum | 1.375711E-03 |
| DNA strand elongation | 4 | Human | Cerebellum | 3.195969E-02 |
| DNA strand elongation | 4 | Human | Ovary | 1.741919E-02 |
| DNA strand elongation | 4 | Opossum | Brain | 1.931746E-03 |
| DNA strand elongation | 4 | Rabbit | Brain | 1.294704E-02 |
| DNA strand elongation | 4 | Rabbit | Cerebellum | 1.053031E-02 |
| DNA strand elongation | 4 | Rabbit | Heart | 3.129424E-02 |
| DNA strand elongation | 4 | Rabbit | Ovary | 4.811454E-02 |
| DNA recombination | 4 | Chicken | Cerebellum | 2.180369E-05 |
| DNA recombination | 4 | Human | Ovary | 3.484059E-05 |
| DNA recombination | 4 | Opossum | Brain | 3.653805E-03 |
| DNA recombination | 4 | Rabbit | Ovary | 3.838964E-02 |
| spliceosomal complex | 3 | Chicken | Cerebellum | 5.479879E-04 |
| spliceosomal complex | 3 | Chicken | Testis | 2.018537E-02 |

| GO term | Number of species enriched | Species | Organ | p-value |
|---|---|---|---|---|
| spliceosomal complex | 3 | Opossum | Brain | 9.081772E-06 |
| spliceosomal complex | 3 | Rabbit | Cerebellum | 4.317471E-04 |
| spindle | 3 | Chicken | Cerebellum | 1.502310E-02 |
| spindle | 3 | Human | Ovary | 2.320243E-02 |
| spindle | 3 | Mouse | Ovary | 1.337777E-02 |
| regulation of small GTPase mediated signal transduction | 3 | Mouse | Testis | 2.352125E-02 |
| regulation of small GTPase mediated signal transduction | 3 | Opossum | Kidney | 4.010739E-04 |
| regulation of small GTPase mediated signal transduction | 3 | Rat | Brain | 2.999281E-02 |
| regulation of organelle assembly | 3 | Human | Brain | 1.902249E-02 |
| regulation of organelle assembly | 3 | Human | Kidney | 1.525222E-02 |
| regulation of organelle assembly | 3 | Opossum | Kidney | 5.976967E-03 |
| regulation of organelle assembly | 3 | RhesusMacaque | Testis | 4.243652E-03 |
| regulation of neurotransmitter levels | 3 | Mouse | Kidney | 6.157664E-04 |
| regulation of neurotransmitter levels | 3 | Rabbit | Testis | 1.442332E-02 |

| GO term | Number of species enriched | Species | Organ | p-value |
|---|---|---|---|---|
| regulation of neurotransmitter levels | 3 | Rat | Brain | 7.557777E-04 |
| regulation of neuron differentiation | 3 | Human | Kidney | 8.243630E-03 |
| regulation of neuron differentiation | 3 | Human | Liver | 1.528634E-04 |
| regulation of neuron differentiation | 3 | Human | Testis | 3.084980E-02 |
| regulation of neuron differentiation | 3 | Rat | Brain | 1.218651E-03 |
| regulation of neuron differentiation | 3 | RhesusMacaque | Brain | 3.084980E-02 |
| regulation of neurogenesis | 3 | Human | Kidney | 1.332596E-02 |
| regulation of neurogenesis | 3 | Human | Liver | 5.618537E-03 |
| regulation of neurogenesis | 3 | Rat | Brain | 3.260207E-03 |
| regulation of neurogenesis | 3 | RhesusMacaque | Brain | 1.648594E-03 |
| regulation of cellular localization | 3 | Rabbit | Testis | 2.083211E-03 |
| regulation of cellular localization | 3 | Rat | Brain | 2.419833E-02 |
| regulation of cellular localization | 3 | RhesusMacaque | Brain | 2.804325E-02 |
| protein modification by small protein conjugation | 3 | Human | Brain | 2.936493E-02 |

| GO term | Number of species enriched | Species | Organ | p-value |
|---|---|---|---|---|
| protein modification by small protein conjugation | 3 | Opossum | Liver | 8.519610E-03 |
| protein modification by small protein conjugation | 3 | Rabbit | Brain | 4.909767E-03 |
| protein modification by small protein conjugation | 3 | Rabbit | Cerebellum | 2.225951E-03 |
| protein modification by small protein conjugation | 3 | Rabbit | Heart | 2.886272E-02 |
| protein modification by small protein conjugation | 3 | Rabbit | Liver | 4.261986E-03 |
| postsynaptic membrane | 3 | Rabbit | Brain | 4.024320E-02 |
| postsynaptic membrane | 3 | Rat | Brain | 1.381961E-02 |
| postsynaptic membrane | 3 | RhesusMacaque | Brain | 7.374446E-03 |
| positive regulation of neuron projection development | 3 | Human | Kidney | 1.744360E-02 |
| positive regulation of neuron projection development | 3 | Human | Liver | 3.624315E-03 |
| positive regulation of neuron projection development | 3 | Rat | Brain | 5.465396E-04 |

| GO term | Number of species enriched | Species | Organ | p-value |
|---|---|---|---|---|
| positive regulation of neuron projection development | 3 | RhesusMacaque | Brain | 3.735939E-03 |
| positive regulation of neuron projection development | 3 | RhesusMacaque | Liver | 1.446244E-02 |
| positive regulation of cellular component biogenesis | 3 | Human | Brain | 3.252327E-02 |
| positive regulation of cellular component biogenesis | 3 | Opossum | Kidney | 4.857377E-02 |
| positive regulation of cellular component biogenesis | 3 | RhesusMacaque | Kidney | 6.621219E-03 |
| nuclear pore organization | 3 | Chicken | Cerebellum | 2.305939E-02 |
| nuclear pore organization | 3 | Mouse | Ovary | 2.399477E-02 |
| nuclear pore organization | 3 | Rabbit | Brain | 4.523089E-03 |
| nuclear pore organization | 3 | Rabbit | Heart | 2.794849E-03 |
| neuron projection extension | 3 | Chicken | Kidney | 1.511249E-03 |
| neuron projection extension | 3 | Human | Liver | 1.092840E-02 |
| neuron projection extension | 3 | Human | Ovary | 4.462184E-03 |

| GO term | Number of species enriched | Species | Organ | p-value |
|---|---|---|---|---|
| neuron projection extension | 3 | Human | Testis | 1.468632E-02 |
| neuron projection extension | 3 | Mouse | Ovary | 2.976538E-02 |
| negative regulation of DNA metabolic process | 3 | Chicken | Cerebellum | 1.148586E-04 |
| negative regulation of DNA metabolic process | 3 | Chicken | Kidney | 1.119814E-02 |
| negative regulation of DNA metabolic process | 3 | Rabbit | Brain | 3.929867E-02 |
| negative regulation of DNA metabolic process | 3 | Rabbit | Ovary | 3.164396E-04 |
| negative regulation of DNA metabolic process | 3 | Rat | Cerebellum | 4.411583E-03 |
| multicellular organism growth | 3 | Mouse | Heart | 4.011279E-02 |
| multicellular organism growth | 3 | Opossum | Kidney | 2.268281E-04 |
| multicellular organism growth | 3 | Opossum | Liver | 3.589663E-02 |
| multicellular organism growth | 3 | Rat | Testis | 2.341089E-02 |
| mRNA splicing, via spliceosome | 3 | Chicken | Cerebellum | 3.002570E-04 |

| GO term | Number of species enriched | Species | Organ | p-value |
|---|---|---|---|---|
| mRNA splicing, via spliceosome | 3 | Opossum | Brain | 8.976118E-05 |
| mRNA splicing, via spliceosome | 3 | Rabbit | Cerebellum | 2.265463E-04 |
| mRNA splicing, via spliceosome | 3 | Rabbit | Ovary | 1.323936E-03 |
| mRNA metabolic process | 3 | Chicken | Cerebellum | 2.942917E-09 |
| mRNA metabolic process | 3 | Opossum | Brain | 1.713945E-03 |
| mRNA metabolic process | 3 | Rabbit | Cerebellum | 2.419446E-06 |
| locomotion | 3 | Chicken | Brain | 7.007859E-03 |
| locomotion | 3 | Opossum | Kidney | 3.012356E-03 |
| locomotion | 3 | Rat | Brain | 5.973536E-03 |
| lipid modification | 3 | Human | Cerebellum | 3.516033E-02 |
| lipid modification | 3 | Rabbit | Liver | 4.747955E-02 |
| lipid modification | 3 | RhesusMacaque | Cerebellum | 1.685080E-02 |
| dendrite | 3 | Human | Brain | 1.071233E-02 |
| dendrite | 3 | Human | Testis | 4.628626E-02 |
| dendrite | 3 | Rat | Cerebellum | 3.158638E-02 |
| dendrite | 3 | RhesusMacaque | Brain | 9.049240E-03 |
| dendrite | 3 | RhesusMacaque | Kidney | 1.667605E-02 |
| chromatin | 3 | Chicken | Cerebellum | 3.252911E-02 |
| chromatin | 3 | Opossum | Brain | 4.940859E-03 |
| chromatin | 3 | Rabbit | Cerebellum | 2.650926E-02 |
| cell-cell junction | 3 | Chicken | Brain | 1.859775E-02 |

| GO term | Number of species enriched | Species | Organ | p-value |
|---|---|---|---|---|
| cell-cell junction | 3 | Human | Liver | 4.201467E-02 |
| cell-cell junction | 3 | Opossum | Kidney | 1.352189E-02 |
| cell motility | 3 | Chicken | Brain | 2.353436E-03 |
| cell motility | 3 | Opossum | Kidney | 3.415647E-03 |
| cell motility | 3 | Rat | Brain | 5.219345E-04 |
| cell morphogenesis | 3 | Chicken | Brain | 8.667244E-03 |
| cell morphogenesis | 3 | Human | Liver | 2.886974E-02 |
| cell morphogenesis | 3 | Opossum | Kidney | 4.496856E-03 |
| cell migration | 3 | Chicken | Brain | 8.242769E-03 |
| cell migration | 3 | Opossum | Kidney | 6.892123E-03 |
| cell migration | 3 | Rat | Brain | 1.251858E-02 |
| axon extension | 3 | Chicken | Kidney | 3.577275E-03 |
| axon extension | 3 | Human | Ovary | 1.388160E-02 |
| axon extension | 3 | Mouse | Ovary | 1.962823E-02 |
| amino acid transport | 3 | Mouse | Cerebellum | 2.202301E-02 |
| amino acid transport | 3 | Rabbit | Testis | 3.180584E-03 |
| amino acid transport | 3 | Rat | Brain | 1.129950E-03 |
| actin cytoskeleton organization | 3 | Human | Brain | 3.213910E-02 |
| actin cytoskeleton organization | 3 | Human | Liver | 3.134432E-02 |
| actin cytoskeleton organization | 3 | Mouse | Testis | 2.714883E-02 |
| actin cytoskeleton organization | 3 | RhesusMacaque | Kidney | 5.459866E-04 |

| GO term | Number of species enriched | Species | Organ | p-value |
|---|---|---|---|---|
| RNA transport | 3 | Chicken | Cerebellum | 1.892417E-03 |
| RNA transport | 3 | Opossum | Brain | 2.354694E-02 |
| RNA transport | 3 | Rabbit | Cerebellum | 1.035762E-02 |
| RNA splicing, via transesterification reactions | 3 | Chicken | Cerebellum | 4.229494E-05 |
| RNA splicing, via transesterification reactions | 3 | Opossum | Brain | 3.064477E-05 |
| RNA splicing, via transesterification reactions | 3 | Rabbit | Cerebellum | 3.792242E-05 |
| RNA splicing, via transesterification reactions | 3 | Rabbit | Ovary | 6.636840E-04 |
| RNA binding | 3 | Chicken | Cerebellum | 3.351550E-06 |
| RNA binding | 3 | Opossum | Brain | 3.763194E-05 |
| RNA binding | 3 | Rabbit | Cerebellum | 1.407857E-04 |
| RNA binding | 3 | Rabbit | Ovary | 9.406953E-03 |
| Golgi apparatus | 3 | Mouse | Testis | 2.088591E-02 |
| Golgi apparatus | 3 | Opossum | Kidney | 4.728863E-06 |
| Golgi apparatus | 3 | Opossum | Liver | 2.499838E-03 |
| Golgi apparatus | 3 | RhesusMacaque | Brain | 1.790525E-03 |
| GTPase regulator activity | 3 | Mouse | Testis | 4.052479E-02 |
| GTPase regulator activity | 3 | Opossum | Kidney | 1.265053E-04 |

| GO term | Number of species enriched | Species | Organ | p-value |
|---|---|---|---|---|
| GTPase regulator activity | 3 | Rat | Brain | 2.248733E-03 |
| GTPase activator activity | 3 | Mouse | Testis | 2.348025E-02 |
| GTPase activator activity | 3 | Opossum | Kidney | 9.435763E-05 |
| GTPase activator activity | 3 | Rat | Brain | 4.230430E-03 |
| DNA damage checkpoint | 3 | Chicken | Ovary | 3.574349E-02 |
| DNA damage checkpoint | 3 | Human | Ovary | 2.045123E-02 |
| DNA damage checkpoint | 3 | Opossum | Brain | 4.606226E-02 |
| DNA conformation change | 3 | Chicken | Cerebellum | 8.779344E-05 |
| DNA conformation change | 3 | Human | Cerebellum | 4.889852E-02 |
| DNA conformation change | 3 | Human | Ovary | 1.092992E-02 |
| DNA conformation change | 3 | Opossum | Brain | 1.952474E-02 |
| vesicle organization | 2 | Mouse | Liver | 4.802849E-02 |
| vesicle organization | 2 | Opossum | Kidney | 2.550075E-02 |
| ubiquitin-protein transferase activity | 2 | Human | Kidney | 3.407428E-03 |
| ubiquitin-protein transferase activity | 2 | Opossum | Kidney | 5.604345E-03 |

| GO term | Number of species enriched | Species | Organ | p-value |
|---|---|---|---|---|
| ubiquitin-like protein transferase activity | 2 | Human | Kidney | 9.557895E-04 |
| ubiquitin-like protein transferase activity | 2 | Opossum | Kidney | 5.718872E-03 |
| synapse | 2 | Human | Liver | 3.438404E-02 |
| synapse | 2 | RhesusMacaque | Brain | 1.118481E-02 |
| response to organic cyclic compound | 2 | Mouse | Brain | 2.802500E-02 |
| response to organic cyclic compound | 2 | RhesusMacaque | Brain | 4.792339E-02 |
| response to drug | 2 | Mouse | Brain | 1.834866E-04 |
| response to drug | 2 | Mouse | Kidney | 6.629219E-03 |
| response to drug | 2 | Rabbit | Brain | 1.778417E-02 |
| response to drug | 2 | Rabbit | Testis | 8.013147E-03 |
| regulation of wound healing | 2 | Chicken | Brain | 5.929254E-03 |
| regulation of wound healing | 2 | Rat | Ovary | 2.648778E-02 |
| regulation of chromosome segregation | 2 | Chicken | Cerebellum | 1.003355E-02 |
| regulation of chromosome segregation | 2 | Opossum | Brain | 1.407687E-03 |
| regulation of chromosome organization | 2 | Chicken | Cerebellum | 1.925993E-03 |
| regulation of chromosome organization | 2 | Opossum | Brain | 1.989828E-05 |

| GO term | Number of species enriched | Species | Organ | p-value |
|---|---|---|---|---|
| regulation of body fluid levels | 2 | Chicken | Brain | 1.365312E-06 |
| regulation of body fluid levels | 2 | Opossum | Kidney | 1.807120E-03 |
| regulation of body fluid levels | 2 | Opossum | Ovary | 1.610921E-03 |
| regulation of axonogenesis | 2 | Human | Kidney | 4.615258E-03 |
| regulation of axonogenesis | 2 | Human | Liver | 4.903100E-04 |
| regulation of axonogenesis | 2 | RhesusMacaque | Brain | 3.511341E-02 |
| regulation of axonogenesis | 2 | RhesusMacaque | Kidney | 1.493680E-02 |
| regulation of MAP kinase activity | 2 | Chicken | Brain | 1.016098E-03 |
| regulation of MAP kinase activity | 2 | RhesusMacaque | Brain | 2.129519E-02 |
| protein-containing complex binding | 2 | Human | Brain | 4.666854E-02 |
| protein-containing complex binding | 2 | Rat | Cerebellum | 1.333141E-02 |
| protein sumoylation | 2 | Opossum | Brain | 1.144752E-02 |
| protein sumoylation | 2 | Rabbit | Brain | 2.640191E-02 |
| protein sumoylation | 2 | Rabbit | Cerebellum | 1.936006E-02 |

| GO term | Number of species enriched | Species | Organ | p-value |
|---|---|---|---|---|
| proteasome-mediated ubiquitin-dependent protein catabolic process | 2 | Opossum | Liver | 8.436892E-03 |
| proteasome-mediated ubiquitin-dependent protein catabolic process | 2 | Rabbit | Heart | 3.674426E-03 |
| proteasomal protein catabolic process | 2 | Opossum | Liver | 3.602501E-02 |
| proteasomal protein catabolic process | 2 | Rabbit | Heart | 1.540827E-02 |
| postreplication repair | 2 | Mouse | Brain | 1.344836E-02 |
| postreplication repair | 2 | Rabbit | Brain | 1.213290E-02 |
| postreplication repair | 2 | Rabbit | Ovary | 1.353701E-02 |
| phosphatase binding | 2 | Chicken | Brain | 1.753039E-02 |
| phosphatase binding | 2 | Opossum | Kidney | 1.148695E-02 |
| perinuclear region of cytoplasm | 2 | Mouse | Ovary | 2.395777E-02 |
| perinuclear region of cytoplasm | 2 | Opossum | Ovary | 1.906315E-02 |
| organic anion transport | 2 | Rat | Brain | 2.077065E-02 |
| organic anion transport | 2 | RhesusMacaque | Kidney | 2.177027E-02 |

| GO term | Number of species enriched | Species | Organ | p-value |
|---|---|---|---|---|
| organic acid transmembrane transport | 2 | Mouse | Cerebellum | 2.596473E-02 |
| organic acid transmembrane transport | 2 | Rabbit | Ovary | 1.483688E-02 |
| organic acid transmembrane transport | 2 | Rabbit | Testis | 1.679064E-02 |
| nucleotide-excision repair | 2 | Human | Ovary | 4.340058E-02 |
| nucleotide-excision repair | 2 | Opossum | Brain | 2.081230E-03 |
| nuclear speck | 2 | Rabbit | Cerebellum | 1.563014E-03 |
| nuclear speck | 2 | Rat | Cerebellum | 8.259890E-03 |
| nuclear envelope organization | 2 | Chicken | Cerebellum | 2.907213E-02 |
| nuclear envelope organization | 2 | Mouse | Brain | 2.657573E-02 |
| nuclear chromosome, telomeric region | 2 | Chicken | Cerebellum | 1.949679E-04 |
| nuclear chromosome, telomeric region | 2 | Rabbit | Ovary | 1.531904E-02 |
| negative regulation of mitotic cell cycle phase transition | 2 | Opossum | Brain | 2.015255E-02 |
| negative regulation of mitotic cell cycle phase transition | 2 | Rabbit | Cerebellum | 4.317471E-04 |

| GO term | Number of species enriched | Species | Organ | p-value |
|---|---|---|---|---|
| negative regulation of mitotic cell cycle phase transition | 2 | Rabbit | Ovary | 2.310885E-02 |
| negative regulation of mitotic cell cycle | 2 | Mouse | Brain | 1.599066E-03 |
| negative regulation of mitotic cell cycle | 2 | Rabbit | Cerebellum | 1.209612E-05 |
| negative regulation of mitotic cell cycle | 2 | Rabbit | Heart | 9.337593E-03 |
| negative regulation of mitotic cell cycle | 2 | Rabbit | Ovary | 3.686719E-03 |
| negative regulation of chromosome organization | 2 | Chicken | Cerebellum | 6.735525E-04 |
| negative regulation of chromosome organization | 2 | Opossum | Brain | 2.985664E-02 |
| negative regulation of cell proliferation | 2 | Chicken | Brain | 1.583677E-03 |
| negative regulation of cell proliferation | 2 | Rat | Cerebellum | 2.579150E-03 |
| negative regulation of DNA recombination | 2 | Chicken | Cerebellum | 2.538191E-04 |
| negative regulation of DNA recombination | 2 | Rabbit | Brain | 1.736323E-02 |

| GO term | Number of species enriched | Species | Organ | p-value |
|---|---|---|---|---|
| negative regulation of DNA recombination | 2 | Rabbit | Ovary | 4.133854E-02 |
| negative regulation of DNA recombination | 2 | Rabbit | Testis | 2.903244E-02 |
| mRNA transport | 2 | Chicken | Cerebellum | 4.605780E-03 |
| mRNA transport | 2 | Opossum | Brain | 3.438578E-02 |
| hemostasis | 2 | Chicken | Brain | 5.318631E-04 |
| hemostasis | 2 | Opossum | Kidney | 3.748531E-02 |
| focal adhesion | 2 | Chicken | Brain | 1.893044E-04 |
| focal adhesion | 2 | Opossum | Kidney | 5.312457E-04 |
| establishment or maintenance of cell polarity | 2 | Chicken | Kidney | 4.576118E-02 |
| establishment or maintenance of cell polarity | 2 | Opossum | Kidney | 9.100962E-03 |
| enzyme activator activity | 2 | Opossum | Kidney | 9.604041E-03 |
| enzyme activator activity | 2 | Rat | Brain | 4.383202E-03 |
| embryo development | 2 | Human | Liver | 3.479821E-02 |
| embryo development | 2 | Opossum | Kidney | 2.706260E-02 |
| developmental growth involved in morphogenesis | 2 | Chicken | Kidney | 2.154971E-03 |
| developmental growth involved in morphogenesis | 2 | Human | Liver | 3.510049E-02 |

| GO term | Number of species enriched | Species | Organ | p-value |
|---|---|---|---|---|
| developmental growth involved in morphogenesis | 2 | Human | Testis | 2.466846E-02 |
| developmental cell growth | 2 | Chicken | Kidney | 3.635818E-03 |
| developmental cell growth | 2 | Human | Ovary | 4.281663E-02 |
| developmental cell growth | 2 | Human | Testis | 8.740058E-03 |
| ciliary basal body | 2 | Chicken | Ovary | 1.533674E-03 |
| ciliary basal body | 2 | Human | Liver | 2.671951E-02 |
| chromosome, telomeric region | 2 | Chicken | Cerebellum | 1.010889E-04 |
| chromosome, telomeric region | 2 | Chicken | Ovary | 3.406617E-02 |
| chromosome, telomeric region | 2 | Rabbit | Brain | 3.987539E-02 |
| chromosome, telomeric region | 2 | Rabbit | Ovary | 2.164253E-03 |
| chromosome segregation | 2 | Chicken | Cerebellum | 3.034682E-02 |
| chromosome segregation | 2 | Opossum | Brain | 1.496933E-02 |
| chromatin organization | 2 | Chicken | Cerebellum | 8.686863E-03 |
| chromatin organization | 2 | Rabbit | Cerebellum | 1.838031E-03 |
| cellular response to drug | 2 | Mouse | Brain | 5.067055E-03 |
| cellular response to drug | 2 | Rabbit | Brain | 2.786291E-03 |

| GO term | Number of species enriched | Species | Organ | p-value |
|---|---|---|---|---|
| cell-substrate junction | 2 | Chicken | Brain | 4.568550E-04 |
| cell-substrate junction | 2 | Opossum | Kidney | 7.895400E-04 |
| cell-substrate adhesion | 2 | Chicken | Brain | 6.232814E-03 |
| cell-substrate adhesion | 2 | Human | Testis | 1.168177E-02 |
| cell-substrate adherens junction | 2 | Chicken | Brain | 3.103119E-04 |
| cell-substrate adherens junction | 2 | Opossum | Kidney | 9.884145E-04 |
| blood coagulation | 2 | Chicken | Brain | 1.676795E-03 |
| blood coagulation | 2 | Opossum | Kidney | 4.190895E-02 |
| behavior | 2 | Rat | Brain | 4.344032E-04 |
| behavior | 2 | RhesusMacaque | Brain | 1.531830E-03 |
| anion transport | 2 | Rat | Brain | 5.337812E-03 |
| anion transport | 2 | Rat | Kidney | 7.987426E-03 |
| anion transport | 2 | RhesusMacaque | Brain | 2.466117E-02 |
| anion transport | 2 | RhesusMacaque | Kidney | 3.420725E-04 |
| adherens junction | 2 | Chicken | Brain | 3.522330E-04 |
| adherens junction | 2 | Opossum | Kidney | 9.175700E-05 |
| activation of protein kinase activity | 2 | Chicken | Brain | 3.865984E-03 |
| activation of protein kinase activity | 2 | RhesusMacaque | Brain | 1.675064E-02 |
| actin filament-based process | 2 | Human | Brain | 3.646592E-02 |
| actin filament-based process | 2 | Human | Liver | 3.067045E-03 |

| GO term | Number of species enriched | Species | Organ | p-value |
|---|---|---|---|---|
| actin filament-based process | 2 | RhesusMacaque | Kidney | 1.012166E-03 |
| Rab GTPase binding | 2 | Human | Kidney | 1.025391E-02 |
| Rab GTPase binding | 2 | RhesusMacaque | Testis | 1.020743E-02 |
| DNA helicase activity | 2 | Chicken | Cerebellum | 5.314638E-03 |
| DNA helicase activity | 2 | Human | Ovary | 8.848004E-03 |
| DNA geometric change | 2 | Chicken | Cerebellum | 2.963881E-03 |
| DNA geometric change | 2 | Opossum | Brain | 4.351714E-02 |
| DNA duplex unwinding | 2 | Chicken | Cerebellum | 3.143952E-03 |
| DNA duplex unwinding | 2 | Opossum | Brain | 1.442802E-02 |
| viral process | 1 | Rabbit | Cerebellum | 5.756722E-03 |
| viral process | 1 | Rabbit | Ovary | 5.869153E-03 |
| transmembrane transporter activity | 1 | Opossum | Cerebellum | 1.592709E-02 |
| transcription-coupled nucleotide-excision repair | 1 | Opossum | Brain | 3.998113E-02 |
| transcription factor binding | 1 | Rabbit | Cerebellum | 1.116410E-02 |
| tRNA aminoacylation for protein translation | 1 | Mouse | Heart | 2.249169E-02 |
| spindle organization | 1 | Mouse | Testis | 1.618004E-02 |
| spindle checkpoint | 1 | Opossum | Brain | 4.381719E-02 |

| GO term | Number of species enriched | Species | Organ | p-value |
|---|---|---|---|---|
| ribonucleoprotein complex assembly | 1 | Rabbit | Cerebellum | 1.944828E-02 |
| rhythmic process | 1 | Chicken | Cerebellum | 5.001715E-03 |
| rhythmic process | 1 | Chicken | Kidney | 2.092650E-02 |
| response to temperature stimulus | 1 | Rabbit | Brain | 3.485635E-02 |
| response to temperature stimulus | 1 | Rabbit | Heart | 3.722039E-02 |
| response to heat | 1 | Rabbit | Brain | 3.327503E-02 |
| response to heat | 1 | Rabbit | Heart | 2.217615E-02 |
| regulation of secretion | 1 | RhesusMacaque | Brain | 1.056018E-02 |
| regulation of mitotic sister chromatid separation | 1 | Opossum | Brain | 1.170344E-03 |
| regulation of mitotic metaphase/anaphase transition | 1 | Opossum | Brain | 5.497571E-03 |
| regulation of macroautophagy | 1 | Rat | Brain | 4.346992E-02 |
| regulation of mRNA processing | 1 | Opossum | Brain | 2.255842E-02 |
| regulation of inflammatory response | 1 | Chicken | Brain | 5.881530E-03 |
| regulation of growth | 1 | RhesusMacaque | Brain | 4.452923E-02 |
| regulation of establishment of protein localization | 1 | RhesusMacaque | Cerebellum | 2.876542E-02 |

| GO term | Number of species enriched | Species | Organ | p-value |
|---|---|---|---|---|
| regulation of cytoskeleton organization | 1 | Opossum | Kidney | 9.922907E-03 |
| regulation of cell-matrix adhesion | 1 | Opossum | Kidney | 1.441588E-02 |
| regulation of cell projection assembly | 1 | Opossum | Kidney | 3.265345E-02 |
| regulation of cell adhesion | 1 | Chicken | Brain | 2.429477E-03 |
| regulation of autophagy | 1 | Rat | Brain | 7.760611E-03 |
| regulation of Ras protein signal transduction | 1 | Opossum | Kidney | 3.747259E-02 |
| regulation of MAPK cascade | 1 | Chicken | Brain | 2.019300E-02 |
| recombinational repair | 1 | Human | Ovary | 1.115336E-02 |
| rRNA metabolic process | 1 | Chicken | Cerebellum | 2.301971E-02 |
| pyruvate metabolic process | 1 | Human | Testis | 4.102295E-03 |
| protein localization to organelle | 1 | Chicken | Testis | 2.711883E-02 |
| protein lipidation | 1 | RhesusMacaque | Liver | 1.756385E-02 |
| protein import into nucleus | 1 | Rat | Testis | 3.026428E-02 |
| protein domain specific binding | 1 | Chicken | Brain | 3.901064E-03 |
| postsynaptic density | 1 | Rat | Brain | 6.391905E-03 |

| GO term | Number of species enriched | Species | Organ | p-value |
|---|---|---|---|---|
| positive regulation of protein serine/threonine kinase activity | 1 | Chicken | Brain | 1.949764E-02 |
| positive regulation of organelle organization | 1 | Rat | Testis | 1.731070E-02 |
| positive regulation of neuron differentiation | 1 | Rat | Brain | 4.734567E-02 |
| positive regulation of neurogenesis | 1 | RhesusMacaque | Brain | 3.073218E-02 |
| positive regulation of intracellular signal transduction | 1 | Chicken | Brain | 2.386614E-02 |
| positive regulation of developmental growth | 1 | RhesusMacaque | Brain | 7.768747E-03 |
| positive regulation of dendrite development | 1 | Human | Testis | 2.670325E-02 |
| positive regulation of MAP kinase activity | 1 | Chicken | Brain | 2.882487E-02 |
| phosphatidylinositol phosphate binding | 1 | Opossum | Ovary | 2.946450E-02 |
| oxaloacetate metabolic process | 1 | Rat | Liver | 2.577279E-02 |
| organophosphate catabolic process | 1 | Human | Ovary | 2.279573E-02 |
| organelle localization | 1 | Opossum | Kidney | 3.749667E-03 |

| GO term | Number of species enriched | Species | Organ | p-value |
| --- | --- | --- | --- | --- |
| nucleotide catabolic process | 1 | Human | Ovary | 3.140782E-04 |
| nucleotide biosynthetic process | 1 | Rat | Liver | 1.979066E-02 |
| nuclear export | 1 | Opossum | Brain | 1.093775E-03 |
| neutral amino acid transport | 1 | Human | Kidney | 4.854323E-02 |
| neurotrophin TRK receptor signaling pathway | 1 | RhesusMacaque | Brain | 4.397053E-02 |
| neuron recognition | 1 | Human | Liver | 2.863791E-02 |
| negative regulation of intracellular signal transduction | 1 | Mouse | Ovary | 2.509192E-02 |
| negative regulation of cell cycle process | 1 | Rabbit | Cerebellum | 1.609084E-03 |
| negative regulation of cell cycle process | 1 | Rabbit | Ovary | 2.412475E-02 |
| modulation by virus of host process | 1 | Rabbit | Brain | 1.284539E-02 |
| microtubule-based movement | 1 | Chicken | Ovary | 5.992457E-03 |
| microtubule anchoring | 1 | Human | Liver | 4.707232E-02 |
| lyase activity | 1 | Rat | Liver | 4.413795E-03 |
| locomotory behavior | 1 | Rat | Brain | 1.967000E-02 |

| GO term | Number of species enriched | Species | Organ | p-value |
|---|---|---|---|---|
| ion transmembrane transporter activity | 1 | RhesusMacaque | Brain | 8.807251E-03 |
| ion transmembrane transport | 1 | RhesusMacaque | Brain | 2.947754E-02 |
| intraciliary transport | 1 | Chicken | Ovary | 2.775594E-02 |
| import into nucleus | 1 | Rat | Testis | 2.101392E-02 |
| hormone-mediated signaling pathway | 1 | RhesusMacaque | Heart | 3.066218E-02 |
| guanyl-nucleotide exchange factor activity | 1 | Rat | Brain | 1.474891E-02 |
| fascia adherens | 1 | Human | Ovary | 4.489533E-03 |
| extracellular structure organization | 1 | Chicken | Brain | 4.459191E-02 |
| extracellular matrix organization | 1 | Chicken | Brain | 2.312317E-02 |
| extracellular matrix | 1 | Chicken | Brain | 2.470015E-02 |
| exopeptidase activity | 1 | Rat | Liver | 3.113602E-02 |
| exonuclease activity | 1 | Human | Ovary | 3.364209E-02 |
| ephrin receptor signaling pathway | 1 | RhesusMacaque | Kidney | 4.568637E-02 |
| endosome membrane | 1 | Human | Kidney | 2.657674E-02 |
| endoplasmic reticulum membrane | 1 | Mouse | Kidney | 3.916311E-03 |

| GO term | Number of species enriched | Species | Organ | p-value |
|---|---|---|---|---|
| endoplasmic reticulum | 1 | Human | Brain | 3.377920E-02 |
| endoplasmic reticulum | 1 | Human | Cerebellum | 1.857703E-02 |
| drug metabolic process | 1 | Rat | Liver | 9.455105E-03 |
| double-strand break repair via homologous recombination | 1 | Human | Ovary | 9.208850E-03 |
| damaged DNA binding | 1 | Chicken | Cerebellum | 3.931751E-02 |
| dATP metabolic process | 1 | Human | Ovary | 2.324417E-02 |
| core promoter binding | 1 | Chicken | Cerebellum | 9.251330E-03 |
| cofactor metabolic process | 1 | Rat | Liver | 6.342641E-04 |
| coenzyme biosynthetic process | 1 | Rat | Liver | 2.160986E-02 |
| coagulation | 1 | Chicken | Brain | 1.975824E-03 |
| ciliary transition zone | 1 | Human | Liver | 1.180329E-02 |
| chromosome | 1 | Chicken | Cerebellum | 2.027265E-04 |
| chemical homeostasis | 1 | Mouse | Kidney | 3.220091E-03 |
| centriole | 1 | Human | Liver | 1.103620E-02 |
| cellular response to organonitrogen compound | 1 | RhesusMacaque | Liver | 4.212086E-02 |
| cellular response to organic cyclic compound | 1 | Mouse | Brain | 2.294064E-02 |

| GO term | Number of species enriched | Species | Organ | p-value |
|---|---|---|---|---|
| cellular macromolecule catabolic process | 1 | Rabbit | Cerebellum | 3.454484E-02 |
| cellular homeostasis | 1 | Mouse | Kidney | 6.981240E-03 |
| cellular amide metabolic process | 1 | Rat | Liver | 2.820824E-05 |
| cell projection morphogenesis | 1 | Rat | Brain | 3.745976E-02 |
| cell projection membrane | 1 | RhesusMacaque | Kidney | 2.451297E-02 |
| cell morphogenesis involved in differentiation | 1 | Chicken | Brain | 3.828154E-02 |
| cell junction assembly | 1 | Opossum | Kidney | 3.295080E-02 |
| cell growth | 1 | Human | Liver | 1.712347E-02 |
| cell growth | 1 | Human | Testis | 2.712700E-03 |
| cell cycle arrest | 1 | Opossum | Heart | 1.558851E-02 |
| carboxylic acid transmembrane transporter activity | 1 | Rat | Brain | 3.260398E-02 |
| calcium ion binding | 1 | Human | Testis | 3.410092E-03 |
| anion transmembrane transport | 1 | RhesusMacaque | Kidney | 2.829469E-02 |
| anatomical structure formation involved in morphogenesis | 1 | Chicken | Brain | 3.579206E-04 |
| anatomical structure formation involved in morphogenesis | 1 | Chicken | Ovary | 1.984966E-02 |

| GO term | Number of species enriched | Species | Organ | p-value |
|---|---|---|---|---|
| anaphase-promoting complex-dependent catabolic process | 1 | Rabbit | Brain | 1.706179E-02 |
| anaphase-promoting complex-dependent catabolic process | 1 | Rabbit | Cerebellum | 3.555209E-03 |
| anaphase-promoting complex-dependent catabolic process | 1 | Rabbit | Ovary | 2.281093E-03 |
| amino acid transmembrane transporter activity | 1 | Rat | Brain | 4.602119E-02 |
| actin binding | 1 | Opossum | Kidney | 1.705985E-02 |
| SH3 domain binding | 1 | Chicken | Brain | 1.129625E-02 |
| RNA phosphodiester bond hydrolysis, exonucleolytic | 1 | Chicken | Cerebellum | 2.233638E-02 |
| L-amino acid transport | 1 | Rat | Brain | 3.725267E-02 |
| Golgi vesicle transport | 1 | Opossum | Liver | 8.368044E-04 |
| ER to Golgi vesicle-mediated transport | 1 | Mouse | Liver | 1.139301E-02 |
| DNA-templated transcription, termination | 1 | Opossum | Brain | 3.858044E-02 |
| DNA-templated transcription, initiation | 1 | Chicken | Cerebellum | 2.767854E-02 |

| GO term | Number of species enriched | Species | Organ | p-value |
| --- | --- | --- | --- | --- |
| DNA-dependent ATPase activity | 1 | Opossum | Brain | 1.093387E-02 |
| DNA synthesis involved in DNA repair | 1 | Rabbit | Brain | 3.629320E-02 |

**Supplementary Table 2** Enriched processes for gene sets that Sagittarius extrapolates accurately. GO terms enriched by the genes that Sagittarius predicts with at least 0.4 test Pearson correlation comparing timepoints, along with the number of species that the term is enriched for and the enrichment analysis's two-sided Fisher exact test p-value (with Bonferroni correction for multiple hypothesis testing) for each time series.

**Supplementary Table 3**

| Gene | Detailed Explanation |
| --- | --- |
| *PTCH1* | Tumor suppressor gene in the HH signaling pathway, which has been linked to medulloblastoma[1], plexiform fibromyxoma[2], and basal cell carcinoma[3]. Loss-of-function mutations have been documented to lead to aberrant activation of the HH pathway and subsequent tumorigenesis[1]. |
| *MYCBP2* | Encodes a protein that promotes degredation of the MYC oncogene[4], which directly regulates GLI1 in Burkitt lymphoma cell lines[5]. Some studies have found that lymphoblastic leukemia patients had high *c-MYC* expression and low *MYCBP2* expression[6]. |
| *ARID2* | Encodes a protein that directly interacts with *GLI1*[7] as a core subunit of the SWI/SNF chromatin remodeling complex[7, 8]. |
| *PREX1* | Member of the PI3K-Akt signaling pathway, which has been associated with GLI code regulation[9] and cross-talk with the HH signaling pathway in melanoma[3,10]. |
| *DNAH17* | Encodes a protein that makes up a subunit of the primary cilium's basic structure[11]. The primary cilia have been shown to be both positive and negative effectors of the HH signaling pathway[11,12]. |
| *EGFLAM* | Induces activation of the PI3K-Akt signaling pathway[13], which includes *PREX1*. |
| *NLGN1* | Found to be significantly enriched with the HH pathway in a colorectal carcinoma study[14]. |

| Gene | Detailed Explanation |
|---|---|
| **Evidence Sources:** | [1] Lo, W. W., Pinnaduwage, D., Gokgoz, N., Wunder, J. S. & Andrulis, I. L. Aberrant hedgehog signaling and clinical outcome in osteosarcoma. *Sarcoma* **2014**, 261804 (2014).<br>[2] Banerjee, S. *et al.* Loss of the PTCH1 tumor suppressor defines a new subset of plexiform fibromyxoma. *J. Transl. Med.* **17**, 246 (2019).<br>[3] Martinez, M. F. *et al.* Nevoid Basal Cell Carcinoma Syndrome: PTCH1 Mutation Profile and Expression of Genes Involved in the Hedgehog Pathway in Argentinian Patients. *Cells* **8**, (2019).<br>[4] Vatapalli, R. *et al.* Histone methyltransferase DOT1L coordinates AR and MYC stability in prostate cancer. *Nat. Commun.* **11**, 4153 (2020).<br>[5] Yoon, J. W. *et al.* Noncanonical regulation of the Hedgehog mediator GLI1 by c-MYC in Burkitt lymphoma. *Mol. Cancer Res.* **11**, 604–615 (2013).<br>[6] Ge, Z. *et al.* Clinical significance of high c-MYC and low MYCBP2 expression and their association with Ikaros dysfunction in adult acute lymphoblastic leukemia. *Oncotarget* **6**, 42300–42311 (2015).<br>[7] Wang, X., Haswell, J. R. & Roberts, C. W. M. Molecular pathways: SWI/SNF (BAF) complexes are frequently mutated in cancer--mechanisms and potential therapeutic insights. *Clin. Cancer Res.* **20**, 21–27 (2014).<br>[8] Tazzari, M. *et al.* Molecular Determinants of Soft Tissue Sarcoma Immunity: Targets for Immune Intervention. *Int. J. Mol. Sci.* **22**, (2021).<br>[9] Stecca, B. & Ruiz i Altaba, A. Context-dependent regulation of the GLI code in cancer by HEDGEHOG and non-HEDGEHOG signals. *J. Mol. Cell Biol.* **2**, 84–95 (2010).<br>[10] Brechbiel, J., Miller-Moslin, K. & Adjei, A. A. Crosstalk between hedgehog and other signaling pathways as a basis for combination therapies in cancer. *Cancer Treat. Rev.* **40**, 750–759 (2014).<br>[11] Fan, X. *et al.* The association between methylation patterns of DNAH17 and clinicopathological factors in hepatocellular carcinoma. *Cancer Med.* **8**, 337–350 (2019).<br>[12] Hassounah, N. B., Bunch, T. A. & McDermott, K. M. Molecular pathways: the role of primary cilia in cancer progression and therapeutics with a focus on Hedgehog signaling. *Clin. Cancer Res.* **18**, 2429–2435 (2012).<br>[13] Chen, J., Zhang, J., Hong, L. & Zhou, Y. EGFLAM correlates with cell proliferation, migration, invasion and poor prognosis in glioblastoma. *Cancer Biomark.* **24**, 343–350 (2019).<br>[14] Yu, Q. *et al.* Upregulated NLGN1 predicts poor survival in colorectal cancer. *BMC Cancer* **21**, 884 (2021). |

**Supplementary Table 3** Literature evidence connecting Sagittarius's extrapolated mutation profile for early-stage sarcoma patients to the HH signaling pathway and *GLI* oncogene.

#### Supplementary Figures

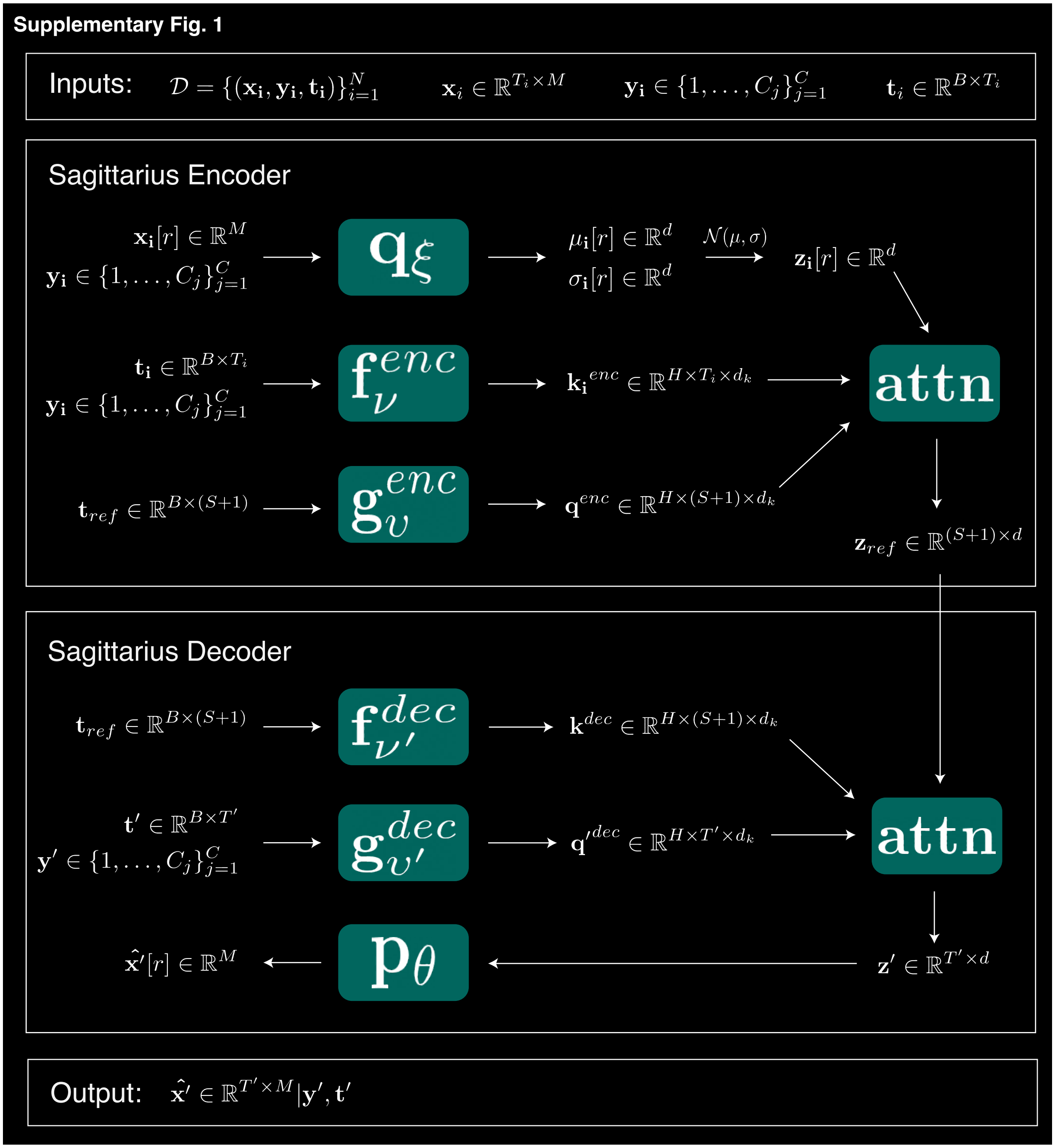

**Supplementary Fig. 1** Flowchart of the overall Sagittarius model workflow. Interaction between main model components, along with labeled input, output, and intermediary result dimensions. The module components are divided into encoder and decoder phases around the shared reference space.

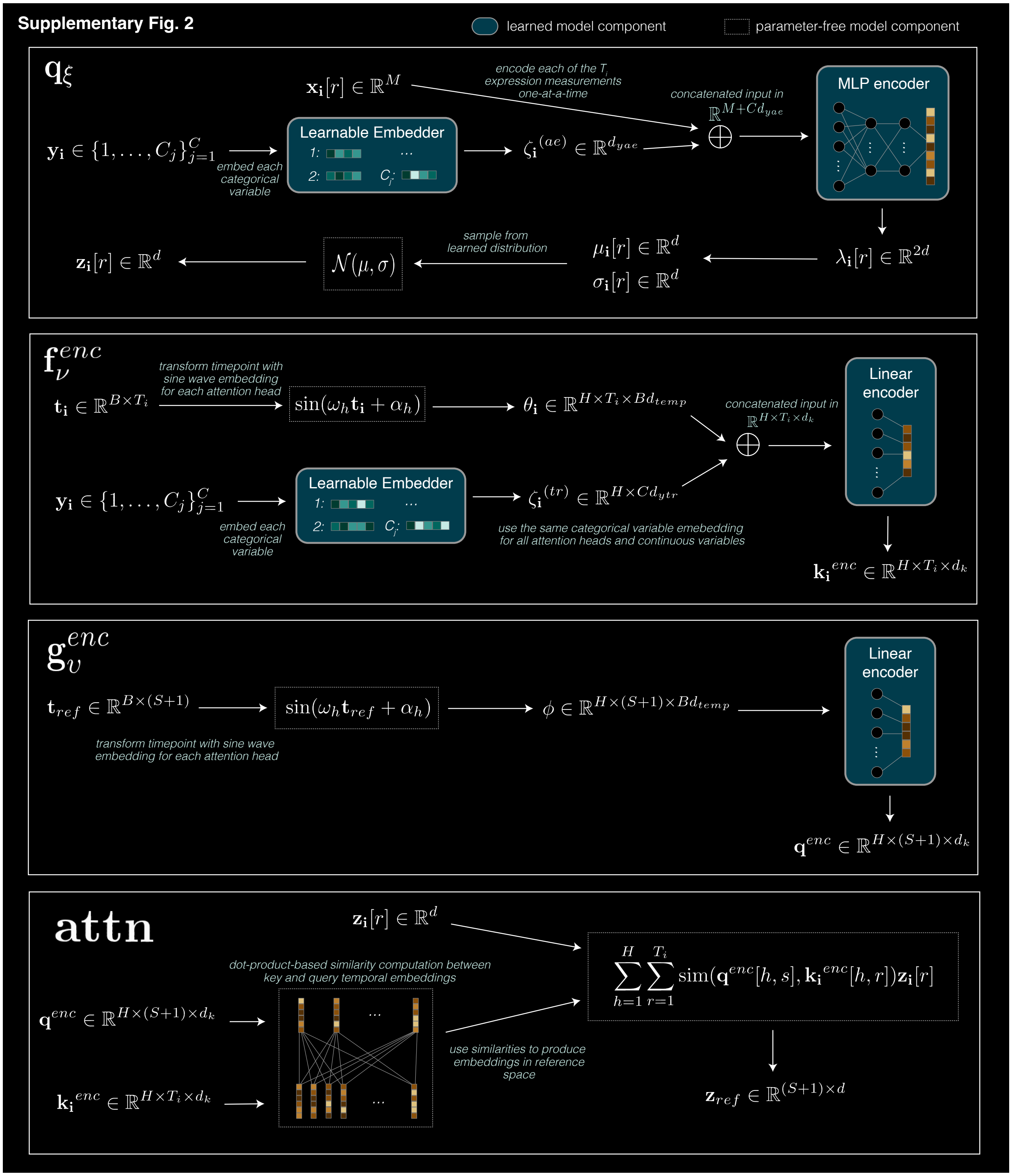

**Supplementary Fig. 2** Flowchart of Sagittarius encoder with model implementation details. Inputs and intermediate results are annotated with their dimensions. The dimensions $\{d, d_{yae}, d_{ytr}, d_{temp}\}$ are all Sagittarius hyperparameters, with $d_k = \sum_{b=1}^{B} d_{temp} + \sum_{c=1}^{C} d_{ytr}$. Model components involving learned parameters are shown in blue boxes; parameter-free components are shown in dotted white boxes.

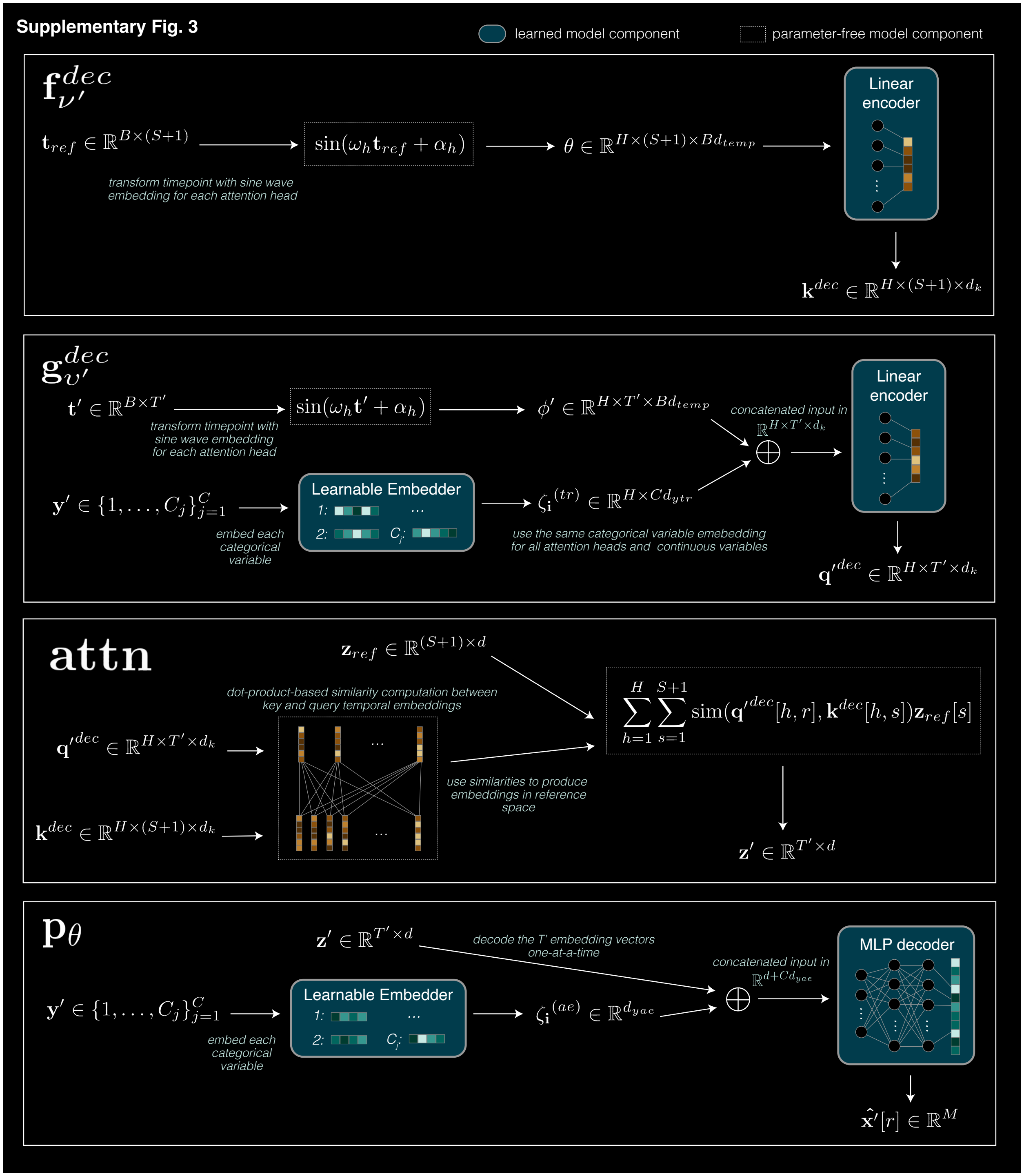

**Supplementary Fig. 3** Flowchart of Sagittarius decoder with model implementation details. Intermediate results and model outputs are annotated with their dimensions. The decoder shares dimension hyperparameters $\{d, d_{yae}, d_{ytr}, d_{temp}\}$ with the encoder module, with $d_k = \sum_{b=1}^{B} d_{temp} + \sum_{c=1}^{C} d_{ytr}$. Model components involving learned parameters are shown in blue boxes; parameter-free components are shown in dotted white boxes.

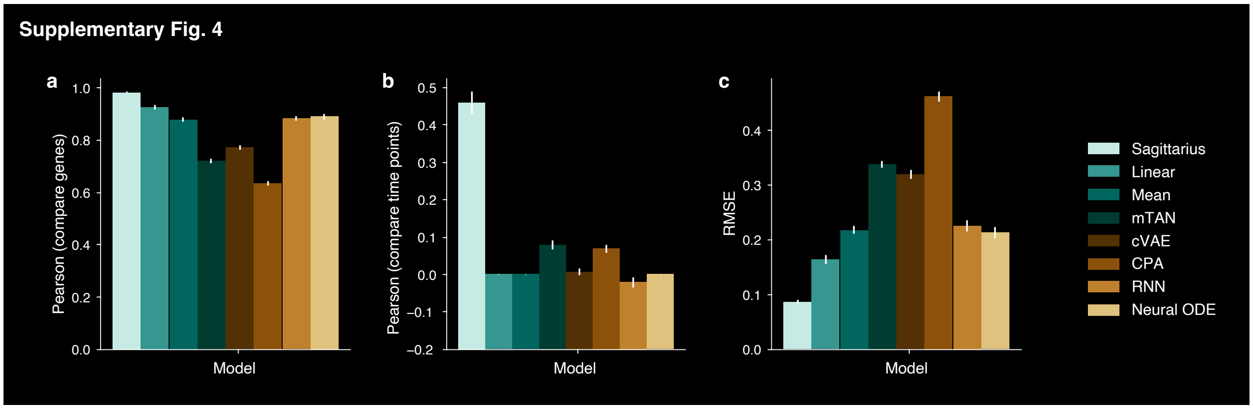

**Supplementary Fig. 4** Overall gene expression prediction performance for Evo-devo extrapolation to late timepoints. **a-c,** Bar plot of Pearson correlation comparing genes (**a**), Pearson correlation comparing timepoints (**b**), and RMSE (**c**) of the extrapolated expression profile and measured expression profile when extrapolating to the last four measured timepoints from each species and organ combination in the Evo-devo dataset for Sagittarius and the comparison approaches. For Pearson correlation, comparing genes or comparing timepoints (**a,b**), higher values indicate better performance; for RMSE (**c**), lower values indicate better performance. Data are presented as mean values +/- standard error, with n=48 species and organ time series.

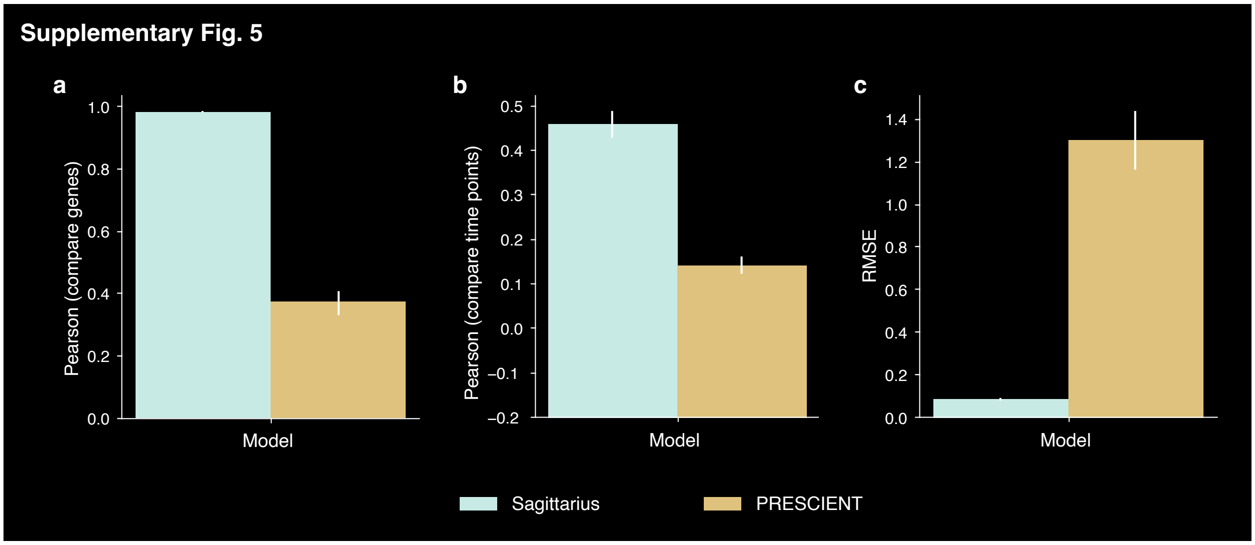

**Supplementary Fig. 5** Comparison of Sagittarius and PRESCIENT models for late timepoint Evo-devo extrapolation performance. **a-c,** Bar plot of Pearson correlation comparing genes (**a**), Pearson correlation comparing timepoints (**b**), and RMSE (**c**) of the predicted expression profile and measured expression profile when extrapolating to the last four measured timepoints from each species and organ combination in the Evo-devo dataset for Sagittarius and the single-cell fate prediction model PRESCIENT. For Pearson correlation (**a,b**), higher values indicate better performance; for RMSE (**c**), lower values indicate better performance. Data are presented as mean values +/- standard error, with n=48 species and organ time series.

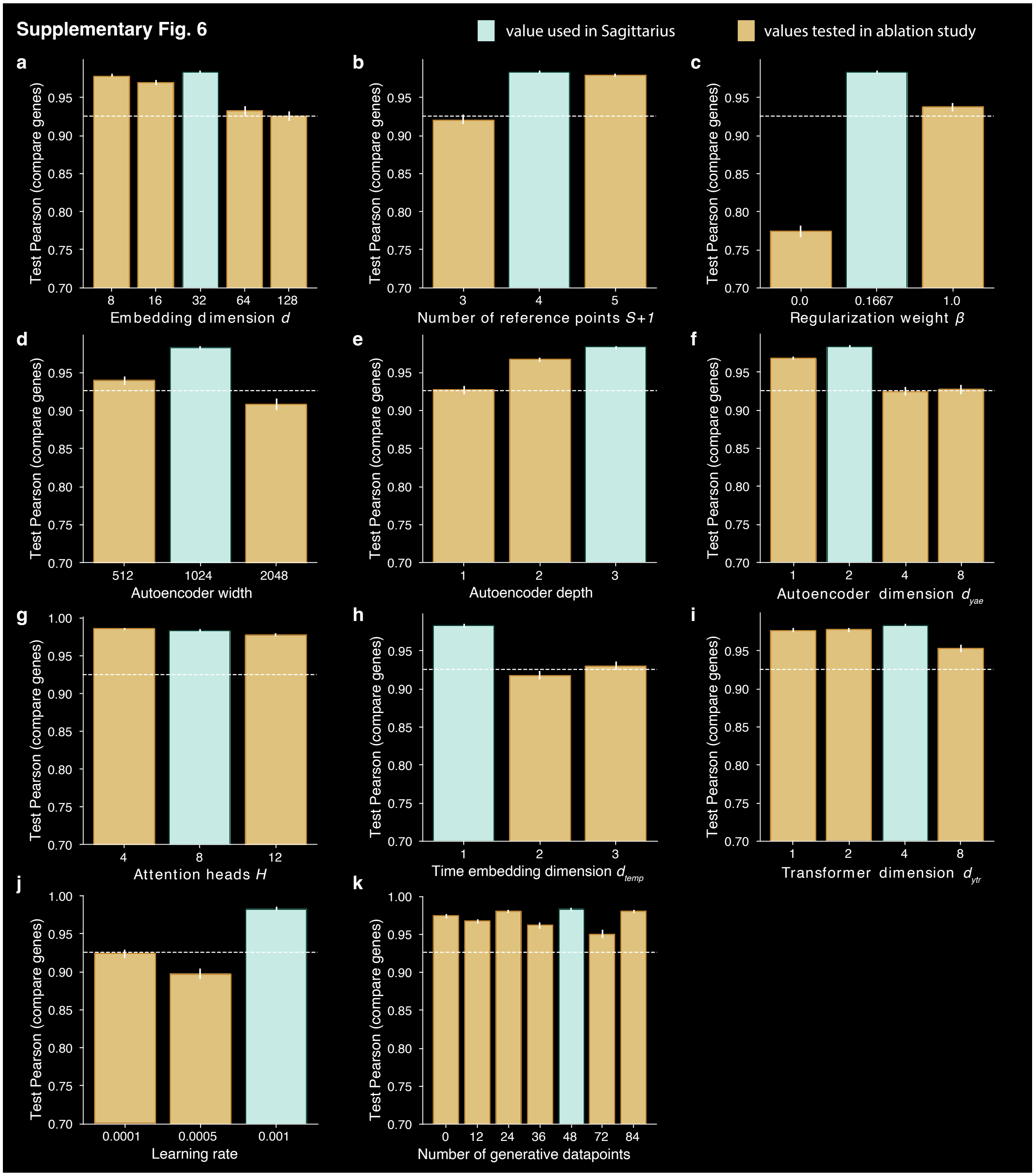

**Supplementary Fig. 6** Test Pearson correlation comparing genes for different choices of Sagittarius hyperparameter settings on the Evo-devo late timepoint extrapolation task. **a-k**, Bar plot comparing Sagittarius performance when holding all but one hyperparameter fixed and varying embedding dimension d (**a**), number of reference points $S + 1$ (**b**), regularization weight (**c**), autoencoder MLP width (**d**), autoencoder MLP depth (**e**), autoencoder categorical embedding dimension $d_{yae}$ (**f**), number of attention heads $H$ (**g**), temporal embedding dimensions $d_{temp}$ (**h**), transformer categorical embedding dimension $d_{ytr}$ (**i**), learning rate (**j**), and number of time series examples for the generative objective while holding the number of reconstructive time series examples fixed at 48 (**k**). The white dotted line shows the Pearson correlation comparing genes of the best-performing baseline for reference. Higher values indicate better performance. Data are presented as mean values +/- standard error, with n=48 species and organ time series.

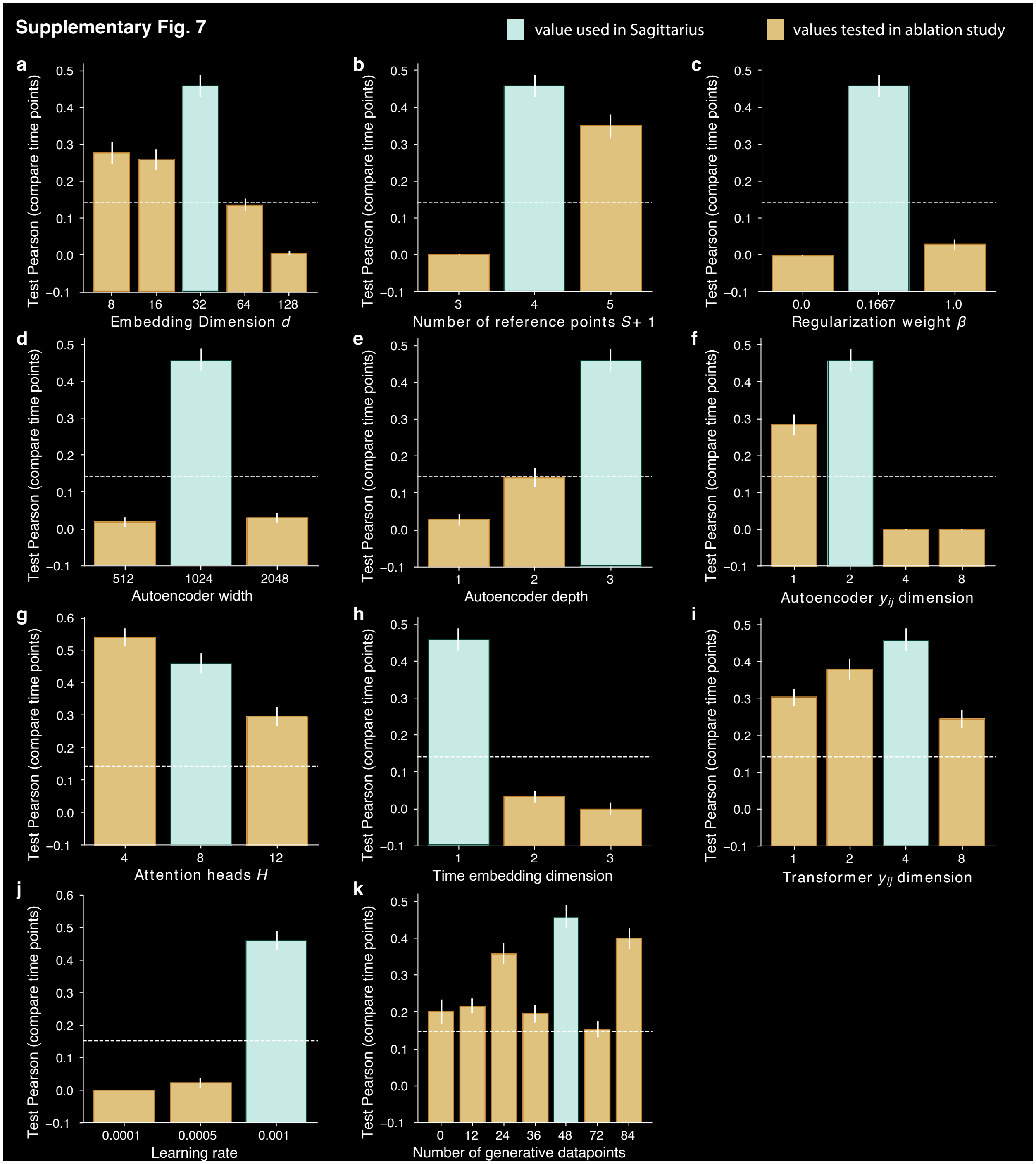

**Supplementary Fig. 7** Test Pearson correlation comparing timepoints for different choices of Sagittarius hyperparameter settings on the Evo-devo late timepoint extrapolation task. **a-k**, Bar plot comparing Sagittarius performance when holding all but one hyperparameter fixed and varying embedding dimension d (**a**), number of reference points $S + 1$ (**b**), regularization weight (**c**), autoencoder MLP width (**d**), autoencoder MLP depth (**e**), autoencoder categorical embedding dimension $d_{yae}$ (**f**), number of attention heads $H$ (**g**), temporal embedding dimensions $d_{temp}$ (**h**), transformer categorical embedding dimension $d_{ytr}$ (**i**), learning rate (**j**), and number of time series examples for the generative objective while holding the number of reconstructive time series examples fixed at 48 (**k**). The white dotted line shows the Pearson correlation comparing timepoints of the best-performing baseline for reference. Higher values indicate better performance. Data are presented as mean values +/- standard error, with n=48 species and organ time series.

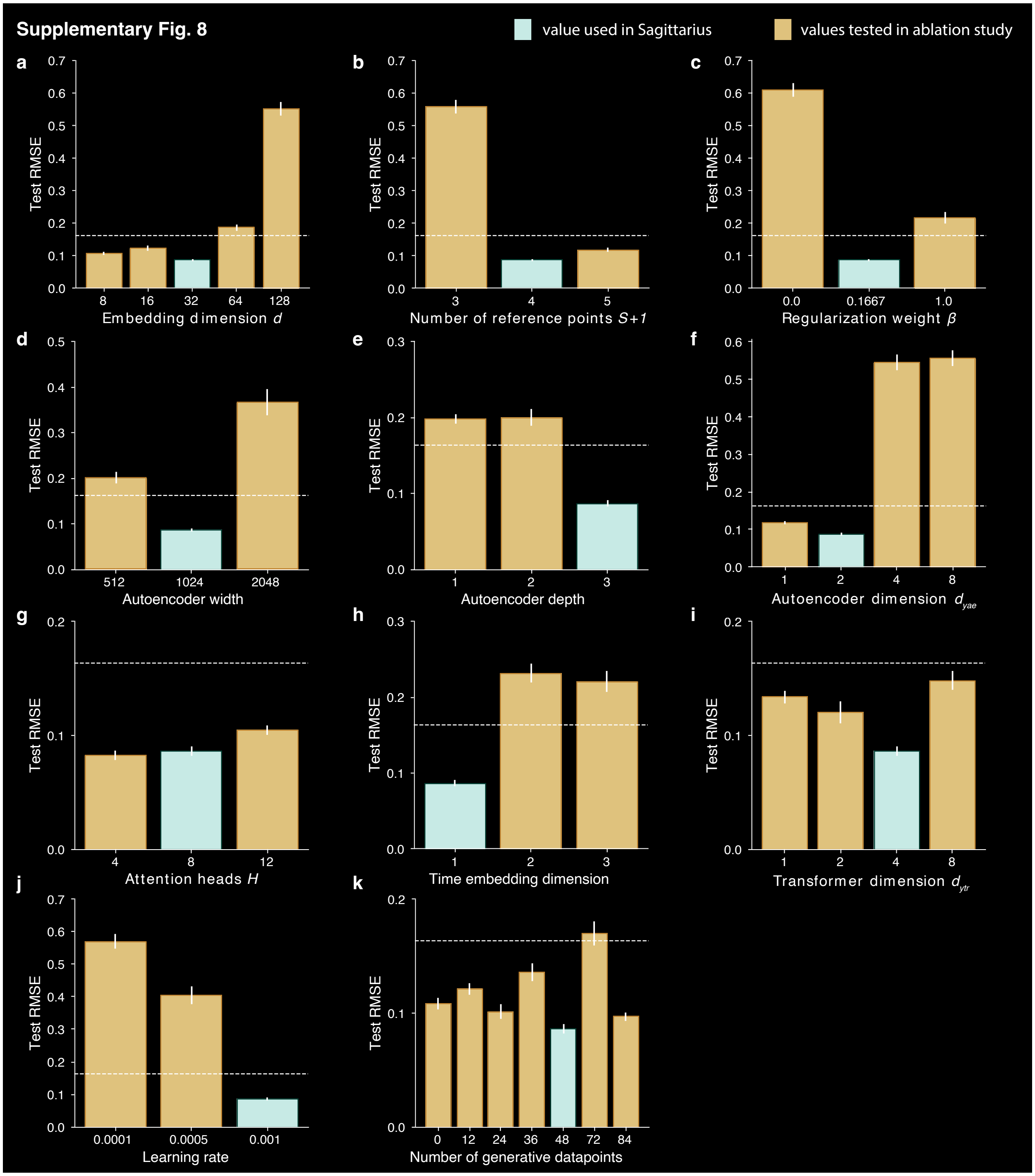

**Supplementary Fig. 8** Test RMSE for different choices of Sagittarius hyperparameter settings on the Evo-devo late timepoint extrapolation task. **a-k**, Bar plot comparing Sagittarius performance when holding all but one hyperparameter fixed and varying embedding dimension d (**a**), number of reference points $S + 1$ (**b**), regularization weight (**c**), autoencoder MLP width (**d**), autoencoder MLP depth I, autoencoder categorical embedding dimension $d_{yae}$ (**f**), number of attention heads $H$ (**g**), temporal embedding dimensions $d_{temp}$ (**h**), transformer categorical embedding dimension

$d_{ytr}$ (**i**), learning rate (**j**), and number of time series examples for the generative objective while holding the number of reconstructive time series examples fixed at 48 (**k**). The white dotted line shows the RMSE of the best-performing baseline for reference. Lower values indicate better performance. Data are presented as mean values +/- standard error, with n=48 species and organ time series.

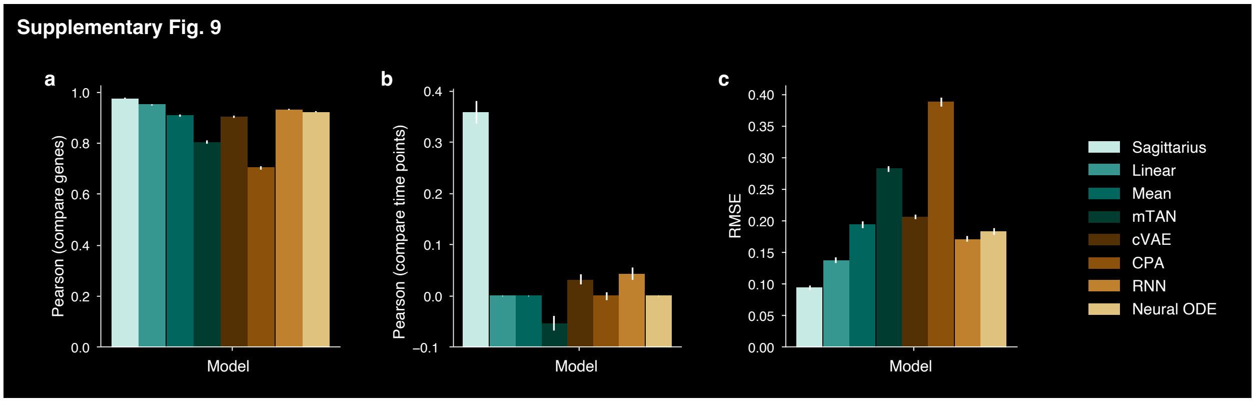

**Supplementary Fig. 9** Overall gene expression extrapolation performance for Evo-devo extrapolation to early timepoints. **a-c**, Bar plot of Pearson correlation comparing genes (**a**), Pearson correlation comparing timepoints (**b**), and RMSE (**c**) of the predicted expression profile and measured expression profile when extrapolating to the first four measured timepoints from each species and organ combination in the Evo-devo dataset for Sagittarius and the comparison approaches. For Pearson correlation, comparing genes or comparing timepoints (**a,b**), higher values indicate better performance; for RMSE (**c**), lower values indicate better performance. Data are presented as mean values +/- standard error, with n=48 species and organ time series.

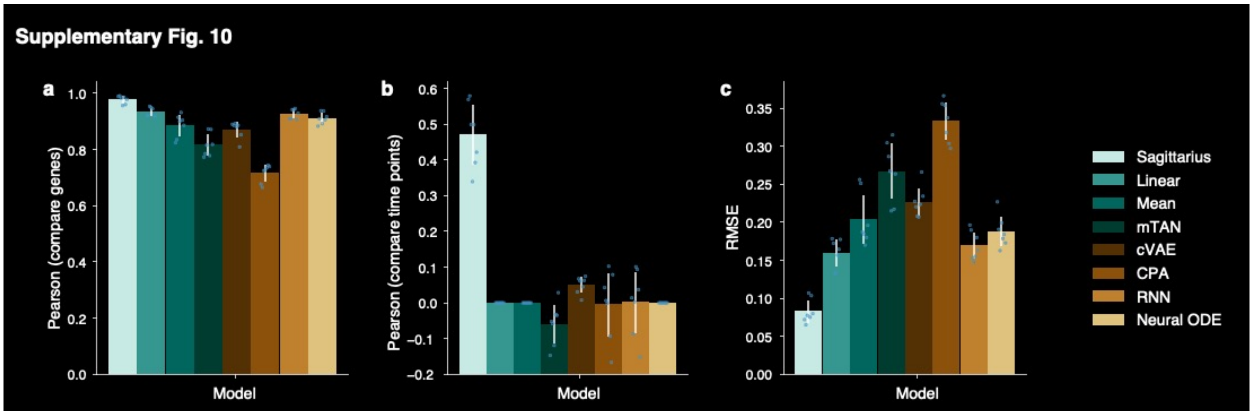

**Supplementary Fig. 10** Human gene expression extrapolation performance for Evo-devo extrapolation performance to early timepoints. **a-c,** Bar plot comparing Sagittarius and existing approaches in terms of Pearson correlation comparing genes (**a**), Pearson correlation comparing timepoints (**b**), and RMSE (**c**) of the predicted human expression profile and measured human expression profile of each organ when extrapolating to the first four measured sequence timepoints in the Evo-devo dataset. For Pearson correlation, comparing genes or comparing timepoints (**a**,**b**), higher values indicate better performance; for RMSE (**c**), lower values indicate better performance. Data are presented as mean values +/- standard error, with n=7 organ time series.

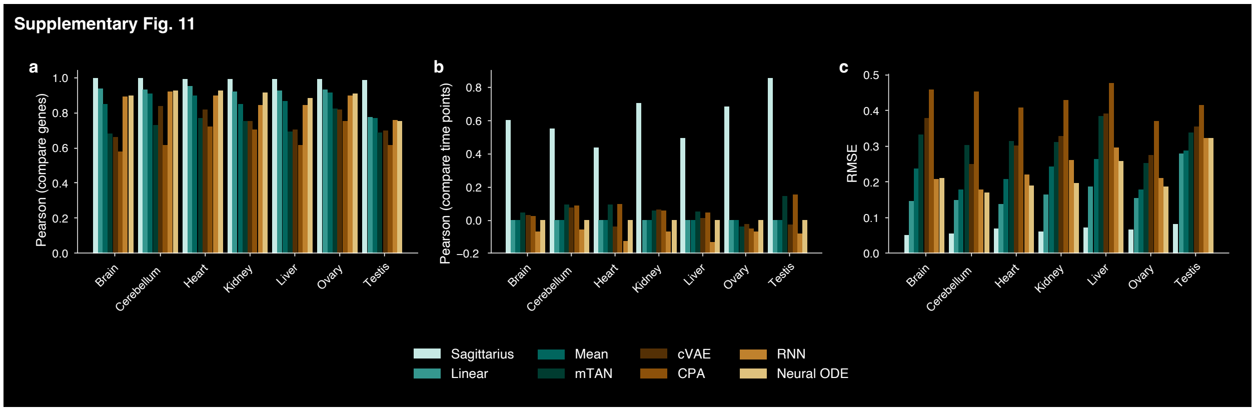

**Supplementary Fig. 11** Mouse gene expression extrapolation performance for Evo-devo extrapolation task to late timepoints. **a-c,** Bar plot comparing Sagittarius and existing approaches in terms of Pearson correlation comparing genes (**a**), Pearson correlation comparing timepoints (**b**), and RMSE (**c**) of the predicted mouse expression profile and measured mouse expression profile of each organ when extrapolating to the final four measured sequence timepoints in the Evo-devo dataset (n=1 time series). For Pearson correlation, comparing genes or comparing timepoints (**a**,**b**), higher values indicate better performance; for RMSE (**c**), lower values indicate better performance.

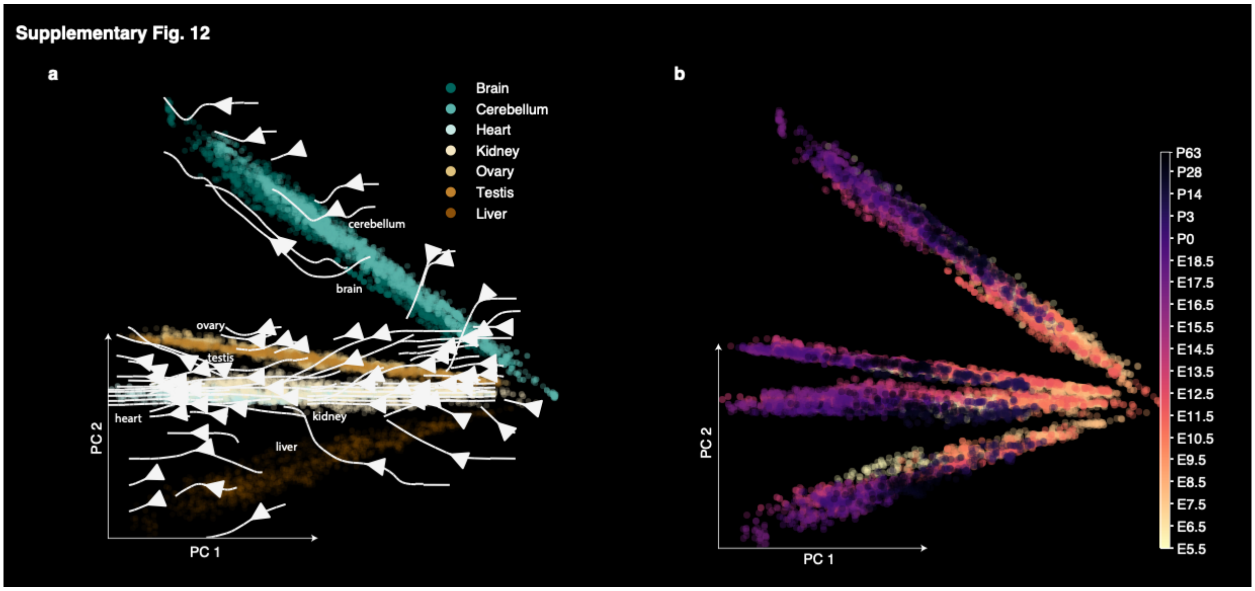

**Supplementary Fig. 12** Mouse transcriptomic velocity across organs. **a,b,** PCA plot showing extrapolated mouse gene expression from E5.5 to P63 for 7 organs, colored by organ (**a**) and time (**b**). The arrows in (**a**) indicate the transcriptomic velocity of each organ. The first PC shows most variation with respect to time, while the second shows most variation with respect to organ. Organ annotations in (**a**) are added to help differentiate between organs, especially in the case of overplotting. An arrow annotation in (**b**) indicates the predicted timepoints that lie outside the range of measured times in the Evo-devo dataset.

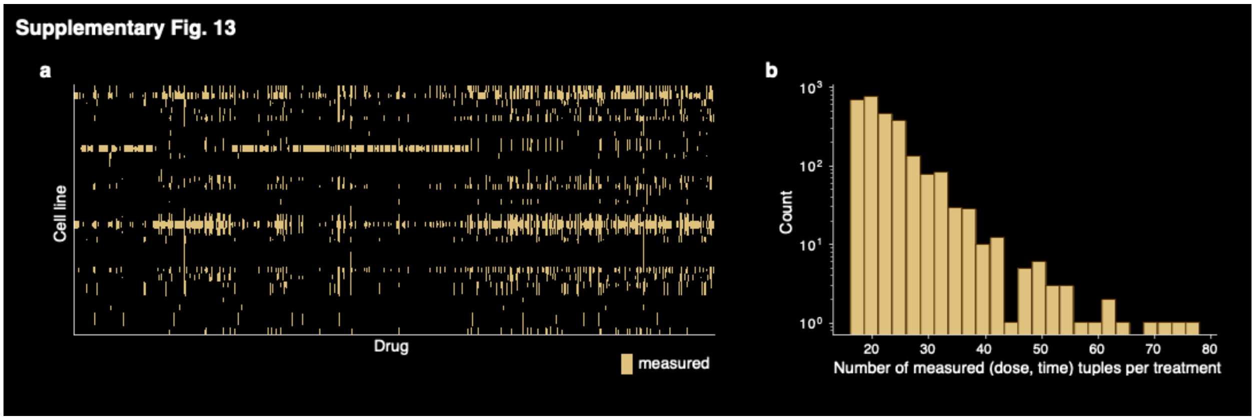

**Supplementary Fig. 13** Time series measured in the restricted LINCS dataset. **a,** Heatmap indicating the drug and cell line combinations that have time series measurements included in the LINCS dataset we use after initial processing. Cell lines tend to be either relatively well-measured or very sparsely measured. **b,** Histogram of the sequence lengths for all measured drug and cell line combinations. The length of the sequence is the number of unique dose and treatment time combinations that the therapeutic combination is measured at.

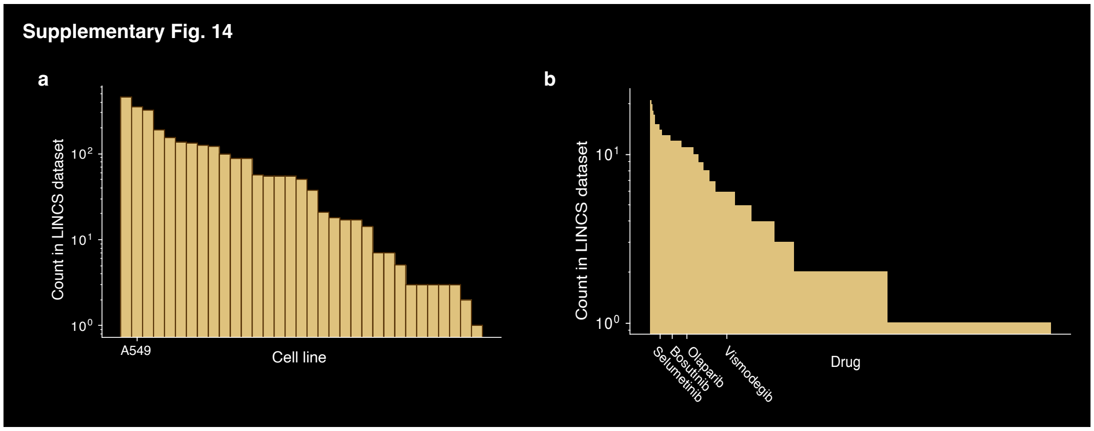

**Supplementary Fig. 14** LINCS measurements with the best-performing cell line and drugs for the $IC_{50}$ prediction task with the GDSC dataset. **a,b,** Histogram of the number of measured drug treatments per cell line (**a**) and cell lines treated per drug (**b**) in the LINCS dataset. The A549 cell line is annotated as the cell line with the most improved predictions from Sagittarius's imputed dataset (**a**). The drugs with the most-improved predictions, Selumetinib, Bosutinib, Olaparib, and Vismodegib, are also annotated (**b**).

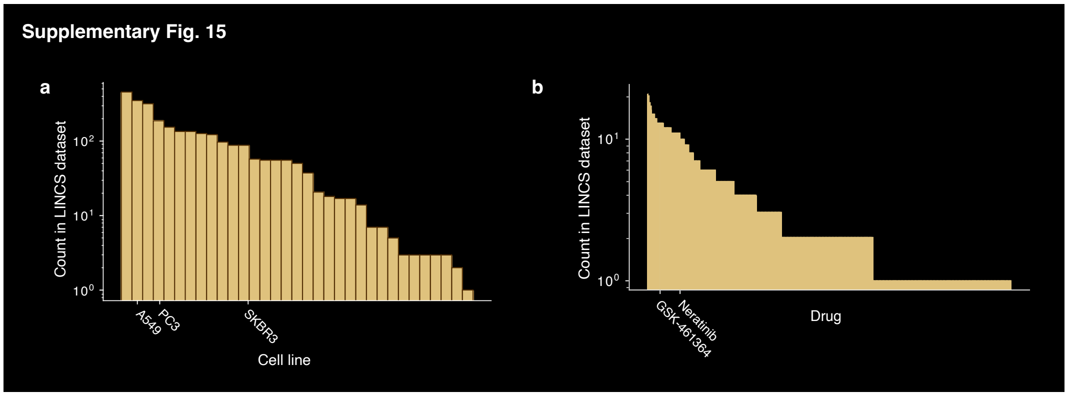

**Supplementary Fig. 15** LINCS measurements with the best-performing cell line and drugs for the $IC_{50}$ prediction task with the CTRP dataset. **a,b,** Histogram of the number of measured drug treatments per cell line (**a**) and cell lines treated per drug (**b**) in the LINCS dataset. A549 and PC3, the cell lines for which Sagittarius's extrapolated data most improves the predictions, are annotated. SKBR3, which Sagittarius struggles on, is also annotated (**a**). GSK-461364 and Neratinib are annotated as the most-improved drugs with Sagittarius's imputed dataset (**b**).

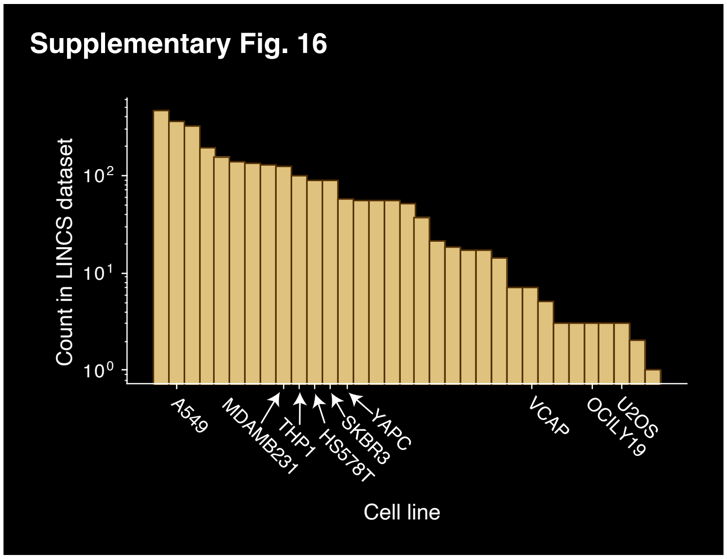

**Supplementary Fig. 16** LINCS measurements with the best-performing cell lines for the gene essentiality prediction task with the DEMETER and CERES DepMap datasets. Histogram of the number of drug treatment experiments measured in the LINCS dataset per cell line. Sagittarius's imputed dataset provided the most benefit are A549, MDAMB231, THP1, HS578T, SKBR3, YAPC, VCAP, OCILY19, and U2OS, which are annotated.

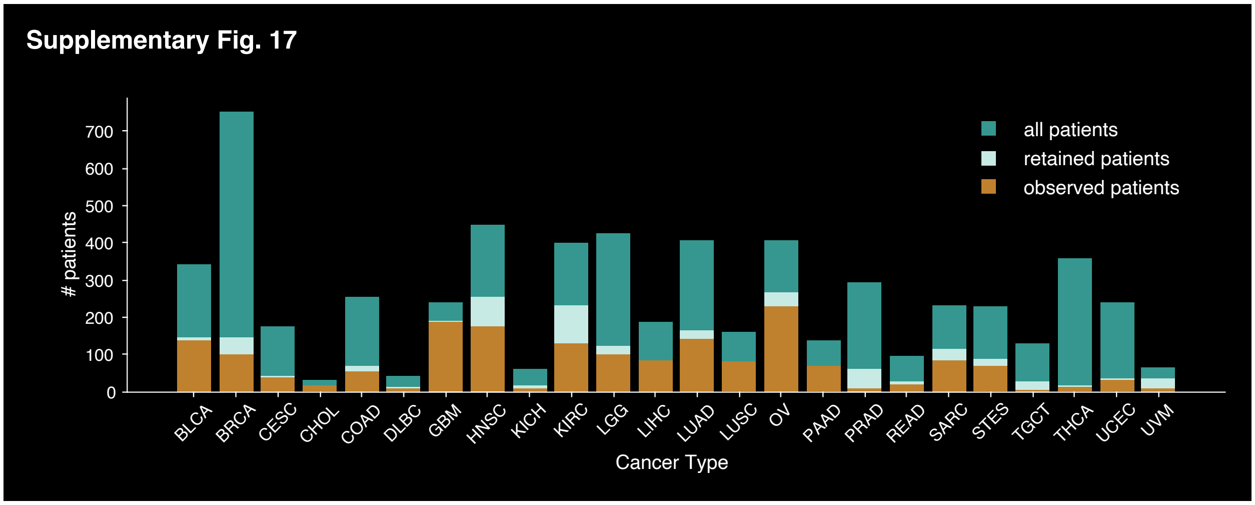

**Supplementary Fig. 17** Distribution of TCGA patients per cancer type in the mutation dataset. Comparison of patient counts if all patients are used in the analysis, patient counts if only retained patients (including all observed patients and some censored patients) are used in the analysis, and patient counts if only observed patients are used in the analysis. By construction, the number of total patients is larger than the number of retained patients, which is in turn at least as large as the number of observed patients. Retaining some censored patients according to the individual survival prediction loss could improve model power without corrupting the time series formulation.

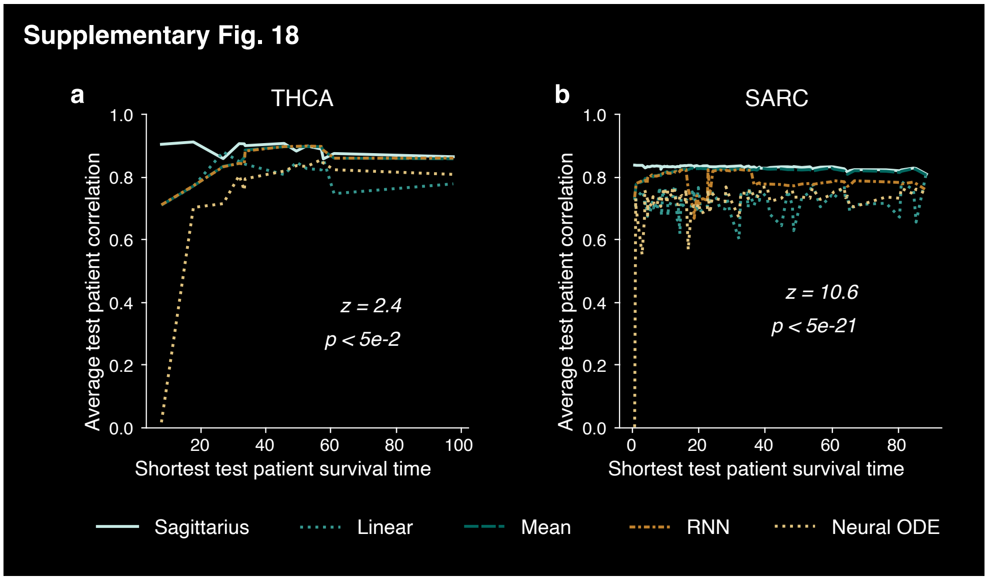

**Supplementary Fig. 18** Performance comparison for cancer patient gene expression extrapolation. **a,b,** Line plot showing average Spearman correlation of predicted expression profile and measured expression profile for THCA (**a**) and SARC (**b**) test splits, ordered according to the shortest survival time in the test set. Annotations indicate the z- and p-value comparing Sagittarius to the best-performing baseline (one-sided Fisher z-transformed test), with p-value = 0.011 for THCA and p-value = 3.92e-21 for SARC.

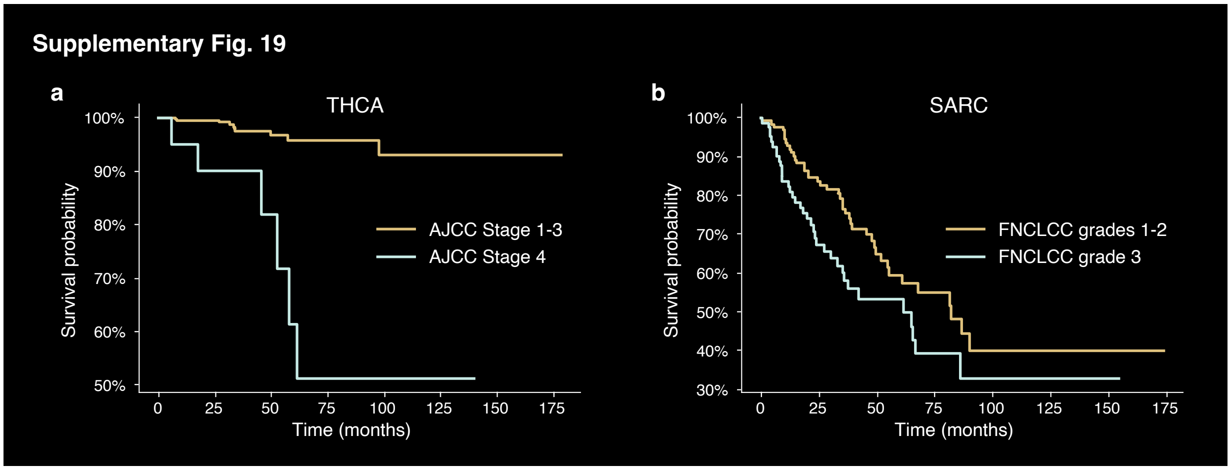

**Supplementary Fig. 19** Comparison of TCGA patient survival for different tumor grades at diagnosis. **a**,**b**, Kaplan-Meier curve showing probability of survival over time for cohorts of THCA (**a**) and SARC (**b**) patients. THCA patients are subdivided according to the American Joint Committee on Cancer (AJCC) stage codes provided by TCGA, with less-severe cancer stages 1, 2, and 3 forming one cohort and the more-severe stage 4 patients in another cohort. Stage 4 THCA patients have worse prognosis than patients diagnosed with stage 1-3 carcinoma (two-sided log rank test, p-value = 1.51e-7). SARC patients are subdivided according to the Fédération Nationale des Centres de Lutte Contre le Cancer (FNCLCC) grade labels provided by TCGA, with less-severe grades 1 and 2 forming one cohort and the more-severe grade 3 patients in another cohort. Grade 3 SARC patients have worse prognosis than grade 1 or 2 sarcoma patients (two-sided log rank test, p-value = 0.027).

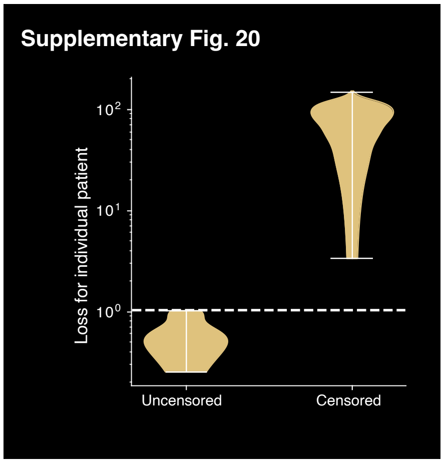

**Supplementary Fig. 20** THCA censored patient analysis. Violin plot of the survival regressor's absolute error for each THCA patient, subdivided into an observed group and a censored group.

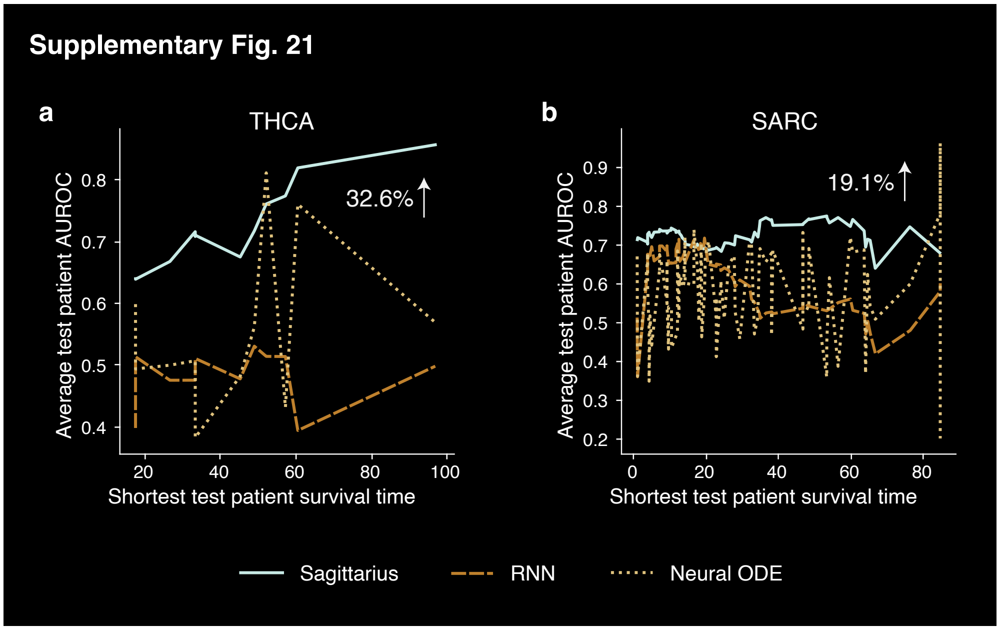

**Supplementary Fig. 21** Patient mutation profile extrapolation performance for Sagittarius and other deep learning models. **a**,**b**, Line plot comparing average test patient AUROC for each of the THCA (**a**) and SARC (**b**) cancer type test splits, ordered according to the shortest survival time in that test set. The annotations indicate Sagittarius's percent AUROC improvement over the best-performing deep learning comparison approach.

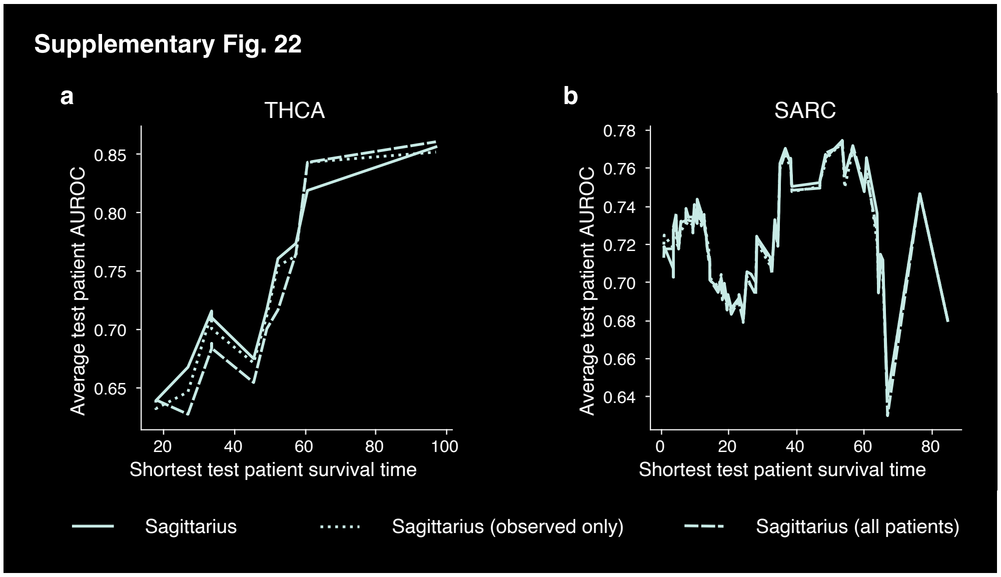

**Supplementary Fig. 22** Comparison of Sagittarius patient extrapolation performance with different inclusion criteria for censored patients. **a**,**b**, Average test patient AUROC for the THCA (**a**) and SARC (**b**) cancer type test splits, ordered by the shortest patient survival time in each test split. The Sagittarius method retains some censored patients; Sagittarius (observed only) excludes all censored patients; Sagittarius (all patients) includes all censored patients.

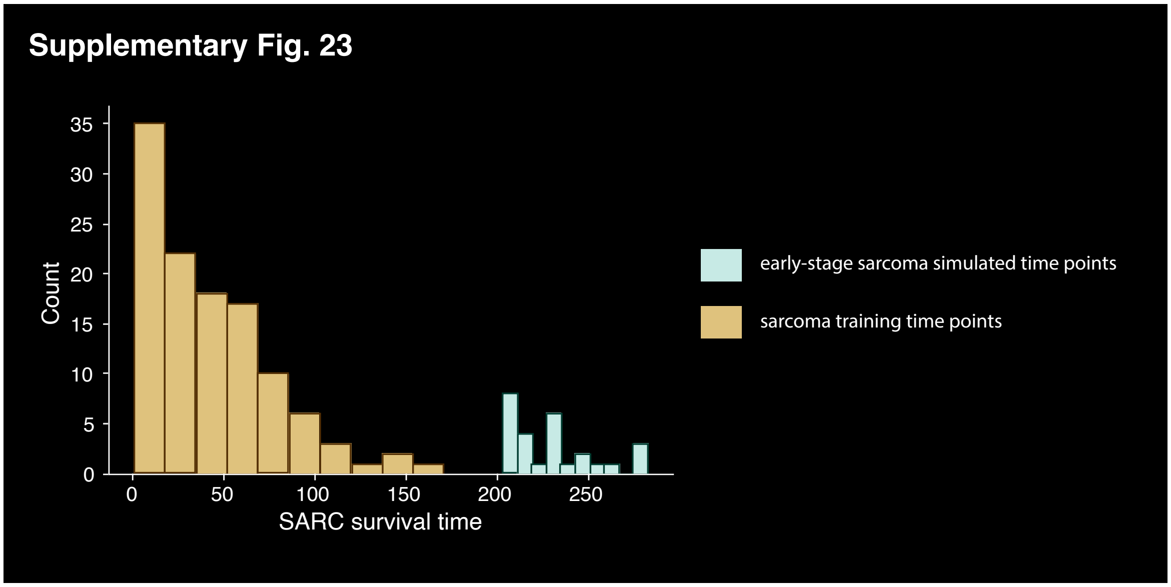

**Supplementary Fig. 23** SARC training and extrapolation timepoint distribution. Histogram showing the measured survival time of patients in the SARC time series as the available sarcoma training data and the extrapolation timepoints used to simulate the expression profile of an early-stage sarcoma patient.

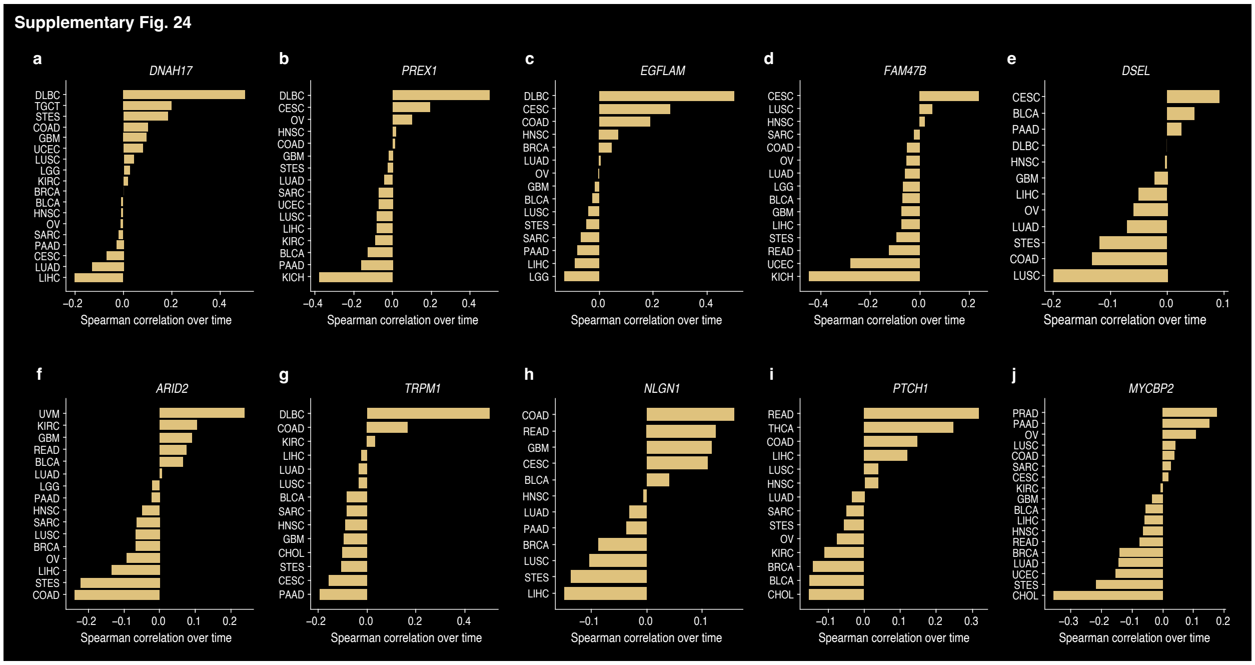

**Supplementary Fig. 24** Normalized mutation rate and survival time for Sagittarius's predicted early-stage sarcoma mutations. **a-j,** Bar plot of the Spearman correlation of survival time and a patient's mutation normalized by their total mutation load for the top-10 predicted mutations in simulated early-stage sarcoma patients, *DNAH17* (**a**), *PREX1* (**b**), *EGFLAM* (**c**), *FAM47B* (**d**), *DSEL* (**e**), *ARID2* (**f**), *TRPM1* (**g**), *NLGN1* (**h**), *PTCH1* (**i**), and *MYCBP2* (**j**). We show the Spearman correlation for each cancer type where at least two patients in the time series have a mutation in the gene.

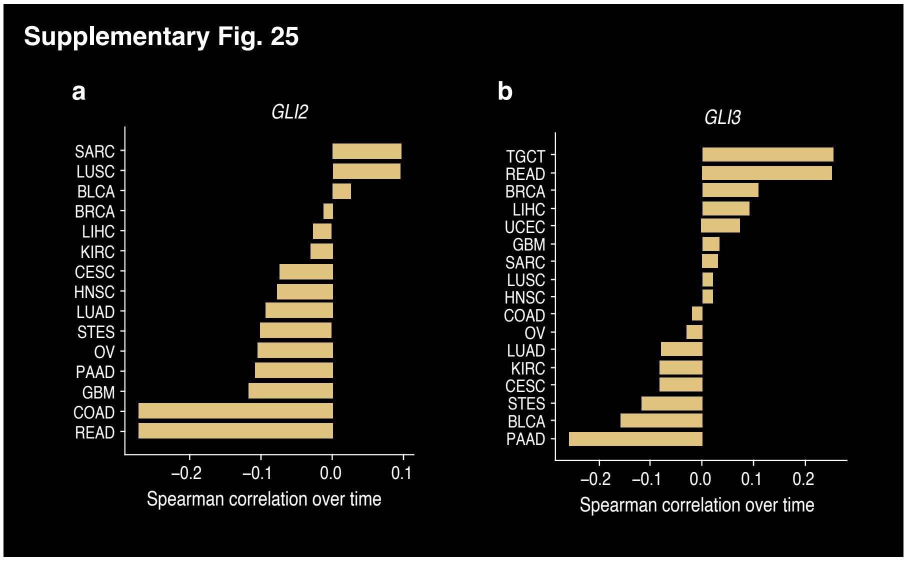

**Supplementary Fig. 25** Normalized mutation rate and survival time for *GLI* mutations. **a,b,** Bar plot of the Spearman correlation of survival time and a patient's mutation normalized by their total mutation load for the *GLI2* (**a**) and *GLI3* (**b**) genes, which are transcription factors in the Hedgehog signaling pathway. We show the Spearman correlation for each cancer type where at least two patients in the time series have a mutation in the gene.

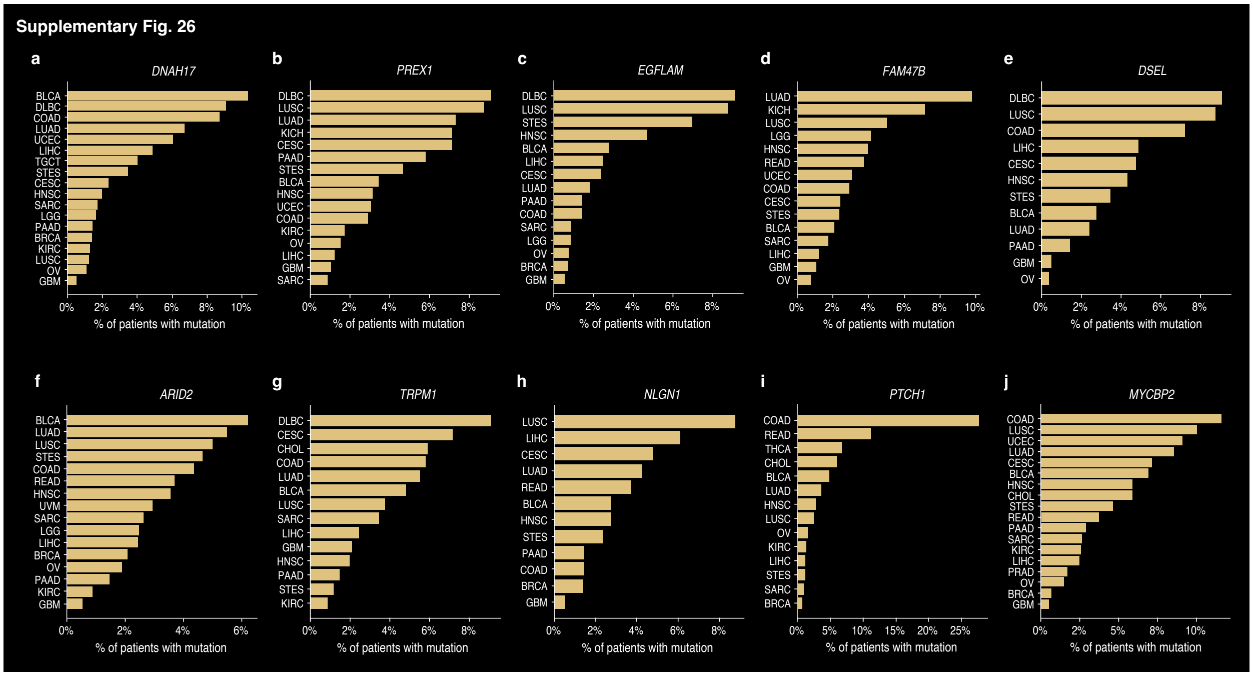

**Supplementary Fig. 26** Mutation frequency across cancer types for Sagittarius's predicted early-stage sarcoma mutations. **a-j,** Bar plot of the percentage of patients in each cancer type with a mutation in the *DNAH17* (**a**), *PREX1* (**b**), *EGFLAM* (**c**), *FAM47B* (**d**), *DSEL* (**e**), *ARID2* (**f**), *TRPM1* (**g**), *NLGN1* (**h**), *PTCH1* (**i**), and *MYCBP2* (**j**) genes. We show a percentage for each cancer type where at least one patient in the time series has a mutation in the gene.

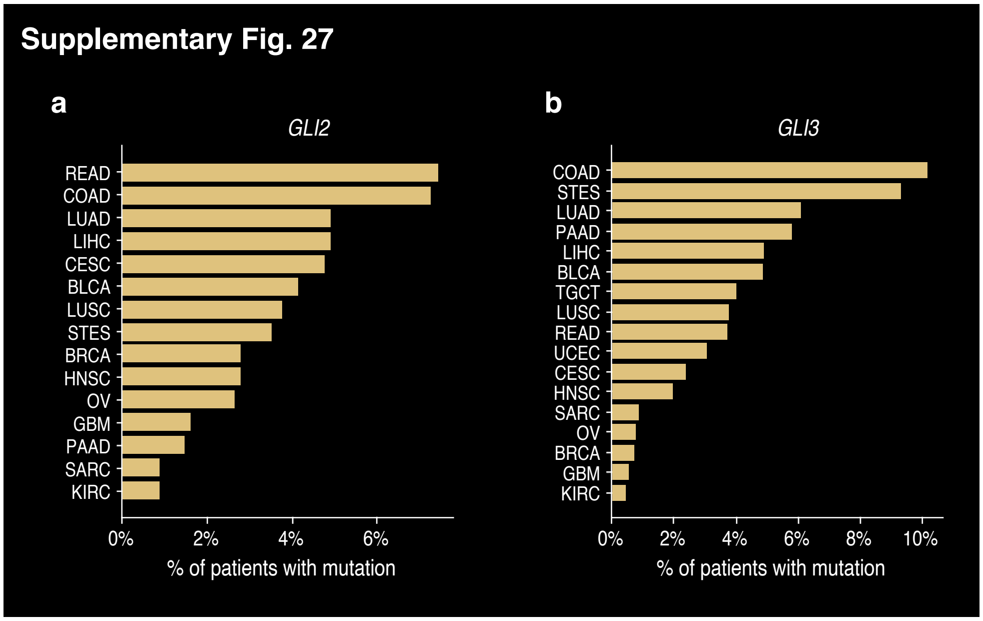

**Supplementary Fig. 27** Mutation frequency across cancer types for *GLI* mutations. **a,b,** Bar plot of the percentage of patients in each cancer type with a mutation in the *GLI2* (**a**) and *GLI3* (**b**) genes, which are transcription factors in the Hedgehog signaling pathway. We show a percentage for each cancer type where at least one patient in the time series has a mutation in the gene.

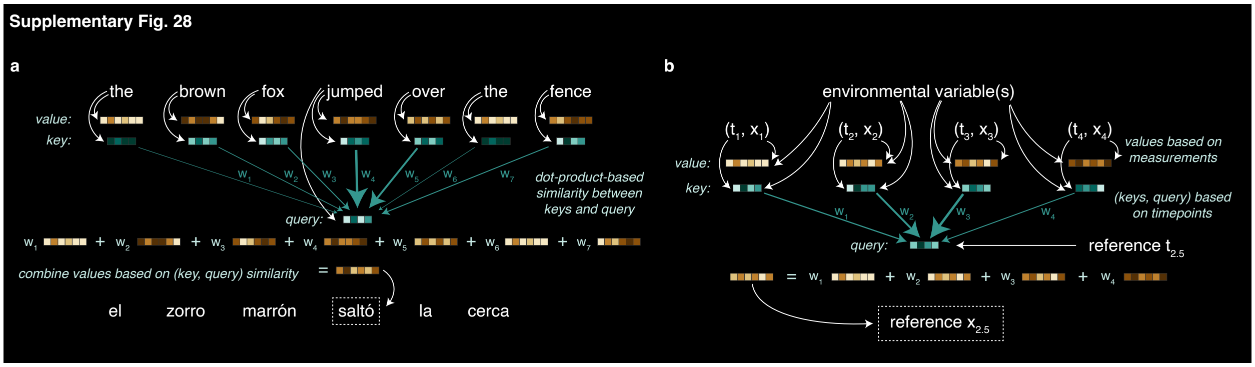

**Supplementary Fig. 28** Illustration of attention mechanisms in different transformer architectures. **a,** Conventional natural language processing transformer with self-attention for a toy machine translation task. The keys, queries, and values of the transformer's attention computation are all based on the words in the input. **b,** Conditional continuous transformer used in Sagittarius, where keys and queries are computed from the timepoints (and conditioned on the environmental variables) and the value is computed from the measurement and environmental variables. For simplicity, we show the encoder mapping to the reference space only; the decoder would proceed analogously.
